# Hematopoietic stem cells undergo bidirectional fate transitions *in vivo*

**DOI:** 10.1101/2025.02.23.639689

**Authors:** Tsuyoshi Fukushima, Trine Ahn Kristiansen, Lai Ping Wong, Yosuke Tanaka, Masaki Yagi, Yu-Hsuan Chang, Takaharu Kimura, Samuel Keyes, Michael Mazzola, Ting Zhao, Lingli He, Susumu Goyama, Ryo Yamamoto, Konrad Hochedlinger, Satoshi Yamazaki, Ruslan I. Sadreyev, David T Scadden

## Abstract

Differentiation of haematopoietic stem and progenitor cells (HSPCs) is widely considered unidirectional *in vivo*. Here, we developed clonal phylogenetic tracing (CP-tracer) by sequential genetic barcoding, which enabled high-resolution analysis of 181,695 subclones derived from 847 individually labelled HSPCs. This approach uncovered bidirectional fate transitions between myeloid-biased (My-) and lineage-balanced haematopoietic stem cells (HSCs). Individual HSC clones, distinguished by temporally unique serial barcodes, exhibited durable lineage bidirectionality, with dynamics favouring progressive accumulation of My-HSCs over time. CRISPR–Cas9 screening identified the homeobox gene *Hhex* as a suppressor of myeloid differentiation that enables pivoting toward balanced output. Hhex function is age-dependent, with its expression declining in aged HSCs. Together, these findings demonstrate unexpected plasticity in HSC differentiation, modulated in part by Hhex-mediated repression, that shifts over time to drive the myeloid bias characteristic of ageing.

## Main

Hematopoietic stem cells (HSCs) are a self-renewing, multipotent population comprised of subsets with differences in the balance of production between myeloid and lymphoid lineages^1–3^. HSCs capable of durable hematopoietic repopulation, corresponding to those with high stemness, are characterized by a bias toward self-renewal over differentiation^4–7^ and typically exhibit myeloid-biased output^3,8–11^. We and others developed systems to clonally trace HSCs^8,11–16^. These defined functional subsets such as “balanced-HSCs” producing myeloid and lymphoid cells, “myeloid biased-HSCs (My-HSCs)” predominantly producing myeloid cells and “lymphoid-biased HSCs (Ly-HSCs)”predominantly producing lymphoid cells. These HSC clones with distinct endogenous outcomes or responses to exogenous stimuli^17,18^ are scripted by epigenetic features^16^. Among HSC features thought to be constrained is directionality of differentiation. For example, HSCs expressing *Vwf* are largely myeloid biased, possess high self-renewal potential and unidirectionaly give rise to *Vwf*□HSCs^8^, but *Vwf□* HSCs did not appear capable of generating *Vwf*^+^ HSCs^9^. Multiple studies support differentiation proceeding down a gradient from the hierarchical apex of platelet biased My-HSC^4,9,12^ to myeloid-HSC and unidirectionally giving rise to balanced-HSC^9,12^. Yamamoto et al. used large scale single cell transplantion to demonstrate that My-HSCs acquire a balanced hematopoietic output upon secondary transplantation^11^. Together, these approaches share the limitation that HSC fate transitions are either inferred indirectly from population markers or the collective output of heterogeneous descendants derived from single HSC. Direct detection of fate transitions between My-HSCs and balanced-HSCs, particularly bidirectional transitions between these differentiation states requires a different approach.

Phylogenetic tracing allows the tracking of individual cells and their descendants by introducing sequential genetic tags over time. Some phylogenetic barcoding methods use a single genomic locus, where the limited capacity to generate diverse barcode variants restricts the ability to resolve complex lineage trajectories^19^. Distributing barcodes across multiple genomic loci has also been employed^20–22^, though this requires integrating barcode information at the single-cell level across genomic loci, limiting throughput and constraining quantification of lineage output within each individual subclone. We created a clonal phylogenic tracer, “CP-tracer,” by fusing a stable barcode with a mutable ‘scratchpad’ to achieve high barcode diversity within a single locus. We postulated that this would enable time dependent sequential labelling, effectively resolving lineage bias within each subclone across complex trajectories. High-resolution analysis of 181,695 subclones derived from 847 individual HSCs revealed previously undocumented bidirectional differentiation fates of HSCs. Furthermore, conducting CRISPR based screening on labeled cells allowed us to define molecular determinants of lineage fate decisions. Among these, the homeodomain transcription factor, *Hhex*, was found to regulate the bifurcation of HSC lineage fate into myeloid- or lymphoid-biased subsets by repressing myeloid competence; a function that declines with age enabling progressive myeloid bias.

## Results

### Time-course tracing by random DNA barcoding identifies novel progenitor types

DNA barcoding inserts a unique sequence into the genome for each cell that is inherited by daughter cells enabling lineage tracing from a single parental cell^6,23^. However, non-nucleated Plt and RBC lineage tracing has been difficult to accomplish. We modified and applied previously described methods^24^ for recovering expressed barcode information from platelet (Plt) and red blood cell (RBC) mRNA (Supplementary Fig.1a-d and Methods) to achieve a more complete definition of clonal differentiation capability. Very low expression of other lineage-specific genes indicated low mRNA contamination (Supplementary Fig.1e,f). Barcode-transduced immunophenotypic HSCs (CD150^+^CD48^-^ckit^+^Sca1^+^Lin^-^) (Supplementary Fig.1g) were transplanted into irradiated mice and the barcodes analyzed sequentially in blood and bone marrow (BM) cells (Fig. 1a and Supplementary Fig.1h-1k). The recovered barcodes were classified by unsupervised k-mean clustering and 6 cell types were identified based on the cell types they produced (Fig. 1b and Supplementary Fig.1l). The functionally classified subgroups are listed on the vertical-axis in the heatmaps throughout the Figures and are based on the presence of the barcode in the immunophenotypically defined hematopoietic cell types indicated on the horizontal-axis; time post-transplant is also indicated on the horizontal-axis. Functional cell grouping of cells bearing the identifying barcodes (represented by a row in the heatmap) included: 1. My-HSCs producing myeloid lineage cells with suppressed or delayed lymphoid output while maintaining immunophenotypic HSCs (CD150^+^CD48^-^Flt3^-^ckit^+^Sca1^+^Lin^-^ for barcode recovery) 20 weeks after transplantation. Because 94.7% of cells in this cluster (216 of 228 clones) satisfied the previously reported criterion for myeloid bias (Myeloid/(Myeloid+B+T) > 2/3)^7^, we defined this cluster as My-HSCs (Fig. 1c). 2. balanced-HSCs producing all blood cell lineages while maintaining immunophenotypic HSCs. 3. Myeloid restricted progenitors (MyRPs) producing myeloid cells over >8 weeks after transplantation without maintaining immunophenotypic HSCs. 4. Lymphoid biased multipotent progenitors (Ly-MPPs) producing all blood cell lineages but without evidence for maintaining immunophenotypic HSCs and decreased myeloid production over time in the primary transplant (Fig. 1b-1d and Supplementary Fig.1l). Cells with similar functional properties as have previously been referred to as HSCs^11^; however, because these cells did not maintain immunophenotypic HSCs in our assay, we refer to them here as Ly-MPPs. These subsets correlate functionally with the 4 cell types reported by others in single cell transplantation studies^8,11,15^. Also observed were two functional groups not previously described. 5. B-biased multipotent progenitors (B-MPPs) that produced myeloid and B cells but not T cells. They did not maintain immunophenotypic HSCs and their myeloid production decreased within the primary transplant (Fig. 1b, c and Supplementary Fig.1l). This pattern was similar to Ly-MPPs but differed in T cell potential. 6. Transient myeloid progenitors (TMPs) that produced Plt, RBC and/or GM only through 4 weeks (Fig. 1b, 1c and Supplementary Fig.1l). This pattern differed from MyRPs in the duration of myeloid production. Only My-HSCs and balanced-HSCs could maintain immunophenotypic HSCs 20 weeks after transplantation; therefore, only these two fractions were regarded as validated stem cells.

**Figure 1.**
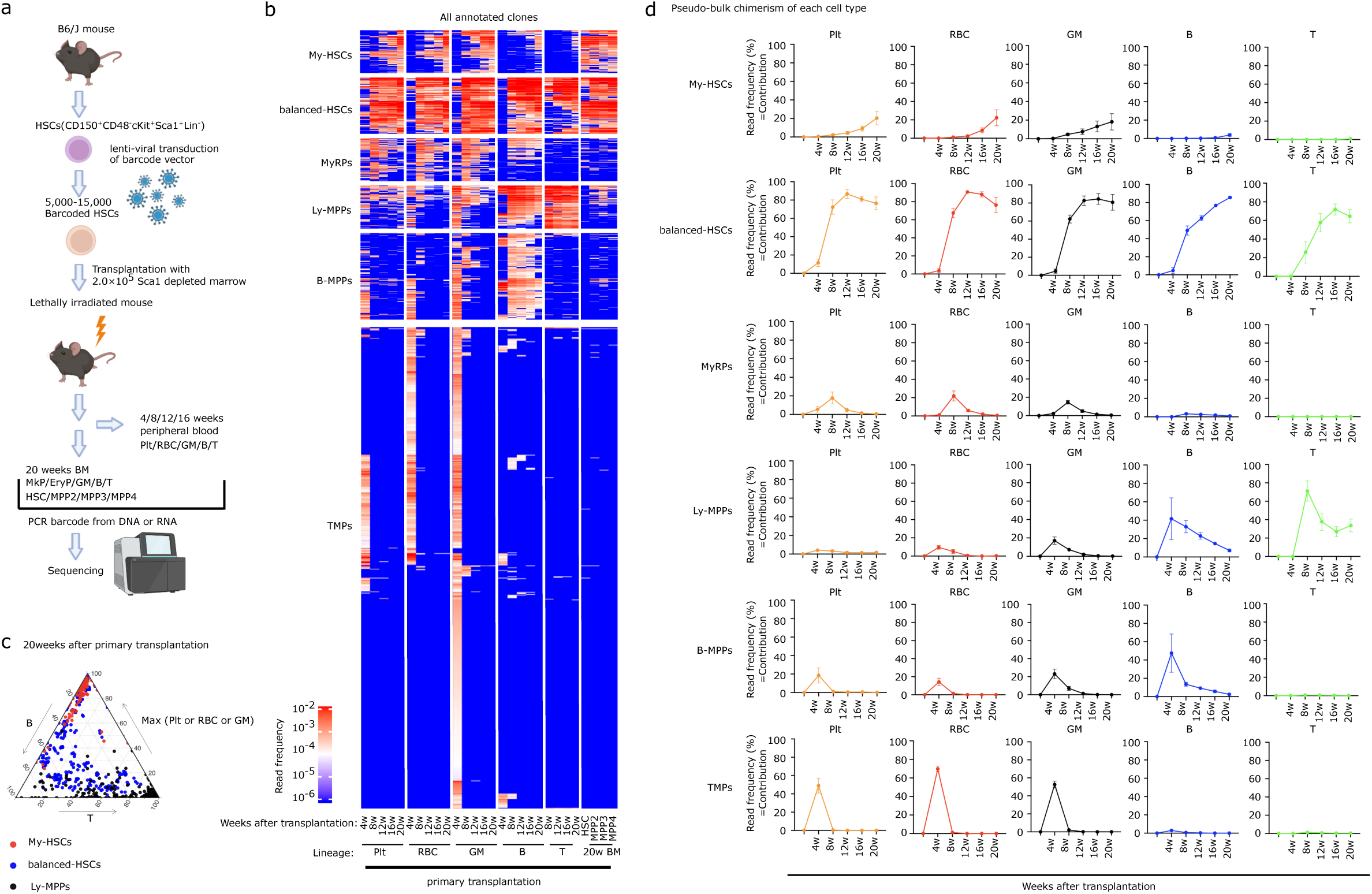
Time-course tracing by random DNA barcoding identifies novel progenitor types

The B-MPP and TMP functional phenotype was further verified by analysis of single-cell transplantation at 24 weeks post-primary transplantation and 20 weeks post-secondary transplantation (Supplementary Fig.1m-n). The production of specific cell types for limited duration was validated for both the B-MPP and TMP. Further support for these classifications was evident in re-analyzing data from others (Supplementary Fig.1o)^25^. Therefore, time-course tracing by random DNA barcoding identified six functional hematopoietic subsets.

We next sought to determine if a similar spectrum of functional cell subsets could be identified in humans. Reanalyzing the long-term lineage output data on lentiviral insertion site marked CD34+ cells from Calabria et al., My-HSC, balanced-HSC, Ly-MPP, B-MPP and TMPs phenotypic profiles were observed^26^ (Supplementary Fig.2a-c). Collectively these data validate the hierarchical presence of phenotypically distinctive subsets of primitive hematopoietic cells in mouse and human including the previously undescribed B-MPP and TMP.

### My-HSCs favor self-renewal and can transition to balanced-HSCs

Multiple studies support differentiation proceeding down a gradient from the hierarchical apex of My-HSCs and giving rise to balanced-HSCs^3,8–11^. Leveraging large-scale time-course tracing by random DNA barcoding, we evaluated the dynamics of My-HSCs and balanced-HSCs. Here, self-renewal was operationally defined as the capacity to repopulate immunophenotypic HSCs at 20 weeks after transplantation, rather than strict maintenance of the same functional HSC subset; clones meeting this criterion were termed “HSC-repopulating clones”. Consistent with a previous report^8^, HSC-repopulating clones could be classified as Plt only (P-HSCs), Plt/RBC/GM (PRG-HSCs), Plt/RBC/GM/B (PRGB-HSCs) and all blood cell lineages (balanced-HSCs) by their lineage output (Fig. 2a, b). P-HSCs, PRG-HSCs and PRGB-HSCs bundled together consitute the My-HSCs group. To account for differences in differentiation kinetics between myeloid and lymphoid lineages, we assessed lineage bias at the time point when myeloid engraftment reached its peak. Controlling for time-dependent effects, 94.3 % (297/315) of My-HSCs consistently showed a higher myeloid contribution (Myeloid/(B+T) > 2)^7^ (Supplementary Fig.3a-g), indicating that their myeloid bias is not attributable to simply delayed lymphoid differentiation, but reflects an intrinsic lineage preference.

**Figure 2.**
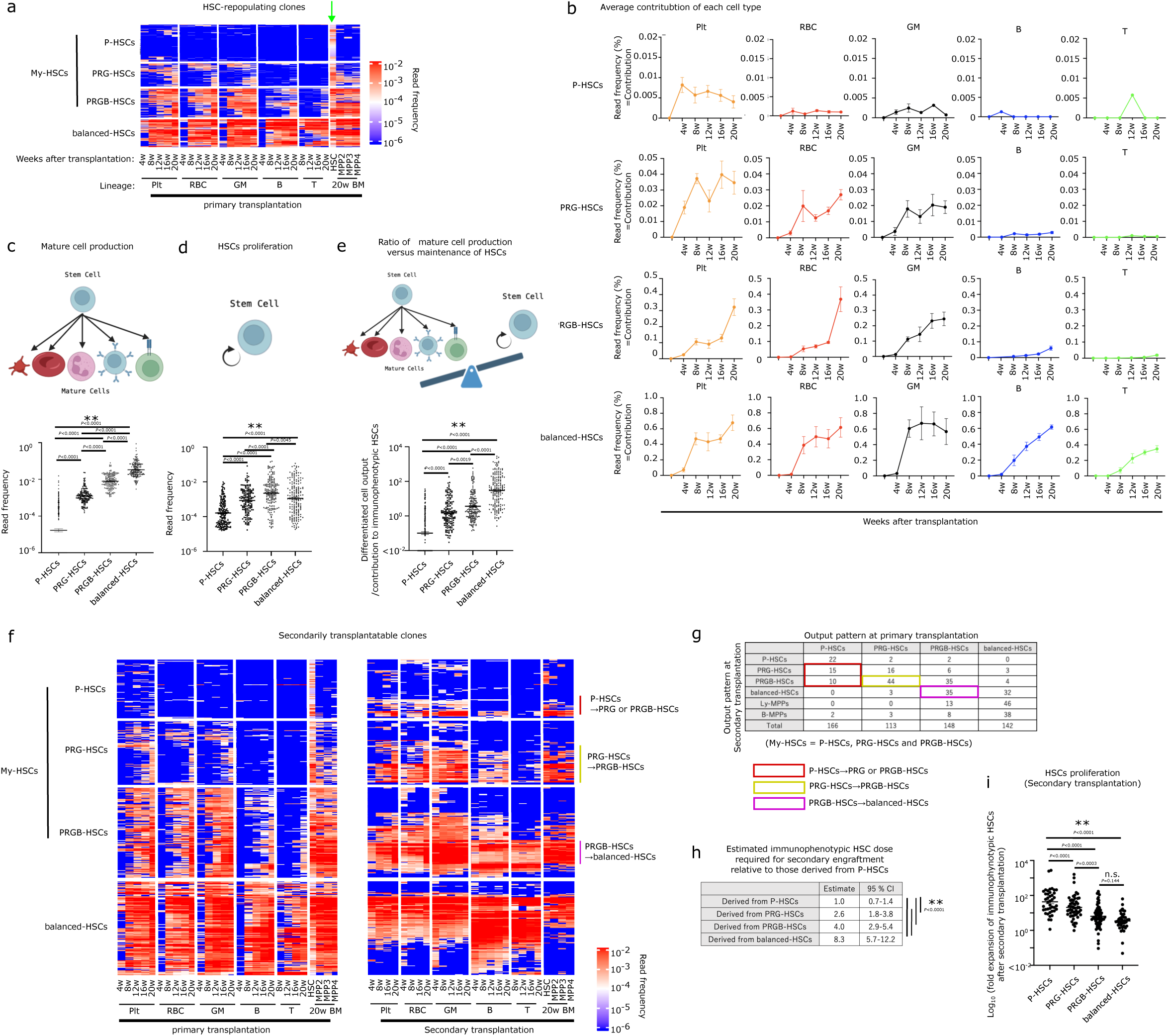
My-HSCs favor self-renewal and can transition to balanced-HSCs

Differentiated blood cell output per clone was significantly higher in balanced-HSCs compared with the My-HSCs (balanced-HSCs > PRGB-HSCs > PRG-HSCs > P-HSCs) (Fig.2c). Since long-term HSC maintenance requires clones to be biased toward self-renewal over differentiation^4–7^(Supplementary Fig.3h), we assessed HSC proliferation as well (Fig.2d) and then compared the relative production of mature cells versus HSC. The ratio of mature cells to immunophenotypic HSCs was markedly higher in balanced-HSCs compared with the My-HSC subsets (balanced-HSCs > PRGB-HSCs > PRG-HSCs > P-HSCs)(Fig.2e). Based on the criterion of relative self-renewal versus differentiation for ‘stemness’^6^, My-HSCs have greater stemness than balanced-HSCs.

The cellular output of immunophenotypic HSC clones on primary and secondary transplantation was assessed. Most My-HSCs that maintained the HSC immunophenotype after secondary transplantation (40/41 P-HSCs, 52/52 PRG-HSCs, and 62/62 PEMB-HSCs) contributed to mature cells and/or MPPs, indicating preserved downstream differentiation capacity. A general loss of primitiveness was observed, however. That is, My-HSC subsets retained their myeloid bias, but produced a broader range of myeloid cells upon secondary transplantation: P-HSCs yielded PRG and PRGB-HSC; PRG-HSCs yielded PRGB-HSC (Fig.2f,g and Supplementary Fig.3i, j).

The ‘stemness’ of My-HSC was further evident in secondary transplant based on the estimated cell dose required for secondary engraftment. The number of immunophenotypic HSC needed for secondary engraftment from each of the My-HSC subsets progressively increased from P-HSCs to PRG-HSC to PRGB-HSC and was significantly higher for balanced-HSC (Fig.2h). Furthermore, My-HSCs derived immunophenotypic HSCs exhibited greater expansion upon secondary transplantation compared with those derived from balanced-HSCs (Fig.2i). These data provide additional evidence that My-HSCs have greater stem cell features and can differentiate to the less self-renewing balanced-HSCs.

### Development of clonal phylogenic tracer: “CP-tracer”

We then explored the capacity to transition between functional subsets by tracing the lineage fate of distinct offspring derived from an individual parental cell. Using CRISPR/Cas9^21,27–29^ with an array of CRISPR targets, called “*Scratchpad*,” ^27^ that accumulate mutations over time, we could track distinct progeny from a parental cell after progressive cell divisions (Fig.3a). Scratchpad allows for multiple sgRNAs to randomly introduce mutations in their targets by editing at random times^27^. However, a single scratchpad provides ∼10^4^ genetic diversity, insufficient to robustly resolve subclones derived from hundreds to thousands of parental cells^19^. Using multiple loci increases barcode diversity, however it requires single-cell analysis for integrating the information, limiting the number of cells for barcode recovery (Supplementary Fig.4a-c) and constraining the quantification of lineage output for individual subclones (Fig.3b and Supplementary Fig.3d)^20,21,30^. To overcome these limitations, we combined random DNA barcodes and scratchpad in a single locus to identify parental cells and their progeny respectively. This allowed for quantification of lineage output for vast numbers of individual subclones without requiring single cell analysis (Fig. 3a-b and Supplementary Fig.4d). We modified the original scratchpad^27^ using 3 sgRNAs with 2 target sequences for each of the 3 sgRNAs in different orientations (Supplementary Fig.4e-z and Methods). This CP-tracer was transduced into immunophenotypic HSCs from Cas9-EGFP mice and transplanted into lethally irradiated recipients, analyzing blood and BM (more than 4×10^7^ hematopoietic cells; F Supplementary Fig.4n-s). The random DNA barcodes of CP tracer could identify clones such as My-HSCs, balanced-HSCs, Ly-MPPs, and B-MPPs (Fig.3d,e). The scratchpad codes indicated the fate of descendant subclones. Subclones were defined by the unique allele (combination of random DNA barcodes and scratchpad variants) in each replicate, with the clone of origin identifiable by the random DNA barcode component (Fig.3b,c). Of a total of 200,204 alleles, 199,337 (99.57%) were unique across 7 replicates, indicating that the allele diversity generated by the CP tracer is sufficiently large relative to the total number of labeled cells. When considering scratchpad variants, 90.8 % (181,695 of 200,204) of alleles carried unique scratchpad variants across all clones. The 9.2% of scratchpad variants that showed overlap between clones were shared across an average of 3.08 clones among the 847 clones analyzed, suggesting that even for scratchpad variants showing overlap, the probability that distinct cells independently generated the same variant was approximately 0.36% (3.08/847). (Supplementary Fig.4t); subsequent analyses focused primarily on alleles with unique scratchpad variants. With this filter, the estimated probability that distinct cells within a clone independently generate the same scratchpad variant is reduced to less than 0.11% (1/847). Most of the scratchpads were edited in 3-5 out of 6 sites generally without large deletions (Supplementary Fig.4u-w). Editing percentage at each site 20 weeks after transplantation was very similar to that at 14 days of culture (S Supplementary Fig.4i,x), and was stable over time (Supplementary Fig.4y). The number of scratchpad variants per clone (i.e., subclones per barcode clone) was highest in balanced-HSCs, with a mean of 621.6 ± 52.2 subclones per clone (Supplementary Fig.4z). Furthermore, CP-tracer labeled around 600 offspring from a single HSC enabling the tracing of early fate segregation from repopulating HSCs.

**Figure 3.**
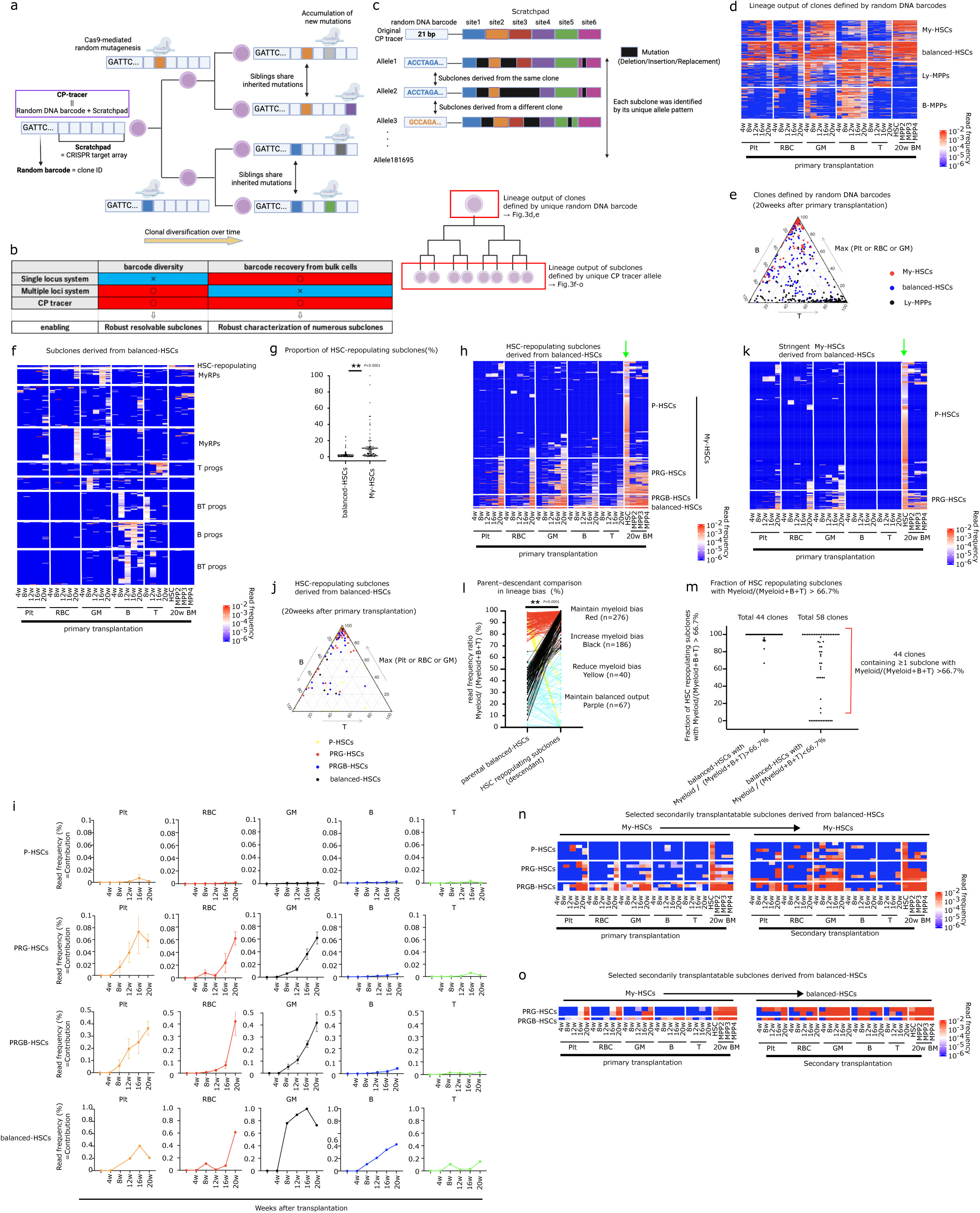
Balanced-HSCs can transition into My-HSCs

### Balanced-HSCs can transition into My-HSCs

Immunophenotypic HSCs transduced with CP tracer were transplanted and the subclones arising were analyzed (Fig.3a-f and Supplementary Fig.4n-r,5a) at 20 weeks. Only 2.33 ± 0.37 % of the subclones derived from balanced-HSCs were HSC-repopulating subclones compared with 10.71 ± 1.70 % of My-HSCs (4.6 fold higher; *P* = 0.0001; Fig.3g and Supplementary Fig.5a). Interestingly, 98.7 ± 0.63% of the HSC-repopulating subclones (989 of 995) derived from balanced-HSCs had an output comparable to My-HSCs in which lymphoid output was suppressed or delayed (Fig.3h-j). These data are consistent with balanced-HSCs transitioning to My-HSCs despite the higher stemness^6^ of My-HSCs. We defined My-HSCs through k-means clustering while some previous studies employed a more stringent definition^8,11^ of clones with no lymphoid output. Even applying the mores stringent definition, balanced-HSC were observed to gave rise to My-HSCs (Fig.3k). Our sequencing depth and sampling scale (∼1.25 million cells per lineage time point for nucleated cells) enables detection of subclonal contributions as low as 0.001–0.01% (Methods). Given that PRG-HSCs and PRGB-HSCs showed myeloid chimerism of approximately 0.06% and 0.4%, respectively (Fig.3i), we reasoned that lack of detected lymphoid output at ≤0.01% likely reflects true lineage bias. In P-HSCs, lineage outputs often fell below the detection threshold for individual subclones, but among 658 evaluable P-HSCs, myeloid-only detection (270 P-HSCs) was significantly more frequent than lymphoid-only detection (25 P-HSCs) (McNemar’s test, p < 0.0001). Chimerism levels were comparable between myeloid and lymphoid lineages when detected (Supplementary Fig.5b), arguing against differential detection sensitivity as the cause of the observed differences.

To directly assess lineage bias dynamics during clonal evolution, we analyzed parent–descendant relationships by quantifying Myeloid/(Myeloid+B+T) ratio for each parent clone and its derived subclone, and plotted the changes in this value across parent–descendant pairs (Fig. 3l). Among HSC-repopulating subclones derived from balanced HSCs with Myeloid/(Myeloid+B+T) < 66.7%, 186 of 253 subclones increased their Myeloid/(Myeloid+B+T) to > 66.7%, with a average increase of 55.74 ± 0.96% (Fig.3l black lines). In addition, 44/58 parent balanced-HSCs with Myeloid/(Myeloid+B+T) < 66.7% produced more than 1 subclone with Myeloid/(Myeloid+B+T) > 66.7% (Fig.3m). Among subclones derived from balanced-HSCs with Myeloid/(Myeloid+B+T) > 66.7%, 276/316 exhibited a further increase of myeloid bias with a average increase of 8.60 ± 0.36% (Fig.3l red lines).These results demonstrate progressive enhancement of myeloid bias during clonal evolution. In addition, My-HSCs derived from balanced-HSCs maintained a My-HSC fate after secondary transplantation (Fig.3n), indicating a durable fate transition and validating that these cells represent bona fide My-HSCs. Futher, My-HSCs derived from balanced HSCs possess nearly equivalent bone marrow reconstitution capacity as My-HSCs derived from My-HSCs (Supplementary Fig.5c). We analysed DARLIN data generated by others^20^ and identified 6 balanced-HSCs with detectable progeny; among these, two clones were found to give rise to My-HSCs (P-HSCs) in their descendant cells (Supplementary Fig.5d,e). Furthermore, studies of human clonal hematopoiesis conducted by others^31^, indicated the emergence of HSCs lacking T-lineage output as descendants of balanced-HSCs and lacking B-lineage output as descendants of PRGB-HSCs. These data suggest a reversion in fate under non-transplant conditions^31^ (Supplementary Fig.5d,e), showing that this is not a phenomenon unique to the transplantation setting. Random DNA barcodes alone also revealed transition from balanced-HSCs to My-HSCs as observed in 3 of 143 secondary transplantable balanced-HSCs (Supplementary Fig.5f). These further support our conclusion that balanced-HSCs transition to My-HSCs.

Combining the data above with evidence for My-HSCs transitioning to functional balanced-HSCs (Fig.2f,g) is consistent with bidirectional fate transitions. Notably, a small number of My-HSCs derived from balanced-HSCs reverted back to balanced-HSCs upon secondary transplantation (Fig.3o). Therefore, bidirectional fate plasticity between balanced-HSCs and My-HSCs occurs at a single cell level.

Because My-HSCs are defined by insufficient lymphoid output during primary transplantation, descendant subclones exhibiting high lymphoid output would not be expected within the primary transplant setting. Accordingly, none of the 412 HSC-repopulating subclones derived from 106 My-HSCs were classified as balanced-HSCs at primary transplantation (Supplementary Fig.5g-i). To directly assess lineage bias dynamics during clonal evolution in My-HSCs, we analyzed parent–descendant relationships by quantifying Myeloid/(Myeloid+B+T) ratio for each parent clone and its derived subclone. Among HSC-repopulating subclones derived from My-HSCs, 160/197 subclones exhibited a further increase of Myeloid/(Myeloid+B+T) ratio, with an average increase of 6.06 ± 0.79%. (Supplementary Fig.5j red lines). Nevertheless, 37/197 subclones showed a reduction in Myeloid/(Myeloid+B+T), with an average decrease of 18.54 ± 5.05% (Supplementary Fig.5j black lines). In rare cases, 3 of 73 parent My-HSCs with Myeloid/(Myeloid+B+T) > 66.7% produced at least one descendant subclone with Myeloid/(Myeloid+B+T) < 66.7% (Supplementary Fig.5k). Consistent with the results obtained using random DNA barcoding (Fig.2g), only a fraction of HSC-repopulating subclones (2 of 25) transitioned to balanced-HSCs even after secondary transplantation (Supplementary Fig.5l,m). In contrast, the majority of balanced-HSCs transitioned to My-HSCs (Fig.3h,l); therefore, the bidirectional transition between balanced-HSCs and My-HSCs is biased in the direction of balanced-HSCs transitioning to My-HSCs. Myeloid bias predominates.

In sum and consistent with previous reports^3,8–10^, My-HSCs possess greater stemness^6^ than balanced-HSCs and can transition to balanced-HSCs (Fig.2a-i) but unexpectedly, CP tracer showed that balanced-HSCs can reverse their trajectory and transition to My-HSCs (Fig.3h-o) a process that results in progressive accumulation of My-HSCs.

### Balanced-HSCs contribute to mature blood cell lineages through My-HSCs or **Ly-MPPs**

To investigate how HSCs give rise to diverse lineages, phylogenetic trees for each clone were reconstructed from scratchpad editing patterns (Fig.4a, details are provided in the Methods section). A branch point in the phylogenetic tree is referred to as a node, representing a common ancestral progenitor from which descendant branches originate. A total of 181,695 unique alleles were generated from combinations of 25,098 distinct mutations. The average occurrence rate of each mutation was 0.022 ± 0.19 % across all alleles, indicating that most mutations shared among different subclones (alleles) within the same clone are likely inherited from a common ancestor cell that harbored these mutations, rather than arising independently through convergent mutations (Supplementary Fig.6a). Bootstrap analysis showed that the branches were reproduced in 81.86 ± 0.33% (Supplementary Fig.6b), even when trees were reconstructed from pseudo-datasets generated by randomly omitting or duplicating portions of the scratchpad sequence information, indicating that the tree structures generated by the CP tracer were robust. We also perfomed a simulation in which four daughter cells were generated at each hierarchical level, and one mutation was added to each daughter cell based on the occurrence probability of individual mutations observed experimentally. This process was repeated across four hierarchical levels, resulting in a lineage tree containing a total of 256 descendant cells. The simulation also incorporated inter-site depletion induced by large deletions spanning pre-existing mutations, resulting in the loss of pre-existing mutations^32,33^. In this simulation, inter-site depletion was 3.41 ± 0.49 % and the concordance between the reconstructed and ground-truth simulated lineage trees was 90.52 ± 0.28% in the CP tracer (Supplementary Fig.6c,d), whereas the corresponding values in DARLIN were 4.57 ± 0.30% and 81.49 ± 0.32%, respectively. The high uniqueness of individual mutations (Supplementary Fig. 6a) and suppression of large deletions (Supplementary Fig. 4u,v) in the CP tracer enable accurate phylogeny reconstruction.

**Figure 4.**
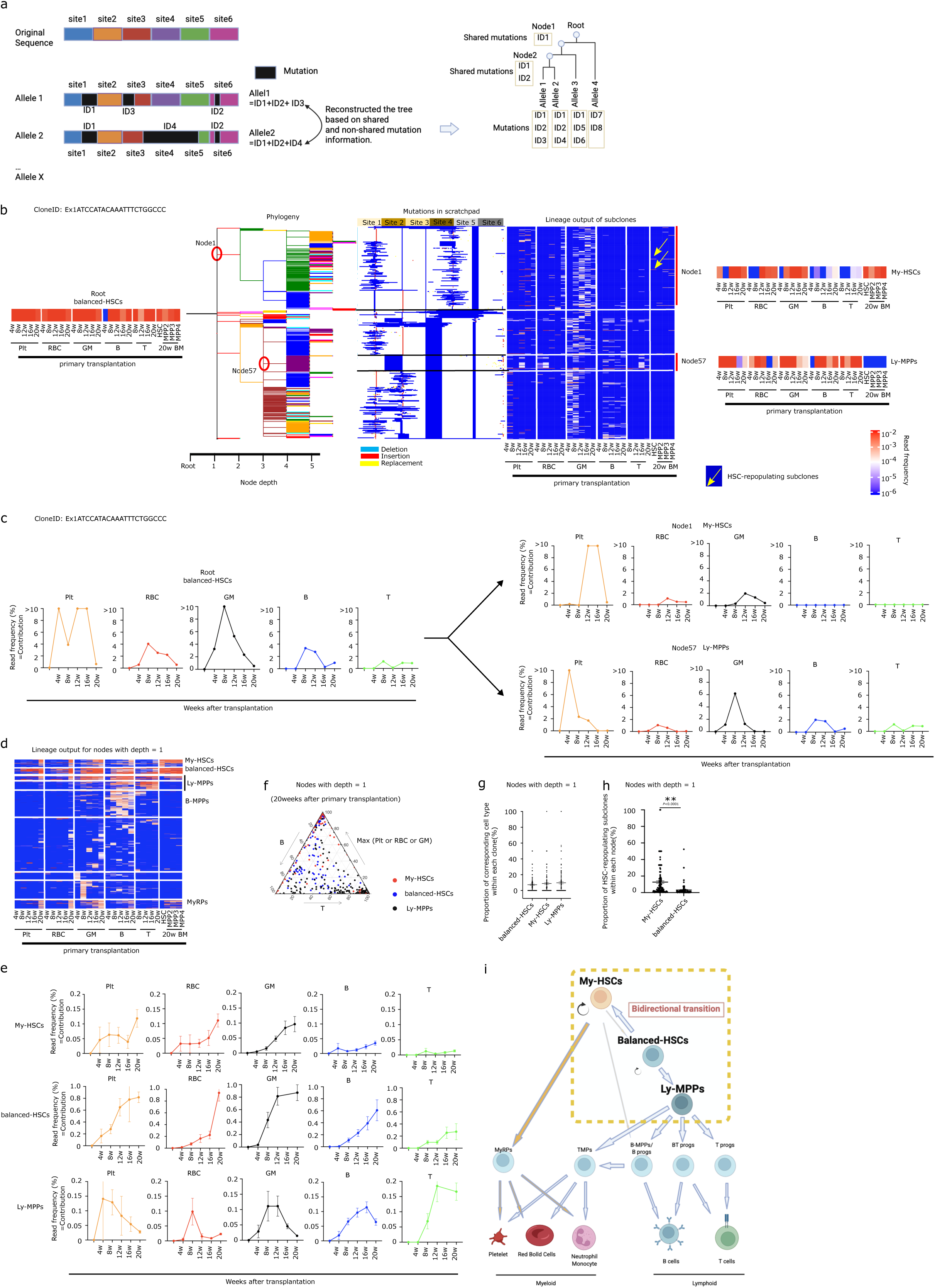
Balanced-HSCs contribute to mature blood cell lineages through My-HSCs or Ly-MPPs

Analyzing the Scratchpad editing pattern, balanced-HSCs branched to balanced-HSCs, My-HSCs, and Ly-MPPs (Fig.4a-c and Supplementary Fig.6e,f). At a branch depth of 1 in the phylogeny derived from balanced-HSCs, maintenance of balanced-HSCs, transition to My-HSCs, and transition to Ly-MPPs were observed at 7.27 ± 0.87%, 8.83 ± 1.26%, and 9.77 ± 1.31%, respectively (Fig.4d-g). Bootstrap analysis confirmed that these branches were repeatedly preserved, indicating that the CP tracer possesses sufficient accuracy to detect the transitions of balanced-HSCs to My-HSCs and Ly-MPPs (Supplementary Fig.6g). Branches that transitioned to My-HSCs have a higher proportion of HSC repopulating subclones among their progeny compared with branches that maintained the balanced-HSC state (Fig.4h), further confirming that My-HSCs possess greater stemness^6^; nevertheless, balanced-HSCs could transition into My-HSCs. Accordingly, both more primitive My-HSCs and more mature Ly-MPPs were confirmed as direct progeny of balanced-HSCs, a finding that was validated by secondary transplantation experiments with random DNA barcodes (Fig.2f,g) and single-cell transplantation (Supplementary Fig.6h). Therefore, the balanced-HSC has two opposing fate options.

In the differentiation pathways from My-HSCs and Ly-MPPs, My-HSCs mainly produced My-HSCs, MyRPs and B-MPPs while Ly-MPPs mainly produced Ly-MPPs, B-MPPs, T progenitors (T progs) and TMPs. Futher, B-MPPs mainly produced B-MPPs and TMPs. (Supplementary Fig.6i-n). BT progenitors (BT progs) observed as descendants of balanced-HSCs were also detected as descendants of Ly-MPPs (Fig.3f, Supplementary Fig.5a,6o). Interestingly, My-HSCs were able to produce B-MPPs/B progenitors (B progs), while that was not observed with Ly-MPPs (Supplementary Fig.6i,j,p), perhaps indicating a parallel path to B cells. Overall, balanced-HSCs contribute to the spectrum of progenitor cells and mature blood cell lineages through two distinct intermediate states, My-HSCs and Ly-MPPs (Fig.4i)

### Combining fate tracing and scRNA-seq identified transcriptional differences between My-HSCs and balanced-HSCs

Combined DNA barcoding and single-cell RNA-seq analysis was conducted to define the gene expression signatures of immunophenotypic HSC according to their functional states *in vivo*. Barcoded immunophenotypic HSCs were transplanted and their lineage output in PB and BM analyzed at 4-20 weeks; scRNA-seq of immunophenotypic HSC from BM was then analyzed 20 weeks after transplantation. The expressed DNA barcodes were aligned by scRNA-seq (Supplementary Fig.1a) and clonal lineage output paired with the transcriptome of the clonal HSC source (Fig.5a and Supplementary Fig.7a-f). Single HSC functional states were annotated according to the differentiation classification shown in Fig.2a. MolO score, defined by the similarity to HSCs with molecular characteristics consistently observed across multiple purification methods^34^ was higher in cells derived from P-HSCs and PRG-HSCs (Fig.5b), indicating that these cells exhibit higher expression of HSC signature genes, consistent with their higher ‘stemness’^6^ defined by functional features (Fig.2e, 2h and 2i). Whereas the cell cycle score,^35^ was low in P-HSCs and PRG-HSCs (Fig.5c) consistent with their limited cell output (Fig.2c,d). Senescence- and cell death–associated molecular signatures in My-HSCs are comparable to, or even lower than, those in balanced HSCs (Supplementary Fig.7g). Functionally, My-HSCs are capable of secondary engraftment with a limited number of cells and show a high expansion rate of the HSC compartment (Fig.2h,i). In addition, previous studies have demonstrated the presence of My-HSCs under physiological conditions without culture or transplantation^13^. Together, these findings indicate that My-HSCs are not a stress-induced state resulting from culture or transplantation. Although the HSC subsets did not form distinct clusters, each subset exhibited a preferential spatial distribution on uniform manifold approximation and projection (UMAP)^36^ (Fig.5d,e). UMAP RNA velocity analysis^37^ demonstrated trajectories of balanced-HSCs transitioning into My-HSCs (Fig.5d and Supplementary Fig.7h), and within My-HSC, PRGB-HSCs transitioning into P/PRG-HSCs (Supplementary Fig.5g-j); consistent with the CP-tracer results.

**Figure 5.**
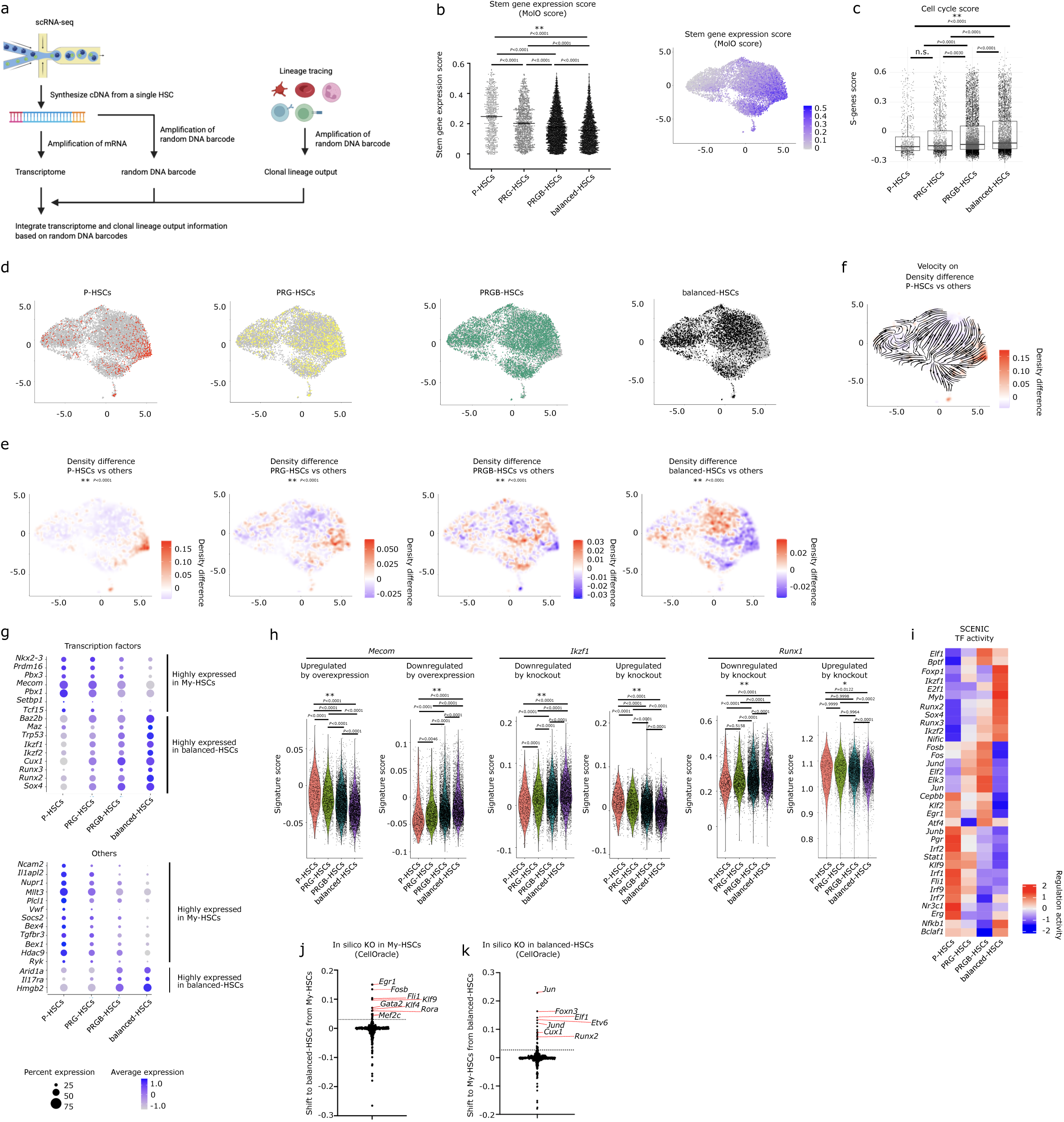
Combining fate tracing and scRNA-seq identified transcriptional differences between My-HSCs and balanced-HSCs.

We next evaluated molecular differences among functionally defined HSC subtypes (Supplementary Fig.7i). Molecules specific for My-HSCs included transcription factors^38^ (*Mecom* and *Tcf15*), known self-renewal regulators (*Mecom*^39^ and *Mllt3*^40^), *Tcf15* that biases toward self-renewal over differentiation^6^, and the My-HSC marker, *Vwf*^9^. Molecules specific for balanced-HSCs included transcription factors (*Runx2*, *Runx3*^41^, and critical lymphoid differentiation regulators, *Ikzf1* and *Ikzf2* (Fig.5g). Signature score analysis showed evidence of high activity of Mecom in My-HSCs (Fig.5h) and combined with gene regulatory network analysis^42,43^ showed evidence of high activity of Runx2/3 and Ikzf1/2 in balanced-HSCs (Fig.5h,i). In silico perturbation analysis identified candidate regulators of the transition between My-HSCs and balanced-HSCs. Several transcription factors identified by CellOracle^44^, including Fli1, Klf9, Fosb, Runx2 and Elf1, also exhibited differential regulatory activity in gene regulatory network analysis (Fig.5h–k). The functional distinctions of HSC subset identified by fate tracing are reflected in confirmatory transcriptional differences.

### Single cell CRISPR screening identifies genes that control HSC heterogeneity

To determine the importance of these transcriptional differences, we combined random DNA barcodes with CRISPR/Cas9 based sgRNA library screening (Fig.6a and Supplementary Fig.8a) focusing on 30 genes selected by their differential expression, gene regulatory network analysis and in silico screening (Fig.6b). The screening vector contained both a random DNA barcode library with approximately 10^6^ variations and a sgRNA library with 100 sgRNAs containing 3 sgRNAs targeting each candidate gene and 10 NT-sgRNAs (Fig.6a and Supplementary Fig.8b-d (details in the Methods section.). Lineage tracing of immunophenotypic HSCs was performed and the effect of gene perturbation on My-HSCs/MyRPs, Ly-MPPs/B-MPPs and TMPs was evaluated (Supplementary Fig.8a and e-g). Of note, HSCs bearing a random DNA barcode and those bearing both barcode and sgRNA performed similarly in vivo (Supplementary Fig.1l and Supplementary Fig.8g). KO of *Junb* increased My-HSCs/MyRPs, KO of *Bex4* and *Junb* decreased Ly-MPPs/B-MPPs, and KO of *Mllt3* and *Hhex* increased TMPs (Fig.6b). *Junb*, *Mllt3* and *Hhex* were reported respectively as a regulator of proliferation^45^, self-renewal^40^ and lymphoid development^46–48^. However, the effects of these genes at the clonal level had not been reported and doing so here validated the method’s utility for functionally defining molecular regulators of individual HSC fate.

**Figure 6.**
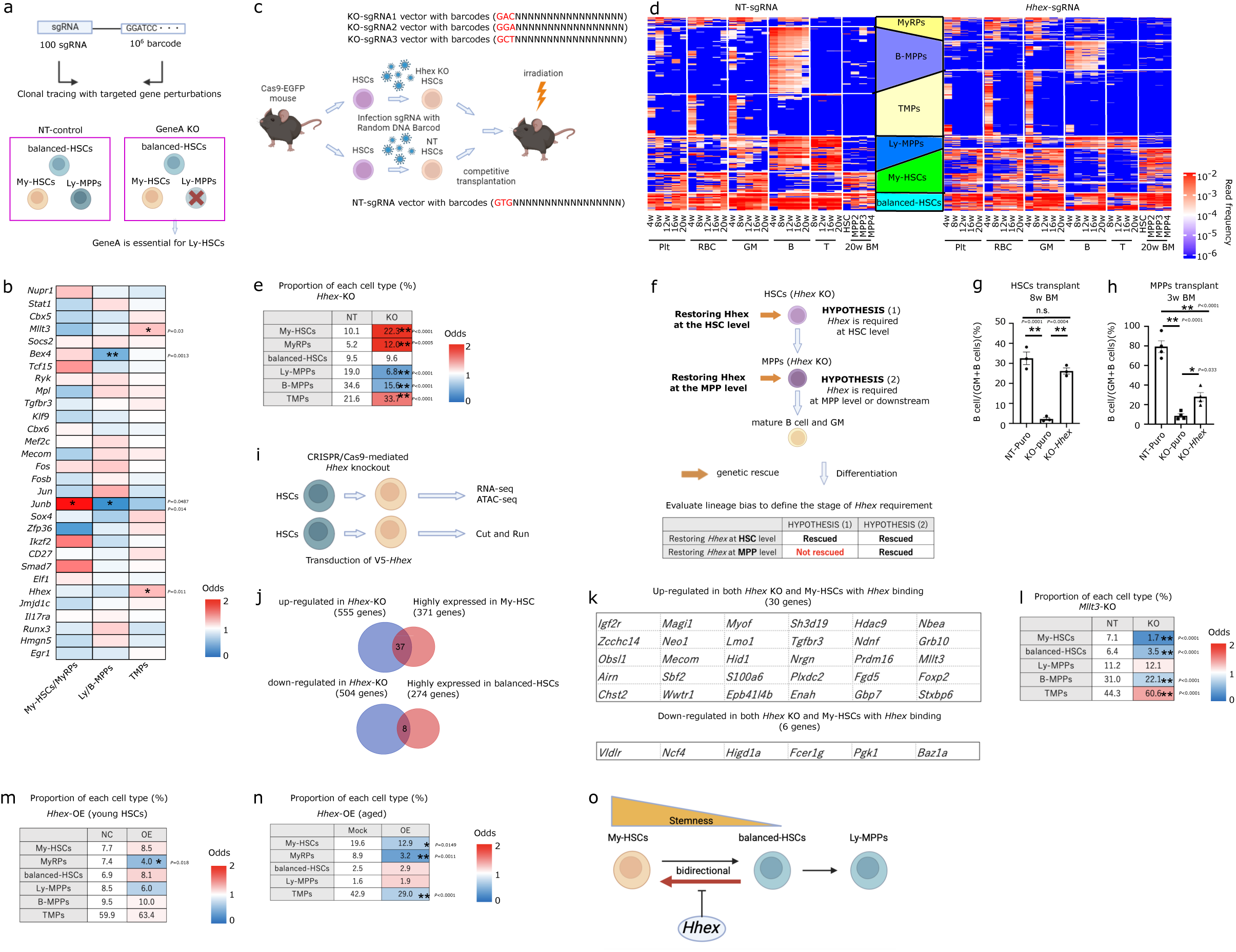
*Hhex* is a suppressor of My-HSC fate

### *Hhex* is a suppressor of My-HSC fate

We further assessed *Hhex,* a homeodomain transcription factor important for embryonic definitive HSC^49^ and for lymphoid differentiation^46–48^. Because the *Hhex* KO has reduced B and T cell progenitors without reduction of either lymphoid primed MPP4 (Flt3^+^ckit^+^Sca1^+^Lin^-^) ^50,51^ or common lymphoid progenitors (CLP: Lin^-^cKit^low^Sca1^low^Flt3^+^Il7R^+^), it has not been thought to participate in early lymphoid fate specification,^46–48^. However, our data indicate multiple paths to MPP4 and CLP that may confound direct linear correlations. That is, both Ly-MPPs and MyRPs produce immunophenotypic MPP4 (Supplementary Fig.9a) and Ly-MPPs, B-MPPs, MyRPs and platelet progenitors were all able to produce immunophenotypic CLP (Supplementary Fig.9a). Therefore, MPP4 and CLP may descend from multiple progenitor populations making their number unreliable as an indicator of lineage source. To account for this, *Hhex* was deleted in clonally marked immunophenotypic HSCs (Fig.6c). *Hhex* KO significantly increased My-HSCs and MyRPs while Ly-MPPs and B-MPPs decreased (Fig.6d,e and Supplementary Fig.9b-e). These data suggest that *Hhex* plays a role early in adult HSC differentiation, likely suppressing My-HSC fate and enabling lymphoid fate competence.

We assessed lineage bias between B-cells and granulocyte/monocytes after transplantation of immunophenotypic HSCs (CD150^+^CD48^-^ckit^+^Sca1^+^Lin^-^) or MPPs (CD48^+^ckit^+^Sca1^+^Lin^-^) derived from cultured Hhex-KO HSCs with and without *Hhex* cDNA add-back (Supplementary Fig.9f-h). Lineage bias was evaluated in the BM, since B cells were not detectable in the PB at 3 weeks after MPP transplantation, even in controls. If the myeloid bias observed in *Hhex*-KO cells primarily results from impaired B-cell differentiation at the MPP stage or downstream^46–48^, lineage bias should be rescued by *Hhex* add-back at both the HSC and MPP levels. In contrast, if the myeloid bias observed in *Hhex*-KO cells originates at the HSC level, lineage bias should be rescued by add-back at the HSC level but not at the MPP level (Fig.6f). Consistent with the latter model, *Hhex* rescued the lineage bias in HSCs (Fig.6g, Supplementary Fig.9i,j). Overexpression in immunophenotypic MPPs was directionally similar but of smaller magnitude (Fig.6h and Supplementary Fig.9i). Therefore, *Hhex* appears to be key for lineage bias at the HSC level. Apoptosis of prepro-B cells by *Hhex* KO was reported by others to be rescued by overexpression of anti-apoptotic factor Bcl2^46^. However, when we overexpressed Bcl2 the *Hhex* KO frequency of My-HSCs/MyRPs was still increased and Ly-MPPs/B-MPPs still decreased (Supplementary Fig.9k-p) arguing against apoptosis causing Ly-MPP reduction.

We performed RNA-seq and ATAC-seq analyses on *Hhex*-deficient HSCs (Fig.6i), complemented by Cut&Run profiling of *Hhex*, to assess the transcriptional regulatory mechanisms governed by *Hhex*. In accordance with previous studies^46,52^, *Cdkn2a* expression was elevated (Supplementary Fig.10a), whereas the expression of other cell-cycle genes remained unchanged (Supplementary Fig.10b). Cut&Run analysis revealed a strong enrichment of *Hhex* binding at promoter regions (Supplementary Fig.10c) and identified AGGAAATTA as the predominant binding motif (Supplementary Fig.10d). The first half of this sequence (AGGAA) corresponds to an ETS motif, whereas the second half (AATTA) matches a homeobox motif. Consistent with this, regions exhibiting altered chromatin accessibility upon *Hhex* KO were significantly enriched for both ETS and homeobox motifs (Supplementary Fig.10e,f), suggesting that the homeobox transcription factor *Hhex* may cooperatively mark the same regulatory regions as ETS family factors. Assessment of accessibility changes further showed that *Hhex* KO resulted in far more sites with increased accessibility (3,623) than decreased accessibility (1,210) (Supplementary Fig.10e), consistent with a transcriptional repressor role for *Hhex*^53^. In wild type cells, while My-HSCs did not have elevated *Hhex* expression, chromatin regions^41^ containing the *Hhex* binding motif were more accessible by ATAC-Seq in My-HSCs than balanced-HSCs (Supplementary Fig.10g,h), further supporting a suppressive role for Hhex in HSCs. Genes upregulated by *Hhex*-KO showed more Hhex promoter binding than genes downregulated by *Hhex*-KO (77.7 % (428 of 555 genes) vs 69.0 % (348 of 504 genes)) (Supplementary Fig.10i). Analysis of lineage bias among these genes demonstrated that only eight downregulated genes were highly expressed in balanced HSCs, whereas 37 upregulated genes were highly expressed in My-HSCs (Fig.6j,k). These included *Wwtr1*, which is known to increase in aged HSCs and is associated with myeloid bias^54^ (Fig.6k and Supplementary Fig.10j), as well as *Mllt3*, whose knockout preferentially reduces My-HSCs over Ly-MPPs (Fig.6k,l, and Supplementary Fig.10k).Taken together, these findings suggest that *Hhex* maintains HSC balance not by activating Ly-MPP gene programs, but rather by repressing My-HSC–associated genes.

Notably, *Hhex* declines in human bone marrow HSC with age^55^ (Supplementary Fig.10l). When we overexpressed *Hhex* in either young or old bone marrow *in vivo,* myeloid suppression was found as expected. However, the effect was more pronounced on aged hematopoietic cells with more substantial suppression of My-HSC and MyRPs than was observed in young mice (Fig.6m,n). Together, these findings indicate that the primary role of *Hhex* in HSCs is in an age-modulated suppression of myeloid gene expression largely mediated by modification of chromatin accessibility and ETS factor binding.

### *Bex4* enhances My-HSC fate

We also assessed whether our system could identifiy genes with no prior association with hematopoiesis. *Bex4* (Brain-expressed X-linked 4) is a member of the Bex gene family, which was originally characterized for roles in neural development and cell survival^56^. Members of the Bex family have been reported to be pseudosubstrates of E3 ligases^57^ and for Bex4 to inhibit the deacetylase, Sirt2, suggesting its potential to modulate cell state. We found that *Bex4* expression was higher in My-HSCs than balanced-HSCs (Fig.5g). When we overexpressed *Bex4* the proportion of the My-HSCs and MyRPs increased (Fig. 7a-d). On the other hand, *Bex4* KO had little effect (Supplementary Fig.11a-d). These findings suggest that *Bex4* promotes a myeloid-biased fate at the HSC level, although its role may be redundant or compensated under loss-of-function conditions. Therefore, our sequential labeling technique combined with molecular analysis and CRISPR screening identified both novel cell lineage dynamics and molecular determinants of those events.

**Figure 7.**
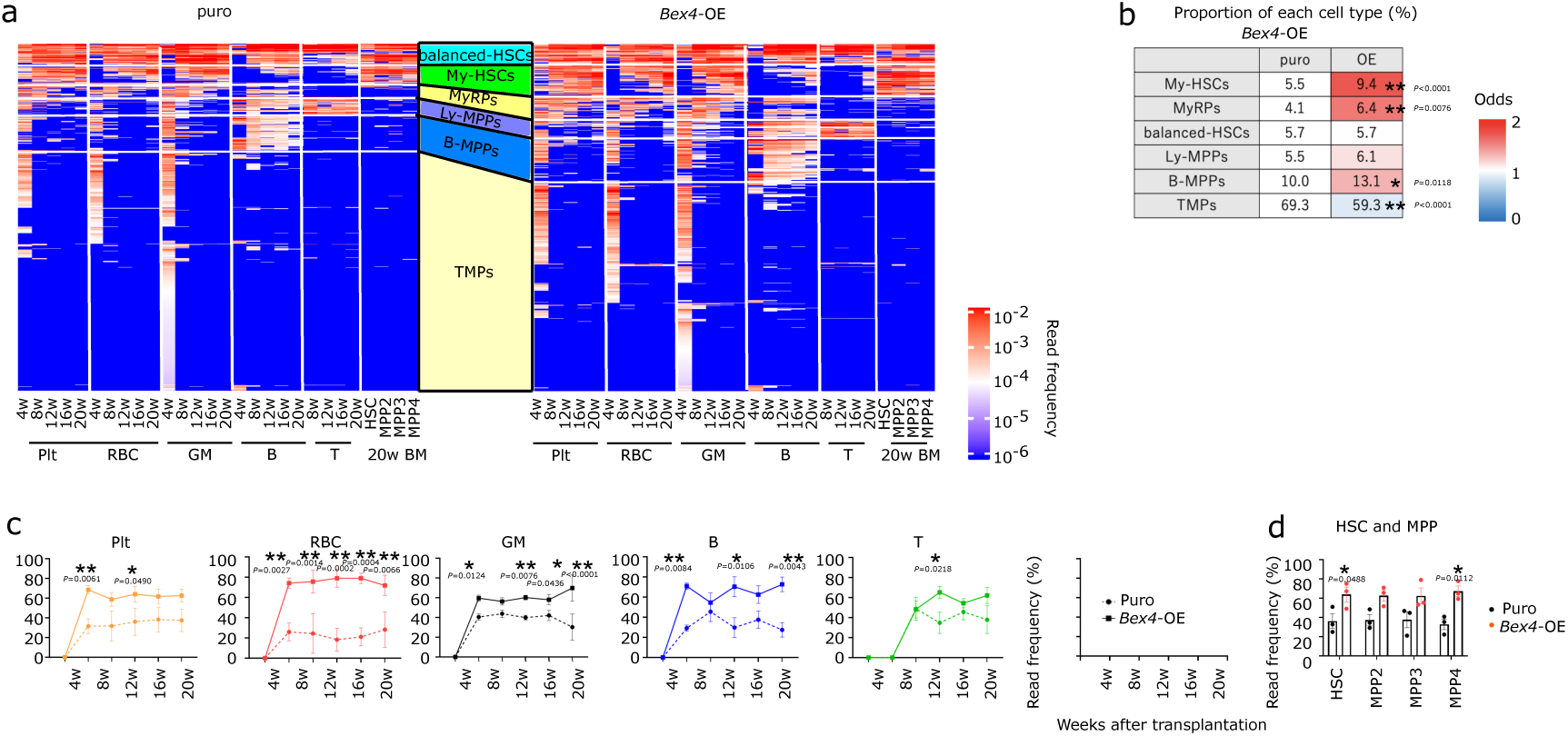
*Bex4* enhances My-HSC fate

## Discussion

Sequential clonal labeling with CP tracer has the ability to identify a clonal cell of origin and to track early fate decisions of an individual clone labeling approximately 600 progeny per HSC clone within 14 days (Supplementary Fig.4g,I,n,x,y). It does so with high barcode diversity (at least 10^10^=2×10^5^ random DNA barcode×2×10^5^ scratchpad) (Supplementary Fig.4a-d) without limitation on the number of cells for barcode recovery, enabling large-scale in vivo phylogenetic tracing of 181,695 descendant subclones originating from 847 single parent cells (Supplementary Fig. 4t), together with increasing the depth of lineage output information to detect 0.001–0.01% chimerism across multiple time points (Fig. 3i and Methods). This scale enabled comprehensive visualization of hematopoietic differentiation that had previously been impossible (Fig4b,i and Supplementary Fig.5e,6e,6f).

Using CP tracer has provided both new and confirmatory information about hematopoiesis. Single cell transplantation studies previously identified heterogeneity within the immunophenotypic HSC fraction^8,11,15^, including My-HSCs transitioning to balanced-HSCs with secondary transplantation^11^. The *Vwf*-GFP+ fraction that enriches My-HSCs, produced the *Vwf*-GFP+ fraction and the *Vwf*-GFP-fraction, while the *Vwf*-GFP-fraction enriched balanced-HSCs^8,9^. Together, these observations support a hierarchical relationship in which My-HSCs represent an upstream population relative to balanced-HSCs^11^. This is consistent with HSCs biased to myeloid cells having the highest stemness^6^ and lose stemness as they gain lymphoid competence^3,10^. However, unexpectedly, CP tracer analyses indicate that balanced-HSCs can also reverse their trajectory and transition to My-HSCs that then show a higher stemness^6^. Therefore, unidirectional differentiation that has generally been ascribed to hematopoiesis *in vivo* is not rigidly maintained.

This study used ex vivo transduction of cells and may not exactly correlate with endogenous cells not subjected to transplantation. Future studies will be necessary using lentiviral delivery *in vivo* that has been demonstrated to liver^58^, lungs^21^, brain^59^, and heart.^60^ Our system may be amenable to that approach, but to date we have not achieved adequate efficiencies of transduction in the endogenous hematopoietic system. Nevertheless, the transition from balanced-HSCs to My-HSCs proposed in our study was also supported under non-transplant settings, including DARLIN lineage tracing^20^(Supplementary Fig.5d,e) and analyses of human clonal hematopoiesis patients^31^.

Studies by others^61^ have indicated that phenotypic classification of HSC subpopulations may be more reflective of the time at which progeny were scored rather than a true distinction in competence. In our study, we have tried to incorporate this concept and defined My-HSCs as a population that predominantly produces myeloid lineage cells during primary transplantation, with lymphoid output being suppressed or delayed. Based on this definition, the observation that some My-HSCs exhibit significant lymphoid output upon secondary transplantation leads to two possible interpretations. One is that My-HSCs may have transitioned into balanced-HSCs; the other is that My-HSCs generated lymphoid progenitors with delayed kinetics, and these progenitors were responsible for lymphoid output in the secondary transplantation. As shown in Supplementary Fig.5m, the lymphoid output of My-HSCs upon secondary transplantation was largely attributable at the subclone level to lymphoid progenitors such as B-MPPs, while true balanced-HSC subclones were rarely observed. Although it remains possible that these lymphoid progenitors, including B-MPPs, were produced through balanced-HSC intermediates, our data suggest that the My-HSC-to-balanced-HSC transition is uncommon.

*Hhex* has previously been shown to regulate HSC self-renewal, and *Cdkn2a* overexpression caused by *Hhex* deficiency has been implicated in progenitor dysfunction following 5-FU treatment^52^. In this study, through single-clone-level analysis of *Hhex* depletion (Fig.6d,e) and stage-specific perturbation experiments targeting *Hhex* (Fig.6f-h), we demonstrate a role for *Hhex* in lineage bias at the HSC stage. Further, we discovered that the function of Hhex is as a myeloid lineage repressor and that Hhex effects are more pronounced in older hematopoietic cells. Coupled with the age related changes observed in *Hhex* expression, our data suggest a key role for *Hhex* in the accumulation of myeloid bias in hematopoiesis of the aged.

Along those lines, sequential labelling may add mechanistic insight to the pathobiology of aging. Myeloid biased hematopoiesis is a well-defined, but poorly understood characteristic of aged individuals and My-HSCs accumulate in aged adults, ^11,41,62,63^ ^64^. CP tracer revealed that balanced-HSCs largely transited toward My-HSCs and that while bidirectional transitions can occur, My-HSCs infrequently transited to balanced-HSCs. These dynamics favor the accumulation of My-HSCs over time and may participate in the altered hematopoiesis observed with aging^11,41,62,63^. Since *Hhex*, defined here by targeted CRISPR screening, serves as a suppressor of myeloid fate choice and its expression declines in aged mouse and human HSC, its role in the aging phenotype is also hypothesized. Further testing of whether altering *Hhex* regulation can affect hematopoietic aging is a topic for future study.

## Acknowledgements

We would like to thank all Scadden lab members, HSCI CRM Flow Cytometry Core, Animal facility at Center for Comparative Medicine at MGH and Bauer Core Facility at Harvard University for assistance with next generation sequencing. This research was supported by the Gerald and Darlene Jordan Professor of Medicine Chair, National Institutes of Health HL131477 and HL142494 and the Harvard Stem Cell to D.T.S and Grant-in-Aid for Challenging Exploratory Research from JSPS Grant Number 20K21613, the Japan Science and Technology Agency ACT-X program Grant Number JPMJAX25L9, grant from the The Chemo-Sero-Therapeutic Research Institute and grant from SENSHIN Medical Research Foundation to T.F. T.F. was supported by a JSPS Overseas Research Fellowships (202160002), The Uehara Memorial Foundation Postdoctoral Fellowship for Research Abroad (2020), The grant of the Mochida Memorial Foundation for Medical and Pharmaceutical Research (2020) and . We used generative AI tools to assist with editing and improving the clarity of the manuscript. Figures were created using FlowJo, Prism 10 and BioRender.com under a licensed agreement.

## Author contributions

Conceptualization, T.F. and D.T.S.; Methodology, T.F. and D.T.S.; Investigation, T.F., T.A.K., M.Y., Y.H.C., S.K., M.M., T.Z., L.H., R.Y.,; Data analysis, T.F., L.P.W., T.K., R.I.S., D.T.S.; Writing – Original Draft, T.F., Y.T., D.T.S.; Writing– Review & Editing: T.F., T.A.K., Y.T., M.M., T.Z., S.G., L.H., S.Y., D.T.S.; Visualization: T.F.; Project Administration, K.H., S.Y., R.I.S.,; Funding Acquisition, T.F. and D.T.S.; Supervision, D.T.S.

## Data and Material Availability

Materials are available, subject to material transfer agreement requests submitted to D. T. S. scRNA-sequencing, bulk RNA sequencing, ATAC-seq, Cut and Run and barcode sequencing data have been deposited at GEO “GSE255989”, “GSE312130”, “GSE312131”, “GSE312132” and BioProject ID”PRJNA1228300” respectively and are publicly available as of the date of publication. All other data available in the manuscript or supplementary materials are available from corresponding author upon reasonable request.

### Isolation of Hematopoietic Stem Cells

Bone marrow cells were harvested from C57BL/6J (JAX, 000664) or B6J.129(Cg)-Gt(ROSA)26Sortm1.1(CAG-cas9*,-EGFP)Fezh/J (JAX, 026179) mice by extracting femurs, tibias, pelvic bones, sternums, humeri, and vertebrae, followed by crushing. The bone marrow cells were passed through a 100 μm cell strainer (Corning, 431752), and ACK lysis (Quality Biological, 118-156-721) was performed using 5 mL at room temperature for 5 minutes. After adding 45 mL of PBS, the cells were passed through a 100 μm cell strainer and centrifuged for 5 minutes. The supernatant was discarded. Bone marrow cells were incubated with Lineage antibodies (1:1000) (Anti-mouse CD5-biotin Antibody (clone 53-7.3) (BioLegend, 100604), Anti-mouse/human CD45R/B220-biotin Antibody (clone RA3-6B2) (BioLegend, 103204), Anti-mouse TER-119-biotin Antibody (clone TER-119) (BioLegend, 116204), Anti-mouse Gr-1-biotin Antibody (clone RB6-8C5) (BioLegend, 108404), Anti-mouse/human CD11b-biotin Antibody (clone M1/70) (BioLegend, 101204)) in 1 mL for 10 minutes at 4 □. After adding 45 mL of PBS and centrifuging for 5 minutes, the supernatant was discarded. This process was repeated twice. The cells were then incubated with 10 mL of PBS and 90 μL of SA microbeads (Miltenyi Biotec, 130-090-858) for 15 minutes at 4 □. After adding 45 mL of PBS and centrifuging for 5 minutes, the supernatant was discarded. This process was repeated twice. Negative selection was then performed using LS columns (Miltenyi Biotec, 130-042-401). The cells from the negative fraction were centrifuged for 5 minutes, and the supernatant was removed. The cells were incubated with a cocktail of antibodies, including Anti-mouse CD150-PE Antibody (clone TC15-12F12.2) (BioLegend, 115904) (1:200), Anti-mouse cKit-PEcy7 Antibody (clone 2B8) (BioLegend, 105814) (1:400), Anti-mouse Sca1-APC Antibody (clone B7) (BioLegend, 108112) (1:400), Anti-mouse CD48-APCcy7 Antibody (clone HAM48-1) (BioLegend, 103432) (1:400), and SA-BV605 (BioLegend, 405229) (1:400), or Anti-mouse cKit-PEcy7 Antibody (1:400), Anti-mouse CD150-APC Antibody (clone TC15-12F12.2) (BioLegend, 115910) (1:400), Anti-mouse CD48-APCcy7 Antibody (1:400), Anti-mouse Sca1-BV421 Antibody (clone B7) (BioLegend, 108127) (1:400), and SA-BV605 (1:400) for 15 minutes at 4 □. Following the addition of 1 mL of PBS and centrifugation for 5 minutes, the supernatant was discarded. The cells were resuspended in 6 mL of PBS and sorted for the CD150^+^CD48^-^Lin^-^cKit^+^Sca1^+^ fraction using FACS Aria.

### Plasmid Preparation and DNA Assembly

The plasmid pMJ114 (Addgene Plasmid #85995) was digested using the restriction enzymes HpaI (NEB, R0105L) and NotI (NEB, R3186L) to linearize the vector. A synthetic DNA fragment containing the hU6 promoter and sgRNA scaffold was obtained from Azenta and used as an insert for HiFi DNA assembly. The linearized vector (50 ng) and synthetic DNA fragment (5 ng) were assembled using HiFi DNA Assembly Master Mix (5 µl) (NEB, E2621L) in a total reaction volume of 10 µl. The reaction mixture was incubated at 50°C for 1 hour. The plasmid containing scratchpad which was obtained from Azenta was digested using the restriction enzymes Kfl1 (Thermo Scientific, FD2164) to linearize the vector. A synthetic DNA fragment containing the sgRNA cassetes was obtained from Azenta and used as an insert for HiFi DNA assembly. The linearized vector (50 ng) and synthetic DNA fragment (5 ng) were assembled using HiFi DNA Assembly Master Mix (5 µl) (NEB, E2621L) in a total reaction volume of 10 µl. The reaction mixture was incubated at 50°C for 1 hour. Following the assembly, the reaction product was transformed into DH5α competent cells (NEB, C2987H) via heat shock. Colonies obtained from the transformation were screened to identify those containing the backbone plasmid with the desired insert. A second HiFi DNA assembly was performed using the previously generated backbone plasmid and an oligo pool containing randomized sequences. The reaction mixture consisted of 500 ng of the backbone vector, 45 nM of the oligo pool, and 50 µl of HiFi DNA Assembly Master Mix, with a total reaction volume of 100 µl. The reaction was incubated at 50°C for 1 hour. Following the assembly, the reaction mixture was purified using a DNA purification kit, with elution performed in 15 µl of molecular-grade water (MQ). Electroporation was performed by adding 5 µl of the purified assembly reaction to 100 µl of MegaX DH10B competent cells (Invitrogen, C640003). The mixture was subjected to electroporation at 2.0 kV, 200 Ω, and 25 µF. Immediately after electroporation, 4 ml of pre-warmed recovery medium was added to the cells, and the mixture was incubated at 37°C for 1 hour. Following recovery, 4 µl of the transformed cells was diluted to 1 ml with LB medium. A 100 µl aliquot of this dilution was spread on an LB plate containing the appropriate antibiotic. The number of resulting colonies was used to estimate the library diversity, which was calculated to be approximately 10^6^ variants. The remaining transformation mixture was spread across four large (15 cm) LB plates containing antibiotics. After overnight incubation, 10 ml of LB liquid medium was added to the surface of each plate, and the bacterial colonies were collected by gently scraping. This process was repeated two to three times, and the collected bacterial suspension was pooled into a single tube. The plasmid library was purified using a Maxi Prep kit, following the manufacturer’s protocol.

### Virus Production

HEK293T cells were seeded at a density of 4 × 10□cells in a 15 cm dish. After 18 hours, the medium was changed. Opti-MEM (Gibco, 31985-062) (900 μL), VSVG (3.5 μg), PAX2 (7 μg), vector (12 μg), and Fugene6 (67.5 μL) (Promega, E2692) were mixed and incubated at room temperature for 15 minutes, then added to the HEK293T cells. After 24 hours, the medium was changed again. An additional 24 hours later, the supernatant was collected and subjected to ultracentrifugation using an SW32 rotor at 20,000 rpm at 4°C for 2 hours. The supernatant was discarded, and the pellet was resuspended in PVA medium^65^ (200–400 μL).

### Virus Infection

RetroNectin (Takara, T100A) was applied to 96-well or 48-well plates at a volume of 100–250 μL per well and incubated at room temperature for 1 hour. The RetroNectin was then removed, and the plate was washed with PBS. Hematopoietic stem cells were seeded at a density of 1–3 × 10□cells per well for 96-well plate and 2-6 × 10□cells per well for 48-well plate. Virus solution (200–400 μL) was added to each well. After 18 hours, the hematopoietic stem cells were collected.

### Bone Marrow Hematopoietic Stem Cell Transplantation

Bone marrow was harvested from the femurs and tibias of C57BL/6J mice aged 8–12 weeks. A cell suspension containing 1 × 10□cells in 90 μL was prepared, and 10 μL of anti-Sca1-FITC Antibody (Miltenyi Biotec, 130123-124) was added, followed by a 10-minute incubation at 4°C. After adding 2 mL of PBS and centrifuging for 5 minutes, the supernatant was discarded. The cells were resuspended in 80 μL of PBS, and 20 μL of anti-FITC microbeads (Miltenyi Biotec, 130123-124) was added, followed by a 15-minute incubation at 4°C. After adding 2 mL of PBS and centrifuging for 5 minutes, the supernatant was discarded. Negative selection was then performed using LD columns (Miltenyi Biotec, 130-042-901). C57BL/6J mice aged 8–12 weeks were irradiated with 475 cGy at 4-hour intervals for two sessions. Hematopoietic stem cells (0.5–3 × 10□) and the Sca1-negative fraction (2 × 10□) were dissolved in 200 μL of PBS and injected into the tail veins of mice. For secondary transplantation, C57BL/6J mice aged 8–12 weeks were irradiated with 475 cGy at 4-hour intervals for two sessions. Subsequently, 1% or 10% of the primary bone marrow was injected into the tail veins of the mice.

### Peripheral Blood Sampling

At 4, 8, 12, and 16 weeks post-transplantation, approximately 120 μL of peripheral blood was collected from the tail vein of the mice into 20 μL of citrate-dextrose solution (Sigma Aldrich, C3821-50ML).

### Red Blood Cell Recovery

Of the total 140 μL of peripheral blood, 40 μL was transferred to a separate Eppendorf tube for red blood cell analysis. One microliter each of anti-mouse CD41-biotin Antibody (clone MWReg30) (BioLegend, 133930) and anti-mouse CD45-biotin Antibody (clone 30-F11) (BioLegend, 103104) was added, followed by a 10-minute incubation at room temperature. After adding 1 mL of PBS and centrifuging for 5 minutes, the supernatant was discarded. This process was repeated twice. The pellet was then resuspended in 600 μL of PBS, and 12 μL of SA microbeads (Miltenyi Biotec, 130-090-858) was added, followed by a 15-minute incubation at room temperature. After adding 1 mL of PBS and centrifuging for 5 minutes, the supernatant was discarded. This process was repeated twice. The sample was resuspended in 1 mL of PBS, loaded onto an LD column (Miltenyi Biotec, 130-042-901) pre-treated with 2 mL of PBS, and immediately loaded with an additional 2 mL of PBS. After the complete passage of liquid through the column, 2 mL of PBS was added to the column. Following the complete passage of the liquid through the column, the negative fraction was centrifuged, and the supernatant was removed.

### Platelet, Granulocyte/Monocyte, and lymphoid cells Recovery

One mL of ACK lysis buffer (Quality Biological, 118-156-721) was added to 100 μL of peripheral blood and incubated at room temperature for 15 minutes. After the addition of 1 mL of PBS and centrifugation for 5 minutes, the supernatant was discarded. Another 1 mL of ACK lysis buffer was added and incubated at room temperature for an additional 15 minutes. After adding 1 mL of PBS and centrifuging for 5 minutes, the supernatant was discarded. The cells were incubated at room temperature for 15 minutes with antibodies in a total volume of 100 μL: Anti-mouse CD19-APCcy7 Antibody (clone 6D5) (BioLegend, 115530) (1:400), Anti-mouse CD3ε-BV785 Antibody (BioLegend, 100355) (1:400), Anti-mouse CD41-APC (clone MWReg30) (BioLegend, 133914) (1:400), Anti-mouse TER-119-biotin Antibody (clone TER-119) (BioLegend, 116204) (1:400), Anti-mouse/human CD11b-PE Antibody (clone M1/70) (BioLegend, 101208) (1:1000) or Anti-mouse/human CD11b-BV421 Antibody (clone M1/70) (BioLegend, 101212) (1:400). After the addition of 1 mL of PBS and centrifugation for 5 minutes, the supernatant was discarded. The cells were resuspended in 6 mL of PBS. FACS Aria was used to sort the FSClowSSClowCD41+Ter119-CD11b-CD19-fraction as platelets and collect 1,250,000 cells. Immediately after collection, the cells were centrifuged at 1,200g for 5 minutes and the supernatant was removed. In the BFP^+^ or RFP^+^ fractions, cells were sorted based on the following markers: CD11b+CD19-as granulocytes/monocytes, CD11b-CD19b+ as B cells, CD11b-CD19b-CD3e+ as T cells.

### Barcode cDNA Synthesis from RNA

To platelet and RBC samples, 350 μL of RLT buffer (RNeasy, Qiagen, 74104) was added. After pipetting, 350 μL of 80% ethanol was added and mixed well. The mixture was loaded onto an RNeasy mini column. Following centrifugation at 12,000g for 1 minute, the liquid was discarded. Subsequently, 500 μL of RW1 buffer was added, and after centrifugation at 12,000g for 1 minute, the flow-through was discarded. Similarly, 500 μL of RPE buffer was added, and after centrifugation at 12,000g for 1 minute, the flow-through was discarded. The column was then placed into a recovery Eppendorf tube. Next, 25 μL of RNase-free water was added. The RNA was eluted by centrifugation at 12,000g for 1 minute. To the RNA, 2 μL each of Oligo(dT) and dNTP were added and incubated at 65°C for 5 minutes. After this, the mixture was placed on ice for 1 minute. Then, 2 μL each of DTT, RNase inhibitor, and SuperScript IV (Invitrogen/Thermo, 18091200) were added, along with 8 μL of buffer. The mixture was incubated in a thermal cycler with the following protocol: 23°C for 10 minutes, 55°C for 10 minutes, and 80°C for 10 minutes. To the cDNA product, 72 μl of AMPureXP beads (Beckman Coulter, A63881) were added and incubated at room temperature for 15 minutes. The mixture was placed into a magnetic stand for 5 minutes, and the supernatant was removed. Subsequently, 200 μl of 80% EtOH was added and left for 1 minute, and the supernatant was removed. The beads were resuspended in 30 μl of Nuclease-Free water (Invitrogen, AM9932) and incubated at room temperature for 15 minutes.

### Cell Recovery from Bone Marrow

Bone marrow cells were harvested from the femurs, tibias, pelvic bones, sternums, humeri, and spines of mice at 20 weeks post-transplantation, followed by crushing. The bone marrow cells were passed through a 100 μm cell strainer (Corning, 431752), and ACK lysis (Quality Biological, 118-156-721) was performed using 5 mL at room temperature for 5 minutes. After adding 45 mL of PBS, the mixture was passed through a 100 μm cell strainer and centrifuged for 5 minutes. The supernatant was discarded. One-fourth of the bone marrow cells were incubated for 15 minutes at 4°C with 1 mL of the following antibodies: Anti-mouse CD19-APCcy7 Antibody (BioLegend, 115530) (1:400), Anti-mouse CD3ε-BV785 Antibody (BioLegend, 100355) (1:400), Anti-mouse CD41-APC (BioLegend, 133914) (1:400), Anti-mouse TER-119-biotin Antibody (BioLegend, 116204) (1:400), Anti-mouse/human CD11b-PE Antibody (BioLegend, 101208) (1:1000) or Anti-mouse/human CD11b-BV421 Antibody (BioLegend, 101212) (1:400). After washing with 1 mL of PBS and centrifuging, the cell suspension was resuspended in 3 mL of PBS and sorted using Aria. In the BFP^+^ or RFP^+^ fractions, cells were sorted based on the following markers: CD11b^+^CD19^-^ as Granulocyte/Monocyte, CD11b^-^CD19^+^ as B cells, CD11b^-^CD19^-^CD3e^+^ as T cells, CD11b^-^CD19^-^CD41^+^ as Megakaryocytes, CD11b-CD19-Ter119^+^ as Erythrocytes. The remaining three-fourths of the bone marrow cells were incubated for 10 minutes at 4°C with Lineage antibodies (1:1000): Anti-mouse CD41-biotin Antibody, Anti-mouse CD5-biotin Antibody, Anti-mouse/human CD45R/B220-biotin Antibody, Anti-mouse TER-119-biotin Antibody, Anti-mouse Gr-1-biotin Antibody, Anti-mouse/human CD11b-biotin Antibody. After washing with 1 mL of PBS and centrifugation, the cell suspension was further incubated for 15 minutes at 4°C with a mixture of antibodies: Anti-mouse cKit-PEcy7 Antibody (1:400), Anti-mouse Sca1-BV785 Antibody (clone B7) (BioLegend, 108139) (1:400), Anti-mouse CD48-APCcy7 Antibody (1:400), SA-BV605 (1:400), Anti-mouse CD150-PE Antibody (1:200), Anti-mouse Flt3-APC Antibody (clone A2F10) (BioLegend, 135310) (1:50) or Anti-mouse CD150-APC Antibody (1:400), Anti-mouse Flt3-BV421 Antibody (BioLegend, 135313) (1:50). After washing with 1 mL of PBS and centrifugation, the cell suspension was resuspended in 6 mL of PBS and sorted using Aria. In the BFP^+^ or RFP^+^ fractions, cells were sorted based on the following markers: CD150^+^CD48^-^Flt3^-^Lin^-^cKit^+^Sca1^+^ as HSC, CD150^+^CD48^+^Flt3^-^Lin^-^cKit^+^Sca1^+^ as MPP2, CD150^-^CD48^+^Flt3^-^Lin^-^cKit^+^Sca1^+^ as MPP3, Flt3^+^Lin^-^cKit^+^Sca1^+^ as MPP4.

### Genomic DNA Recovery from Nucleated Cells

Genomic DNA was extracted from nucleated cells using the DNAeasy Blood Kit (Qiagen, 69504). Cells were suspended in 200 μl of PBS, and 20 μl of Protein K along with 200 μl of AL buffer were added. The suspension was incubated at 56°C for at least 15 minutes. Subsequently, 200 μl of 100% EtOH (Fisher, BP28184) was added, and the mixture was loaded onto a DNAeasy Blood column. After centrifugation at 12,000 rpm for 1 minute, the flow-through was discarded. Then, 500 μl of AW1 buffer was loaded onto the column. After centrifugation at 12,000 rpm for 1 minute, the flow-through was discarded. Next, 500 μl of AW2 buffer was loaded onto the column. After centrifugation at 12,000 rpm for 1 minute, the flow-through was discarded. The column was then centrifuged at 12,000 rpm for 1 minute to remove any residual wash buffer. Afterward, the column was placed into a recovery Eppendorf tube. Next, 25 μl of AE buffer was added, and the genomic DNA was eluted by centrifugation at 12,000g for 1 minute.

### Amplification of Barcodes

For both genomic DNA and cDNA (25 μl), 0.25 μl of 100 μM primers (F: CACGAGGTGGCAGTGGCCAGATAC, R for cDNA: GGCAAACAACAGATGGCTGGCAACTAG, R for genomic DNA: GGGACAGCAGAGATCCAGTTTGGTTAGTAC, F for Scratchpad: tctagacgtttaaactagcctgaggattcc, R for Scratchpad: ttgattcgaagttgagctcgactagctagG) and 25 μl of KOD One (Toyobo KMM-101) were added. The mixture was incubated in a thermal cycler with the following protocol: 98°C for 2 minutes, 22 cycles of (98°C for 10 seconds, 60°C for 30 seconds, and 68°C for 30 seconds), followed by 72°C for 10 minutes. To the PCR product, 90 μl of AMPureXP beads (Beckman Coulter, A63881) were added and incubated at room temperature for 15 minutes. The mixture was placed into a magnetic stand for 5 minutes, and the supernatant was removed. Subsequently, 200 μl of 80% EtOH was added and left for 1 minute, and the supernatant was removed. The beads were resuspended in 50 μl of Nuclease-Free water (Invitrogen, AM9932) and incubated at room temperature for 15 minutes. Finally, the mixture was placed into a magnetic stand for 5 minutes and the supernatant was collected. For secondary PCR, to 10 μl of the PCR products, 0.1 μl each of 100 μM primers (F: CTTTCCCTACACGACGCTCTTCCGATCTGACCTCCCTAGCAAACTGGGGCACAAG R: GAGTTCAGACGTGTGCTCTTCCGATCTGGCAAACAACAGATGGCTGGCAACTAG, F for Scratchpad: CTTTCCCTACACGACGCTCTTCCGATCTtctagacgtttaaactagcctgaggattcc, R for Scratchpad: GAGTTCAGACGTGTGCTCTTCCGATCTttgattcgaagttgagctcgactagctagG) and 10 μl of KOD One (Toyobo KMM-101) were added. The mixture was incubated in a thermal cycler with the following protocol: 98°C for 2 minutes, 10 cycles of (98°C for 10 seconds, 60°C for 30 seconds, and 68°C for 30 seconds), followed by 72°C for 10 minutes. To the PCR product, 36 μl of AMPureXP beads (Beckman Coulter, A63881) were added and incubated at room temperature for 15 minutes. The mixture was placed into a magnetic stand for 5 minutes, and the supernatant was removed. Subsequently, 200 μl of 80% EtOH was added and left for 1 minute, and the supernatant was removed. The beads were resuspended in 50 μl of Nuclease-Free water (Invitrogen, AM9932) and incubated at room temperature for 15 minutes. Finally, the mixture was placed into a magnetic stand for 5 minutes and the supernatant was collected. For indexing PCR, to 1 ng of secondary PCR product, 1 μl each of 10 μM primers (F: AATGATACGGCGACCACCGAGATCTACACNNNNNNNNACACTCTTTCCCTACACGACGC TCTTCCGATC*T, R: NEBNext® Multiplex Oligos for Illumina® (Index Primers Set, NEB E7335S, E7500S, E7710S)) and 10 μl of KOD One (Toyobo KMM-101) were added. The mixture was incubated in a thermal cycler with the following protocol: 98°C for 2 minutes, 5 cycles of (98°C for 10 seconds, 60°C for 30 seconds, and 68°C for 30 seconds), followed by 72°C for 10 minutes. To the indexing PCR product, 36 μl of AMPureXP beads (Beckman Coulter, A63881) were added and incubated at room temperature for 15 minutes. The mixture was placed into a magnetic stand for 5 minutes, and the supernatant was removed. Subsequently, 200 μl of 80% EtOH was added and left for 1 minute, and the supernatant was removed. The beads were resuspended in 50 μl of Nuclease-Free water (Invitrogen, AM9932) and incubated at room temperature for 15 minutes. Finally, the mixture was placed into a magnetic stand for 5 minutes and the supernatant was collected. Individual libraries were diluted to 4nM and pooled for sequencing. Pools were sequenced with 100 cycle run kits (75bp Read1, 8bp Index1, 8bp Index2) on the NextSeq 1000 P1 Sequencing System (Illumina). For Scratchpad, pools were sequenced with 300 cycle run kits (300bp Read1, 8bp Index1, 8bp Index2) on the NextSeq 1000 P1 Sequencing System (Illumina).

### The modification of the barcode system for phylogeny analysis

The rate at which mutations accumulate in scratchpad is a critical factor in tracing. If the accumulation of mutations is too rapid, labeling might be completed before a sufficient number of descendants are formed from HSCs. It leads to the labeling of an insufficient number of subclones from the pearent cell. On the other hand, if labeling is too slow, tracing might not refrect early fate segregation from HSCs. Editing efficiency at the target sequences is influenced by mismatches between the target sequence and its sgRNA. It is known that editing efficiency decreases more significantly when mutations occur closer to the PAM sequence^29^. To address this, we created three types of Scratchpads—“V1,” “V2,” and “V3”—with target sites exhibiting progressively lower editing efficiencies by CRISPR/Cas9 in the order of V3 < V2 < V1 (Supplementary Fig.4e,f). These three types were transduced into immunophenotypic HSCs from Cas9-EGFP mice, followed by an analysis of the editing pattern of the scratchpad 14 days after transduction (Supplementary Fig.4g). Most of the scratchpads produced in V1-V3 were unique to each clone (Supplementary Fig.4h). A decrease in editing efficiency and large deletions were observed in V2 and V3 (Supplementary Fig.4i,j). The number of subclones per clone was greatest in V3 (Supplementary Fig.4k). These findings indicate that V3 was the most suitable for phylogeny analysis. Based on these results, we used V3 in this paper. HSCs transduced with CP tracer were treated with CDK4/6 inhibitors, which are known to fully arrest cell-cycle progression^66^. If mutation accumulation in the CP tracer were strictly dependent on cell division, CDK4/6 inhibition would be expected to markedly reduce mutation acquisition. However, our analysis revealed no significant difference in the rate of mutation accumulation between CDK4/6-inhibited and control cells (Supplementary Fig.4l). No difference in engraftment after transplantation was observed between the CP tracer and the random DNA barcode vector (Supplementary Fig.4m).

### Single cell RNA-seq

scRNA-seq libraries were constructed using the Chromium Single Cell 3′v3.1 Reagent Kit (10x Genomics, PN-1000269, PN-1000127, and PN-1000213/2000240) according to the manufacturer’s protocol. Briefly, the post-sorting sample volume was reduced. Cells were then loaded into each channel with a target output of approximately 5,000 cells. Single cells were encapsulated into emulsion droplets using the Chromium Controller (10x Genomics). Reverse transcription, fragmentation, and indexing PCR were performed in a thermal cycler. cDNA and final libraries were purified using AMPureXP beads (Beckman Coulter, A63881). A random DNA barcode sequence was amplified using the SI primer (10x Genomics, PN-2000095) and the Reverse primer (GAGTTCAGACGTGTGCTCTTCCGATCTGACCTCCCTAGCAAACTGGGGCACAAG) following this protocol: 98°C for 2 minutes, 10 cycles of (98°C for 10 seconds, 60°C for 30 seconds, and 68°C for 30 seconds), and 72°C for 10 minutes. Indexing PCR was then performed using the SI primer and NEBNext® Multiplex Oligos for Illumina® (Index Primers Set, NEB E7335S) following this protocol: 98°C for 2 minutes, 5 cycles of (98°C for 10 seconds, 60°C for 30 seconds, and 68°C for 30 seconds), and 72°C for 10 minutes. Amplified cDNA and final libraries were evaluated on an Agilent BioAnalyzer using a High Sensitivity DNA Kit (Agilent Technologies, 5067-5584 and 5067-5585). Individual libraries were diluted to 4nM and pooled for sequencing. The pools were sequenced using 150-cycle run kits (26bp Read1, 8bp Index1, and 90bp Read2) on the Nova-seq SP Sequencing System (Illumina).

### Screening vector validation

First, 10 μg of vector was incubated with 5 μl of NsiI (NEB R0127S), 5 μl of MscI (NEB R0534S), and 10 μl of rCutsmart, making up to a total volume of 100 μl with water. The reaction was incubated at 37°C overnight. After gel electrophoresis, the vector was purified into 50 μl of water using a column (Qiagen Cat#: 28706). For 50 μl of the extracted vector, 5 μl of T4 polymerase (NEB M0203), 10 μl of rCutsmart, and water were added to make up the total volume of 100 μl. This mixture was incubated at 12°C for 15 minutes. The vector was then purified into 50 μl of water using a column (Qiagen Cat#: 28706). Next, for 50 μl of the extracted vector, 2 μl of Exonuclease V (NEB M0345S), 10 μl of 10 mM ATP, 10 μl of NEBuffer4, and water were added to make up the total volume of 100 μl. The mixture was incubated at 37°C for 30 minutes. The vector was purified into 50 μl of water using a column (Qiagen Cat#: 28706). For the next step, for 50 μl of the extracted vector, 5 μl of T4 ligase (NEB M0202L), 10 μl of T4 ligase buffer, and water were added to make up the total volume of 100 μl. The mixture was incubated at room temperature overnight. The vector was purified into 50 μl of water using a column (Qiagen Cat#: 28706). For 50 μl of the extracted vector, 50 μl of KOD One (Toyobo KMM-101), 1 μM of F primer (CTTTCCCTACACGACGCTCTTCCGATCTCTTGTGGAAAGGACGAAACACCG), and 1 μM of R primer (GAGTTCAGACGTGTGCTCTTCCGATCTGGCAAACAACAGATGGCTGGCAACTAGC) were added. The mixture was incubated in a thermal cycler with the following protocol: 98°C for 2 minutes, 15 cycles of (98°C for 10 seconds, 60°C for 30 seconds, and 68°C for 30 seconds), and 72°C for 10 minutes. To the PCR product, 180 μl of AMPureXP beads (Beckman Coulter, A63881) was added and incubated at room temperature for 15 minutes. The mixture was placed in a magnetic stand for 5 minutes, and the supernatant was removed. Then, 200 μl of 80% EtOH was added and left for 1 minute, after which the supernatant was removed. The beads were resuspended in 50 μl of Nuclease-Free water (Invitrogen, AM9932) and incubated at room temperature for 15 minutes. Finally, the mixture was placed in a magnetic stand for 5 minutes and the supernatant was collected. For 8 μl of the PCR product, 10 μl of KOD One (Toyobo KMM-101), 1 μl of NEBNext® multiplex primer index 1 primer (NEB E7335S), and 1 μl of 10 μM primer (AATGATACGGCGACCACCGAGATCTACACTATAGCCTACACTCTTTCCCTACACGACGCTCTTCCGATC*T) were added. The mixture was incubated in a thermal cycler with the following protocol: 98°C for 2 minutes, 15 cycles of (98°C for 10 seconds, 60°C for 30 seconds, and 68°C for 30 seconds), and 72°C for 10 minutes. To the PCR product, 36 μl of AMPureXP beads (Beckman Coulter, A63881) was added and incubated at room temperature for 15 minutes. The mixture was placed into a magnetic stand for 5 minutes, and the supernatant was removed. Then, 200 μl of 80% EtOH was added and left for 1 minute, after which the supernatant was removed. The beads were resuspended in 50 μl of Nuclease-Free water (Invitrogen, AM9932) and incubated at room temperature for 15 minutes. Finally, the mixture was placed in a magnetic stand for 5 minutes and the supernatant was collected.

### Combined sgRNA-random DNA barcode library validation

To create the random DNA barcode-sgRNA lookup table for the screening vector (Fig.6a), two methods were considered to examine the correspondence between the random DNA barcode and the sgRNA. In Method A, the entire region from the sgRNA to the random DNA barcode was amplified by PCR, followed by ligation of the ends of the PCR product to bring the sgRNA and random DNA barcode information into proximity. Subsequently, the adjacent sgRNA and random DNA barcode information were amplified by PCR. In Method B, the extra sequence between the sgRNA and the random DNA barcode was excised using a restriction enzyme, followed by ligation to bring the sgRNA and random DNA barcode information into proximity. The adjacent sgRNA and random DNA barcode information were then amplified by PCR (Supplementary Fig.8b). Template switching during PCR is known to cause information from multiple template molecules to be mixed into one molecule of PCR product^67^. To test for template switching, two vectors with known structures were mixed in equal amounts, and the mixture was processed using both methods. Then, the final products were sequenced (Supplementary Fig.8c). As a result, many template switches were observed with Method A. In contrast, only a few template switches were observed with Method B (Supplementary Fig.8b,c). Subsequently, the vector containing the random DNA barcode and the sgRNA library was validated using Method B. For vector validation screening, sgRNA and random DNA barcode sequences were extracted using Bartender. The sgRNA and random DNA barcode sequences on the same reads were connected using Python scripts, and sequences were clustered using Bartender^68^. Among 97.7% of the random DNA barcodes, over 70% of the sgRNA information originated from a single sgRNA (Supplementary Fig.8d), indicating the successful integration of the random DNA barcode and sgRNA information. The code is available at https://github.com/MGH-IMSUT/sgRNA-barcode_screening.

### RNAseq analysis

Gene editing for HSCs were performed as described previously^69^. Recombinant S. pyogenes Cas9 (S.p. Cas9 Nuclease V3, IDT) was complexed with single guide RNA (sgRNA, synthesized at IDT) at a molar ratio of 1:2.5 for 10 min at 25 C to form ribonucleoprotein (RNP) complexes. Sequences of sgRNA targeting murine Hhex (target sequence:AAAGGAAAGGCGGTCAAGTG). 10^5^ HSCs (CD150^+^CD48^-^Lin^-^cKit^+^Sca1^+^) were sorted from C57BL/6J (JAX, 000664) mouse and washed twice with PBS, pelleted, and resuspended in 20 μl electroporation buffer P3 (Lonza). The RNP duplex was gently added to the cells, and the suspension was transferred to a single 20 μl electroporation cuvette on a 16 well strip (P3 Primary Cell 96-well-Nucleofector Kit, Lonza). Electroporation was conducted using programs EO-100 on a 4D nucleofector device (Lonza). Cells were immediately recovered in pre-warmed medium and gently split-transferred into 24-well plates (Corning). One day after nucleofection, a medium change was performed, and further medium changes were performed every 2–3 days. To quantify knockout rates from bulk cultured cells, genomic DNA was extracted using DNAeasy Blood Kit (Qiagen, 69504). Polymerase chain reactions (PCR) were performed on 10ng of gDNA, formulated with 0.5 μM forward (GCCGGGGCCTTGGTTAACCAC) and reverse primers (ATGGTGTGTTTGGGTGTGAGG), 10 μl of KOD One (Toyobo KMM-101) in a 20 μl reaction. The PCR products were purified and subjected to Sanger sequencing (FASMAC, Japan) using the forward primer. ICE analysis^70^ confirmed that more than 93% of the alleles harbored frameshift mutations. For RNAseq analysis, after 7 days culture, 10000 cells of HSCs were transferred into 1.5 mL tubes and lysed in 600 ml Trizol LS reagent (Thermo Fisher Scientific, 10296010). Subsequently, RNA purification, library preparation, and next-generation sequencing were carried out by Tsukuba i-Laboratory, LLC. Libraries were generated using the SMARTer cDNA synthesis kit (Takara, A48571) and the high-output kit v2 (Illumina), followed by sequencing on a NextSeq 500 sequencer (Illumina) at paired end reads. DESeq2 package in R^71^ was utilized for data normalization and comparative analysis, and genes with an p < 0.01 were considered differentially expressed.

### ATAC-seq and Cut and Run

HSCs (CD150^+^CD48^-^Lin^-^cKit^+^Sca1^+^) were isolated from B6J.129(Cg)-Gt(ROSA)26Sortm1.1(CAG-cas9*,-EGFP)Fezh/J (JAX, 026179) or C57BL/6J (JAX, 000664) mouse bone marrow by FACS. The sorted HSCs were transduced with lentiviral vectors encoding either sgRNA or Hhex cDNA. Following transduction, cells were cultured for 7days. ATAC-seq was performed essentially as previously described by Buenrostro et al^72^. Briefly, sorted 2×10^4^ HSCs (BFP^+^CD150^+^CD48^-^Lin^-^cKit^+^Sca1^+^) were were pelleted by centrifugation (500 × g, 5 min, 4 °C), washed once with cold PBS, and resuspended in lysis buffer (10 mM Tris-HCl pH 7.4, 10 mM NaCl, 3 mM MgClL, 0.1% IGEPAL CA-630) to isolate nuclei. Immediately after lysis, nuclei were subjected to transposition using Tn5 transposase (Illumina Nextera kit) in a 50 µL reaction containing TD buffer and transposase, followed by incubation at 37 °C for 30 min. Transposed DNA was purified using a MinElute PCR purification kit (QIAGEN, 28004) and eluted in 10 µL of elution buffer (10 mM Tris-HCl, pH 8.0). Libraries were amplified by PCR using customized Nextera primers. Final libraries were purified using a PCR cleanup kit (QIAGEN, 28104) and eluted in 20 µL of elution buffer. CUT&RUN was performed using the CUT&RUN Assay Kit (Cell Signaling Technology, #86652) according to the manufacturer’s instructions. Briefly, nuclei derived from sorted 5×10^5^ HSCs (BFP^+^ after 7 days culture) were incubated with the indicated antibodies (Abcom, ab15828 and ab171870), followed by protein A–micrococcal nuclease treatment to release targeted chromatin fragments. DNA was subsequently purified using the DNA Purification Kit (Cell Signaling Technology, #14209) according to the manufacturer’s protocol. Sequencing reads were aligned to the mouse reference genome using Bowtie2^73^. Peaks were called using MACS2^73^ with a q-value threshold of < 0.05. Differentially accessible sites were identified using DiffBind^74^, applying a cutoff of q < 0.05.

### Barcode extraction and normarization clustering, and clonal assignment

Random DNA barcodes were extracted and clustered using Bartender^68^. For CP tracer, Raw sequencing reads were processed to extract combinations of random DNA barcodes and scratchpad sequences. First, the first 30 bp of each read were trimmed to remove sequences upstream of the random DNA barcode, which begins at position 31. Reads were then filtered to retain only those containing the invariant sequence GAGTC immediately downstream of the barcode, ensuring correct junction identification, using FASTX-toolkit (http://hannonlab.cshl.edu/fastx_toolkit), Seqkit^75^. Constant sequences located downstream of the scratchpad were removed using Cutadapt^76^, with an error tolerance of 20% and minimum overlap of 16 bp. Reads not terminating with the expected AGG sequence were excluded. Only sequences with a maximum length of 269 bp were retained to eliminate erroneous or truncated reads. The resulting FASTQ files were reformatted into a text file containing unique sequences and their supporting read counts, in a format compatible with Bartender. Barcode clustering was then performed using Bartender, with a maximum edit distance threshold of 27 bp, under the assumption that closely related reads originated from the same true sequence. This yielded consensus barcode–scratchpad combinations along with their read count distributions. Clones were defined by the unique random DNA barcodes. Subclones were defined by the unique allele (combination of random DNA barcodes and scratchpad variants) in each replicate, with the clone of origin identifiable from the random DNA barcode component. Analyses focused primarily on alleles with unique scratchpad variants across all clones. The sequencing depth ranges from approximately 100,000 to 1,000,000 reads for each lineage/timepoint. For the clone analysis, read counts below 50 were treated as zero; this threshold corresponds approximately to a chimerism level of 0.005–0.05%. Similarly, for the subclone analysis, read counts below 10 were set to zero; this threshold corresponds approximately to a chimerism level of 0.001–0.01%. the count data was normalized by log_10_(count frequency + 0.000001). Sampling scale was ∼1.25 million nucleated cells per lineage at each time point, corresponding to a theoretical detection limit of approximately 0.00008% chimerism.

### K-means clustering and annotation of clones and subclones

K-means clustering was performed using ComplexHeatmap^77,78^. Clusters were annotated based on their lineage output patterns, and annotated clusters were used for downstream analysis. Barcodes representing different conditions within the same mouse, distinguished by their initial letters (GTG, GAC, GGA, GCT), were separated after k-means clustering. This allowed the same cell-type classification criteria to be applied across different conditions.

### Phylogenetic tree reconstructiion

To identify sequence variations among alleles, each allele sequence was globally aligned to the reference sequence “GAGTCGAGACGCTGACGATACCTTAGTCGACGTGCGCGCTCTCTGACGACTGCACGAAAGTCGACGAAGGGATAGTATGCGTACACGCGACCTTCGTCTGCGCGCTACGTATCACTGGAGAGCGCGCTCGACGACTAAGGATGAGACATGCAGCCACATCCCTTCGTCGTCTGTCGTGCAGTCTCAGGATACGTAGCACGGAGACGAAGG” using the Needleman–Wunsch algorithm^79^ (Biopython pairwise2.align.globalms function^80^). For each alignment, mismatched or gapped regions were extracted and assigned unique mutation identifiers. Insertions were mapped to the nearest preceding reference position, while deletions and substitutions were assigned based on the aligned reference coordinates. A comprehensive mutation dictionary was generated by integrating all observed variants across alleles. Each allele was then represented by a set of mutation IDs corresponding to its deviations from the reference. For each clone, allelic mutation patterns were analyzed to infer phylogenetic relationships. Mutations were represented as sets of identifiers derived from the mutation list. Pairwise distances between alleles were computed using a weighted Jaccard distance, defined as:

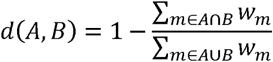

where A and B denote the sets of mutations in two alleles, and *W*_m_ is the weight of mutation *m,* defined as:

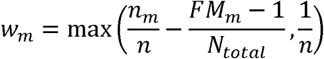

where *n*_m_ is the number of alleles harboring mutation ***m*** within a clone, ***n*** is the total number of alleles in the clone, *FM _m_* is the number of mutation ***m*** was detected within all allele, and ***N****_total_* is the number of all alleles. Phylogenetic trees were constructed using the Unweighted Pair Group Method with Arithmetic mean (UPGMA) algorithm^81^. A branching point in the phylogenetic tree is referred to as a node, representing a common ancestral progenitor from which descendant branches originate. Internal node distances were defined as half of the inter-cluster distance at the time of merging. We reconstructed allele-based phylogenetic trees and extracted node-associated features using custom Python scripts. First, mutation information for each allele was parsed from the input dataset, and each allele was represented as a set of mutations. For every internal node in the phylogenetic tree, common mutations were computed as the intersection of mutation sets across descendant alleles. Nodes sharing identical mutation sets with their parent were collapsed to simplify tree topology. Each internal node was then assigned a unique identifier (e.g., Node1, Node2, …) to enable systematic referencing. The hierarchical depth of each node was calculated recursively from the root. For each internal node, the set of descendant alleles was collected, and allele-associated parameter values were aggregated across descendants. Bootstrap analysis was performed by resampling mutation sites with replacement, reconstructing phylogenetic trees for each bootstrap replicate, and calculating bootstrap support values as the frequency at which each Node was reproduced across replicates. Lineage trees were simulated by iterative cell divisions with stochastic mutation acquisition. Starting from a single ancestral cell, cells underwent four rounds of division, with each parent generating four daughter cells per generation. During each division, daughter cells inherited all mutations from the parent cell and could additionally acquire one new mutation. Mutations were randomly selected according to their observed frequencies. To model inter-site depletion, only mutations whose both end target sites had not been previously edited were allowed to occur. When a newly acquired mutation overlapped with existing mutations, the overlapping mutations were replaced by the new mutation. After all divisions, the proportion of inter-site depletion was calculated, and mutation profiles of descendant cells were used for phylogenetic reconstruction. The similarity between the reconstructed trees and the corresponding ground-truth lineage trees was then evaluated.The code is available at https://github.com/MGH-IMSUT/CPTracer-pipeline-code.

### DARLIN data analysis

DARLIN data (https://zenodo.org/records/11929508) in which both transcriptomic profiles and DARLIN alleles were captured in parallel using scRNA-seq. Doxycycline (Dox) was administered to pregnant mice via orbital injection at embryonic day 17.0 (E17.0) in the dataset used in this study. Following Dox administration, unedited targets in the fetus continued to decrease over a 48-hour period^20^, indicating that the editing window extends for at least 48 hours (Supplementary Fig.5d). DARLIN alleles were recovered from single-cell RNA sequencing (scRNA-seq) data, enabling integration of cell type identity with subclonal information defined by DARLIN allele combinations. By aggregating the cell types associated with each allele combination, the lineage output of individual subclones was inferred (Supplementary Fig.5d). To reconstruct lineage relationships, DARLIN alleles were compared across subclones to identify sibling subclones based on shared mutations. Mutations shared among all sibling subclones were considered to originate from the parental clone. The lineage output of each inferred parent clone was defined as the sum of the outputs of its descendant subclones (Supplementary Fig.5d). Analysis was restricted to parent clones defined by allele combinations with a probability of random occurrence below 5 × 10LL, following criteria established in the original study^20^. Under this threshold, six balanced HSC clones contributing to myeloid, lymphoid, and HSC compartments were identified (Supplementary Fig5e).

### Estimation of immunophynotypic HSC dose required for 63.2% secondary engraftment

Extreme limiting dilution analysis (ELDA)^82^ was used to estimated immunophenotypic HSC dose required to achieve secondary bone marrow reconstitution of immunophenotypic HSCs with a 63.2% probability. Because only the frequency of immunophenotypic HSCs in the bone marrow at 20 weeks after primary transplantation was available, the total number of immunophenotypic HSCs was assumed to be 10,000 for the purpose of ELDA calculations. Using this assumed total and the binary outcome of immunophenotypic HSC detection in the bone marrow at 20 weeks after secondary transplantation, ELDA was performed for each HSC subtype–derived cell population. Estimated doses were subsequently normalized to those derived from P-HSCs. Sensitivity analyses were performed by varying the assumed total number of immunophenotypic HSCs (2,500, 10,000, 25,000, and 100,000). The resulting estimates differed by less than 0.1%, confirming that the results were robust to the assumed total number of immunophenotypic HSCs.

### scRNA-seq data analysis

Single-cell RNA-seq FASTQ files were analyzed with 10x Genomics cloud analysis via Cell Ranger Count v7.1.0^83^. The resulting filtered feature-barcode matrix was then subjected to downstream analysis using Seurat^84^. Of the 14,411 cells, 13,701 cells meeting the criteria nFeature_RNA > 300 & nFeature_RNA < 10,000 and percent.mt < 5 were used for downstream processing. Dimensionality reduction and clustering were performed using UMAP, and clusters corresponding to differentiated cells or cells in the cell cycle were removed based on marker gene analysis (Supplementary Fig.7a–d). The remaining 11,717 cells were used for subsequent analyses (Supplementary Fig.7a). Subsequently, batch correction was performed using Harmony^85^, followed by dimensionality reduction with UMAP (Supplementary Fig.7e,f). Random DNA barcodes from the single-cell RNA-seq library were extracted using FASTX-toolkit, Python scripts and bartender. Sequences were then clustered using bartender. For each single cell, the random DNA barcode with the highest read count was assigned as the clone identity of that cell. Using this method, the random DNA barcodes were successfully identified in 10,487 out of 11,717 cells (89.5 %) (Supplementary Fig.7a). The code is available at https://github.com/MGH-IMSUT/single_cell_RNA-seq_annotation. MolO scores were calculated using the following a publicly available pipeline (https://github.com/fionahamey/hscScore)^86^. Briefly, MolO scores were inferred by applying a pretrained hscScore regression model to log-transformed, total-count–normalized expression values derived from our scRNA-seq data. Gene sets used to assess gene signatures were obtained from Enrichr^87^, SenMayo^88^ FerrDb^89^, and Reactome^90^. To assess the spatial distributions of different HSC subsets in UMAP space, we performed two-dimensional kernel density estimation (KDE). To evaluate the statistical significance of the spatial distribution difference, we performed a permutation test by randomly shuffling cluster labels and calculating the sum of squared differences between the KDE densities. RNA velocity was analyzed by scVelo^37^. The gene regulatory network was analyzed using SCENIC^42,43^. Genes with log2(fold change)≥0.2 and adjusted p-value≤0.01 in P-HSCs/PRG-HSCs compared with balanced-HSCs were defined as My-HSC marker genes, whereas genes with log2(fold change)≤ − 0.2 and adjusted p-value≤0.01 were defined as balanced-HSC marker genes. Using CellOracle^44^, we performed in silico gene knockout perturbation analysis and inferred the resulting gene expression state of each single cell. My-HSC and balanced-HSC signature scores were then calculated separately using the AddModuleScore function in Seurat. To identify genes with strong perturbation effects on each signature, robust Z-scores were calculated across perturbed genes using the median absolute deviation method. Specifically, each perturbation score was normalized as robust Z-score=(x−median)/scaled MAD, where scaled MAD was defined as MAD×1.4826 to approximate the standard deviation under a normal distribution. Genes with absolute robust Z-scores>4 were selected as strong-effect perturbation genes.

## Statistics and reproducibility

Statistical and graphical data analyses were performed using GraphPad Prism 9 or other specified software. The number of samples and details of the statistical tests used are provided in the figures or figure legends.

## Declaration of generative AI and AI-assisted technologies in the manuscript preparation process

During the preparation of this work the authors used ChatGPT (OpenAI) to assist with language editing and improvement of readability. After using this tool, the authors reviewed and edited the content as needed and take full responsibility for the content of the published article.

## References

1. Busch, K., Klapproth, K., Barile, M., Flossdorf, M., Holland-Letz, T., Schlenner, S.M., Reth, M., Höfer, T., and Rodewald, H.R. (2015). Fundamental properties of unperturbed haematopoiesis from stem cells in vivo. Nature 518, 542–546. 10.1038/nature14242.

2. Sawai, C.M., Babovic, S., Upadhaya, S., Knapp, D., Lavin, Y., Lau, C.M., Goloborodko, A., Feng, J., Fujisaki, J., Ding, L., et al. (2016). Hematopoietic Stem Cells Are the Major Source of Multilineage Hematopoiesis in Adult Animals. Immunity 45, 597–609. 10.1016/j.immuni.2016.08.007.

3. Chen, J.Y., Miyanishi, M., Wang, S.K., Yamazaki, S., Sinha, R., Kao, K.S., Seita, J., Sahoo, D., Nakauchi, H., and Weissman, I.L. (2016). Hoxb5 marks long-term haematopoietic stem cells and reveals a homogenous perivascular niche. Nature 530, 223–227. 10.1038/nature16943.

4. Ito, K., Turcotte, R., Cui, J., Zimmerman, S.E., Pinho, S., Mizoguchi, T., Arai, F., Runnels, J.M., Alt, C., Teruya-Feldstein, J., et al. (2016). Self-renewal of a purified Tie2+ hematopoietic stem cell population relies on mitochondrial clearance. Science 354, 1156–1160. 10.1126/science.aaf5530.

5. Loeffler, D., Wehling, A., Schneiter, F., Zhang, Y., Müller-Bötticher, N., Hoppe, P.S., Hilsenbeck, O., Kokkaliaris, K.D., Endele, M., and Schroeder, T. (2019). Asymmetric lysosome inheritance predicts activation of haematopoietic stem cells. Nature 573, 426–429. 10.1038/s41586-019-1531-6.

6. Rodriguez-Fraticelli, A.E., Weinreb, C., Wang, S.W., Migueles, R.P., Jankovic, M., Usart, M., Klein, A.M., Lowell, S., and Camargo, F.D. (2020). Single-cell lineage tracing unveils a role for TCF15 in haematopoiesis. Nature 583, 585–589. 10.1038/s41586-020-2503-6.

7. Dykstra, B., Kent, D., Bowie, M., McCaffrey, L., Hamilton, M., Lyons, K., Lee, S.J., Brinkman, R., and Eaves, C. (2007). Long-term propagation of distinct hematopoietic differentiation programs in vivo. Cell Stem Cell 1, 218–229. 10.1016/j.stem.2007.05.015.

8. Carrelha, J., Meng, Y., Kettyle, L.M., Luis, T.C., Norfo, R., Alcolea, V., Boukarabila, H., Grasso, F., Gambardella, A., Grover, A., et al. (2018). Hierarchically related lineage-restricted fates of multipotent haematopoietic stem cells. Nature 554, 106–111. 10.1038/nature25455.

9. Sanjuan-Pla, A., Macaulay, I.C., Jensen, C.T., Woll, P.S., Luis, T.C., Mead, A., Moore, S., Carella, C., Matsuoka, S., Bouriez Jones, T., et al. (2013). Platelet-biased stem cells reside at the apex of the haematopoietic stem-cell hierarchy. Nature 502, 232–236. 10.1038/nature12495.

10. Morita, Y., Ema, H., and Nakauchi, H. (2010). Heterogeneity and hierarchy within the most primitive hematopoietic stem cell compartment. J Exp Med 207, 1173–1182. 10.1084/jem.20091318.

11. Yamamoto, R., Wilkinson, A.C., Ooehara, J., Lan, X., Lai, C.Y., Nakauchi, Y., Pritchard, J.K., and Nakauchi, H. (2018). Large-Scale Clonal Analysis Resolves Aging of the Mouse Hematopoietic Stem Cell Compartment. Cell Stem Cell 22, 600–607.e604. 10.1016/j.stem.2018.03.013.

12. Rodriguez-Fraticelli, A.E., Wolock, S.L., Weinreb, C.S., Panero, R., Patel, S.H., Jankovic, M., Sun, J., Calogero, R.A., Klein, A.M., and Camargo, F.D. (2018). Clonal analysis of lineage fate in native haematopoiesis. Nature 553, 212–216. 10.1038/nature25168.

13. Pei, W., Shang, F., Wang, X., Fanti, A.K., Greco, A., Busch, K., Klapproth, K., Zhang, Q., Quedenau, C., Sauer, S., et al. (2020). Resolving Fates and Single-Cell Transcriptomes of Hematopoietic Stem Cell Clones by PolyloxExpress Barcoding. Cell Stem Cell 27, 383–395.e388. 10.1016/j.stem.2020.07.018.

14. Pei, W., Feyerabend, T.B., Rössler, J., Wang, X., Postrach, D., Busch, K., Rode, I., Klapproth, K., Dietlein, N., Quedenau, C., et al. (2017). Polylox barcoding reveals haematopoietic stem cell fates realized in vivo. Nature 548, 456–460. 10.1038/nature23653.

15. Yamamoto, R., Morita, Y., Ooehara, J., Hamanaka, S., Onodera, M., Rudolph, K.L., Ema, H., and Nakauchi, H. (2013). Clonal analysis unveils self-renewing lineage-restricted progenitors generated directly from hematopoietic stem cells. Cell 154, 1112–1126. 10.1016/j.cell.2013.08.007.

16. Yu, V.W.C., Yusuf, R.Z., Oki, T., Wu, J., Saez, B., Wang, X., Cook, C., Baryawno, N., Ziller, M.J., Lee, E., et al. (2017). Epigenetic Memory Underlies Cell-Autonomous Heterogeneous Behavior of Hematopoietic Stem Cells. Cell 168, 944–945. 10.1016/j.cell.2017.02.010.

17. Challen, G.A., Boles, N.C., Chambers, S.M., and Goodell, M.A. (2010). Distinct hematopoietic stem cell subtypes are differentially regulated by TGF-beta1. Cell Stem Cell 6, 265–278. 10.1016/j.stem.2010.02.002.

18. Pinho, S., Marchand, T., Yang, E., Wei, Q., Nerlov, C., and Frenette, P.S. (2018). Lineage-Biased Hematopoietic Stem Cells Are Regulated by Distinct Niches. Dev Cell 44, 634–641.e634. 10.1016/j.devcel.2018.01.016.

19. Bowling, S., Sritharan, D., Osorio, F.G., Nguyen, M., Cheung, P., Rodriguez-Fraticelli, A., Patel, S., Yuan, W.C., Fujiwara, Y., Li, B.E., et al. (2020). An Engineered CRISPR-Cas9 Mouse Line for Simultaneous Readout of Lineage Histories and Gene Expression Profiles in Single Cells. Cell 181, 1410–1422.e1427. 10.1016/j.cell.2020.04.048.

20. Li, L., Bowling, S., McGeary, S.E., Yu, Q., Lemke, B., Alcedo, K., Jia, Y., Liu, X., Ferreira, M., Klein, A.M., et al. (2023). A mouse model with high clonal barcode diversity for joint lineage, transcriptomic, and epigenomic profiling in single cells. Cell. 10.1016/j.cell.2023.09.019.

21. Yang, D., Jones, M.G., Naranjo, S., Rideout, W.M, 3rd., Min, K.H.J., Ho, R., Wu, W., Replogle, J.M., Page, J.L., Quinn, J.J., et al. (2022). Lineage tracing reveals the phylodynamics, plasticity, and paths of tumor evolution. Cell 185, 1905–1923.e1925. 10.1016/j.cell.2022.04.015.

22. Kong, W., Biddy, B.A., Kamimoto, K., Amrute, J.M., Butka, E.G., and Morris, S.A. (2020). CellTagging: combinatorial indexing to simultaneously map lineage and identity at single-cell resolution. Nat Protoc 15, 750–772. 10.1038/s41596-019-0247-2.

23. Naik, S.H., Perié, L., Swart, E., Gerlach, C., van Rooij, N., de Boer, R.J., and Schumacher, T.N. (2013). Diverse and heritable lineage imprinting of early haematopoietic progenitors. Nature 496, 229–232. 10.1038/nature12013.

24. Fan, X., Wu, C., Truitt, L.L., Espinoza, D.A., Sellers, S., Bonifacino, A., Zhou, Y., Cordes, S.F., Krouse, A., Metzger, M., et al. (2020). Clonal tracking of erythropoiesis in rhesus macaques. Haematologica 105, 1813–1824. 10.3324/haematol.2019.231811.

25. Feng, J., Jang, G., Esteva, E., Adams, N.M., Jin, H., and Reizis, B. (2024). Clonal barcoding of endogenous adult hematopoietic stem cells reveals a spectrum of lineage contributions. Proc Natl Acad Sci U S A 121, e2317929121. 10.1073/pnas.2317929121.

26. Calabria, A., Spinozzi, G., Cesana, D., Buscaroli, E., Benedicenti, F., Pais, G., Gazzo, F., Scala, S., Lidonnici, M.R., Scaramuzza, S., et al. (2024). Long-term lineage commitment in haematopoietic stem cell gene therapy. Nature 636, 162–171. 10.1038/s41586-024-08250-x.

27. McKenna, A., Findlay, G.M., Gagnon, J.A., Horwitz, M.S., Schier, A.F., and Shendure, J. (2016). Whole-organism lineage tracing by combinatorial and cumulative genome editing. Science 353, aaf7907. 10.1126/science.aaf7907.

28. Perli, S.D., Cui, C.H., and Lu, T.K. (2016). Continuous genetic recording with self-targeting CRISPR-Cas in human cells. Science 353. 10.1126/science.aag0511.

29. Chan, M.M., Smith, Z.D., Grosswendt, S., Kretzmer, H., Norman, T.M., Adamson, B., Jost, M., Quinn, J.J., Yang, D., Jones, M.G., et al. (2019). Molecular recording of mammalian embryogenesis. Nature 570, 77–82. 10.1038/s41586-019-1184-5.

30. Weng, C., Yu, F., Yang, D., Poeschla, M., Liggett, L.A., Jones, M.G., Qiu, X., Wahlster, L., Caulier, A., Hussmann, J.A., et al. (2024). Deciphering cell states and genealogies of human haematopoiesis. Nature 627, 389–398. 10.1038/s41586-024-07066-z.

31. Stable, intrinsically programmed lineage restriction of human hematopoietic stem cells. (2025). Nat Genet 57, 2958–2959. 10.1038/s41588-025-02425-6.

32. Gong, W., Granados, A.A., Hu, J., Jones, M.G., Raz, O., Salvador-Martínez, I., Zhang, H., Chow, K.K., Kwak, I.Y., Retkute, R., et al. (2021). Benchmarked approaches for reconstruction of in vitro cell lineages and in silico models of C. elegans and M. musculus developmental trees. Cell Syst 12, 810–826.e814. 10.1016/j.cels.2021.05.008.

33. Jones, M.G., Khodaverdian, A., Quinn, J.J., Chan, M.M., Hussmann, J.A., Wang, R., Xu, C., Weissman, J.S., and Yosef, N. (2020). Inference of single-cell phylogenies from lineage tracing data using Cassiopeia. Genome Biol 21, 92. 10.1186/s13059-020-02000-8.

34. Wilson, N.K., Kent, D.G., Buettner, F., Shehata, M., Macaulay, I.C., Calero-Nieto, F.J., Sanchez Castillo, M., Oedekoven, C.A., Diamanti, E., Schulte, R., et al. (2015). Combined Single-Cell Functional and Gene Expression Analysis Resolves Heterogeneity within Stem Cell Populations. Cell Stem Cell 16, 712–724. 10.1016/j.stem.2015.04.004.

35. Tirosh, I., Izar, B., Prakadan, S.M., Wadsworth, M.H, 2nd., Treacy, D., Trombetta, J.J., Rotem, A., Rodman, C., Lian, C., Murphy, G., et al. (2016). Dissecting the multicellular ecosystem of metastatic melanoma by single-cell RNA-seq. Science 352, 189–196. 10.1126/science.aad0501.

36. Becht, E., McInnes, L., Healy, J., Dutertre, C.A., Kwok, I.W.H., Ng, L.G., Ginhoux, F., and Newell, E.W. (2018). Dimensionality reduction for visualizing single-cell data using UMAP. Nat Biotechnol. 10.1038/nbt.4314.

37. Bergen, V., Lange, M., Peidli, S., Wolf, F.A., and Theis, F.J. (2020). Generalizing RNA velocity to transient cell states through dynamical modeling. Nat Biotechnol 38, 1408–1414. 10.1038/s41587-020-0591-3.

38. Lambert, S.A., Jolma, A., Campitelli, L.F., Das, P.K., Yin, Y., Albu, M., Chen, X., Taipale, J., Hughes, T.R., and Weirauch, M.T. (2018). The Human Transcription Factors. Cell 172, 650–665. 10.1016/j.cell.2018.01.029.

39. Kataoka, K., Sato, T., Yoshimi, A., Goyama, S., Tsuruta, T., Kobayashi, H., Shimabe, M., Arai, S., Nakagawa, M., Imai, Y., et al. (2011). Evi1 is essential for hematopoietic stem cell self-renewal, and its expression marks hematopoietic cells with long-term multilineage repopulating activity. J Exp Med 208, 2403–2416. 10.1084/jem.20110447.

40. Calvanese, V., Nguyen, A.T., Bolan, T.J., Vavilina, A., Su, T., Lee, L.K., Wang, Y., Lay, F.D., Magnusson, M., Crooks, G.M., et al. (2019). MLLT3 governs human haematopoietic stem-cell self-renewal and engraftment. Nature 576, 281–286. 10.1038/s41586-019-1790-2.

41. Meng, Y., Carrelha, J., Drissen, R., Ren, X., Zhang, B., Gambardella, A., Valletta, S., Thongjuea, S., Jacobsen, S.E., and Nerlov, C. (2023). Epigenetic programming defines haematopoietic stem cell fate restriction. Nat Cell Biol 25, 812–822. 10.1038/s41556-023-01137-5.

42. Van de Sande, B., Flerin, C., Davie, K., De Waegeneer, M., Hulselmans, G., Aibar, S., Seurinck, R., Saelens, W., Cannoodt, R., Rouchon, Q., et al. (2020). A scalable SCENIC workflow for single-cell gene regulatory network analysis. Nat Protoc 15, 2247–2276. 10.1038/s41596-020-0336-2.

43. Aibar, S., González-Blas, C.B., Moerman, T., Huynh-Thu, V.A., Imrichova, H., Hulselmans, G., Rambow, F., Marine, J.C., Geurts, P., Aerts, J., et al. (2017). SCENIC: single-cell regulatory network inference and clustering. Nat Methods 14, 1083–1086. 10.1038/nmeth.4463.

44. Kamimoto, K., Stringa, B., Hoffmann, C.M., Jindal, K., Solnica-Krezel, L., and Morris, S.A. (2023). Dissecting cell identity via network inference and in silico gene perturbation. Nature 614, 742–751. 10.1038/s41586-022-05688-9.

45. Santaguida, M., Schepers, K., King, B., Sabnis, A.J., Forsberg, E.C., Attema, J.L., Braun, B.S., and Passegué, E. (2009). JunB protects against myeloid malignancies by limiting hematopoietic stem cell proliferation and differentiation without affecting self-renewal. Cancer Cell 15, 341–352. 10.1016/j.ccr.2009.02.016.

46. Jackson, J.T., O’Donnell, K., Light, A., Goh, W., Huntington, N.D., Tarlinton, D.M., and McCormack, M.P. (2020). Hhex regulates murine lymphoid progenitor survival independently of Stat5 and Cdkn2a. Eur J Immunol 50, 959–971. 10.1002/eji.201948371.

47. Goodings, C., Smith, E., Mathias, E., Elliott, N., Cleveland, S.M., Tripathi, R.M., Layer, J.H., Chen, X., Guo, Y., Shyr, Y., et al. (2015). Hhex is Required at Multiple Stages of Adult Hematopoietic Stem and Progenitor Cell Differentiation. Stem Cells 33, 2628–2641. 10.1002/stem.2049.

48. Jackson, J.T., Nasa, C., Shi, W., Huntington, N.D., Bogue, C.W., Alexander, W.S., and McCormack, M.P. (2015). A crucial role for the homeodomain transcription factor Hhex in lymphopoiesis. Blood 125, 803–814. 10.1182/blood-2014-06-579813.

49. Migueles, R.P., Shaw, L., Rodrigues, N.P., May, G., Henseleit, K., Anderson, K.G., Goker, H., Jones, C.M., de Bruijn, M.F., Brickman, J.M., and Enver, T. (2017). Transcriptional regulation of Hhex in hematopoiesis and hematopoietic stem cell ontogeny. Dev Biol 424, 236–245. 10.1016/j.ydbio.2016.12.021.

50. Oguro, H., Ding, L., and Morrison, S.J. (2013). SLAM family markers resolve functionally distinct subpopulations of hematopoietic stem cells and multipotent progenitors. Cell Stem Cell 13, 102–116. 10.1016/j.stem.2013.05.014.

51. Pietras, E.M., Reynaud, D., Kang, Y.A., Carlin, D., Calero-Nieto, F.J., Leavitt, A.D., Stuart, J.M., Göttgens, B., and Passegué, E. (2015). Functionally Distinct Subsets of Lineage-Biased Multipotent Progenitors Control Blood Production in Normal and Regenerative Conditions. Cell Stem Cell 17, 35–46. 10.1016/j.stem.2015.05.003.

52. Jackson, J.T., Shields, B.J., Shi, W., Di Rago, L., Metcalf, D., Nicola, N.A., and McCormack, M.P. (2017). Hhex Regulates Hematopoietic Stem Cell Self-Renewal and Stress Hematopoiesis via Repression of Cdkn2a. Stem Cells 35, 1948–1957. 10.1002/stem.2648.

53. Laidlaw, B.J., Duan, L., Xu, Y., Vazquez, S.E., and Cyster, J.G. (2020). The transcription factor Hhex cooperates with the corepressor Tle3 to promote memory B cell development. Nat Immunol 21, 1082–1093. 10.1038/s41590-020-0713-6.

54. Kim, K.M., Mura-Meszaros, A., Tollot, M., Krishnan, M.S., Gründl, M., Neubert, L., Groth, M., Rodriguez-Fraticelli, A., Svendsen, A.F., Campaner, S., et al. (2022). Taz protects hematopoietic stem cells from an aging-dependent decrease in PU.1 activity. Nat Commun 13, 5187. 10.1038/s41467-022-32970-1.

55. Aksöz, M., Gafencu, G.A., Stoilova, B., Buono, M., Zhang, Y., Turkalj, S., Meng, Y., Jakobsen, N.A., Metzner, M., Clark, S.A., et al. (2024). Hematopoietic stem cell heterogeneity and age-associated platelet bias are evolutionarily conserved. Sci Immunol 9, eadk3469. 10.1126/sciimmunol.adk3469.

56. Wang, P., Chen, Z., Zhang, H., Lu, Y., Zhou, L., Gong, C., An, D., Sang, X., Wang, K., Hao, M., and Cao, G. (2025). Biological potential and mechanisms of Brain-Expressed X-linked family proteins in cancers: an updated review. J Adv Res. 10.1016/j.jare.2025.07.034.

57. Manford, A.G., Mena, E.L., Shih, K.Y., Gee, C.L., McMinimy, R., Martínez-González, B., Sherriff, R., Lew, B., Zoltek, M., Rodríguez-Pérez, F., et al. (2021). Structural basis and regulation of the reductive stress response. Cell 184, 5375–5390.e5316. 10.1016/j.cell.2021.09.002.

58. Dalsgaard, T., Cecchi, C.R., Askou, A.L., Bak, R.O., Andersen, P.O., Hougaard, D., Jensen, T.G., Dagnæs-Hansen, F., Mikkelsen, J.G., Corydon, T.J., and Aagaard, L. (2018). Improved Lentiviral Gene Delivery to Mouse Liver by Hydrodynamic Vector Injection through Tail Vein. Mol Ther Nucleic Acids 12, 672–683. 10.1016/j.omtn.2018.07.005.

59. Humbel, M., Ramosaj, M., Zimmer, V., Regio, S., Aeby, L., Moser, S., Boizot, A., Sipion, M., Rey, M., and Déglon, N. (2021). Maximizing lentiviral vector gene transfer in the CNS. Gene Ther 28, 75–88. 10.1038/s41434-020-0172-6.

60. Higuchi, K., Ayach, B., Sato, T., Chen, M., Devine, S.P., Rasaiah, V.I., Dawood, F., Yanagisawa, T., Tei, C., Takenaka, T., et al. (2009). Direct injection of kit ligand-2 lentivirus improves cardiac repair and rescues mice post-myocardial infarction. Mol Ther 17, 262–268. 10.1038/mt.2008.244.

61. Rodríguez-Correa, E., Grünschläger, F., Nizharadze, T., Anstee, N., Al-Sabah, J., Kumpost, V., Sedlmeier, A., Li, C., Ball, M., Fotopoulou, F., et al. (2025). A kinetics-based model of hematopoiesis reveals extrinsic regulation of skewed lineage output from stem cells. bioRxiv, 2025.2002.2004.636388. 10.1101/2025.02.04.636388.

62. Belander Strålin, K., Carrelha, J., Winroth, A., Ziegenhain, C., Hagemann-Jensen, M., Kettyle, L.M., Hillen, A., Högstrand, K., Markljung, E., Grasso, F., et al. (2023). Platelet and myeloid lineage biases of transplanted single perinatal mouse hematopoietic stem cells. In Cell Res, pp. 883–886. 10.1038/s41422-023-00866-4.

63. Grover, A., Sanjuan-Pla, A., Thongjuea, S., Carrelha, J., Giustacchini, A., Gambardella, A., Macaulay, I., Mancini, E., Luis, T.C., Mead, A., et al. (2016). Single-cell RNA sequencing reveals molecular and functional platelet bias of aged haematopoietic stem cells. Nat Commun 7, 11075. 10.1038/ncomms11075.

64. Ross, J.B., Myers, L.M., Noh, J.J., Collins, M.M., Carmody, A.B., Messer, R.J., Dhuey, E., Hasenkrug, K.J., and Weissman, I.L. (2024). Depleting myeloid-biased haematopoietic stem cells rejuvenates aged immunity. Nature 628, 162–170. 10.1038/s41586-024-07238-x.

65. Wilkinson, A.C., Ishida, R., Kikuchi, M., Sudo, K., Morita, M., Crisostomo, R.V., Yamamoto, R., Loh, K.M., Nakamura, Y., Watanabe, M., et al. (2019). Long-term ex vivo haematopoietic-stem-cell expansion allows nonconditioned transplantation. Nature 571, 117–121. 10.1038/s41586-019-1244-x.

66. Fukushima, T., Tanaka, Y., Hamey, F.K., Chang, C.H., Oki, T., Asada, S., Hayashi, Y., Fujino, T., Yonezawa, T., Takeda, R., et al. (2019). Discrimination of Dormant and Active Hematopoietic Stem Cells by G(0) Marker Reveals Dormancy Regulation by Cytoplasmic Calcium. Cell Rep 29, 4144–4158.e4147. 10.1016/j.celrep.2019.11.061.

67. Davidsson, M., Diaz-Fernandez, P., Schwich, O.D., Torroba, M., Wang, G., and Björklund, T. (2016). A novel process of viral vector barcoding and library preparation enables high-diversity library generation and recombination-free paired-end sequencing. Sci Rep 6, 37563. 10.1038/srep37563.

68. Zhao, L., Liu, Z., Levy, S.F., and Wu, S. (2018). Bartender: a fast and accurate clustering algorithm to count barcode reads. Bioinformatics 34, 739–747. 10.1093/bioinformatics/btx655.

69. Becker, H.J., Ishida, R., Wilkinson, A.C., Kimura, T., Lee, M.S.J., Coban, C., Ota, Y., Tanaka, Y., Roskamp, M., Sano, T., et al. (2023). Controlling genetic heterogeneity in gene-edited hematopoietic stem cells by single-cell expansion. Cell Stem Cell 30, 987–1000.e1008. 10.1016/j.stem.2023.06.002.

70. Conant, D., Hsiau, T., Rossi, N., Oki, J., Maures, T., Waite, K., Yang, J., Joshi, S., Kelso, R., Holden, K., et al. (2022). Inference of CRISPR Edits from Sanger Trace Data. Crispr j 5, 123–130. 10.1089/crispr.2021.0113.

71. Love, M.I., Huber, W., and Anders, S. (2014). Moderated estimation of fold change and dispersion for RNA-seq data with DESeq2. Genome Biol 15, 550. 10.1186/s13059-014-0550-8.

72. Buenrostro, J.D., Giresi, P.G., Zaba, L.C., Chang, H.Y., and Greenleaf, W.J. (2013). Transposition of native chromatin for fast and sensitive epigenomic profiling of open chromatin, DNA-binding proteins and nucleosome position. Nat Methods 10, 1213–1218. 10.1038/nmeth.2688.

73. Feng, J., Liu, T., Qin, B., Zhang, Y., and Liu, X.S. (2012). Identifying ChIP-seq enrichment using MACS. Nat Protoc 7, 1728–1740. 10.1038/nprot.2012.101.

74. Ross-Innes, C.S., Stark, R., Teschendorff, A.E., Holmes, K.A., Ali, H.R., Dunning, M.J., Brown, G.D., Gojis, O., Ellis, I.O., Green, A.R., et al. (2012). Differential oestrogen receptor binding is associated with clinical outcome in breast cancer. Nature 481, 389–393. 10.1038/nature10730.

75. Shen, W., Le, S., Li, Y., and Hu, F. (2016). SeqKit: A Cross-Platform and Ultrafast Toolkit for FASTA/Q File Manipulation. PLoS One 11, e0163962. 10.1371/journal.pone.0163962.

76. Martin, M. (2011). Cutadapt removes adapter sequences from high-throughput sequencing reads. EMBnet. journal 17, 10–12.

77. Gu, Z., Eils, R., and Schlesner, M. (2016). Complex heatmaps reveal patterns and correlations in multidimensional genomic data. Bioinformatics 32, 2847–2849. 10.1093/bioinformatics/btw313.

78. Gu, Z. (2022). Complex heatmap visualization. Imeta 1, e43.

79. Needleman, S.B., and Wunsch, C.D. (1970). A general method applicable to the search for similarities in the amino acid sequence of two proteins. J Mol Biol 48, 443–453. 10.1016/0022-2836(70)90057-4.

80. Cock, P.J., Antao, T., Chang, J.T., Chapman, B.A., Cox, C.J., Dalke, A., Friedberg, I., Hamelryck, T., Kauff, F., Wilczynski, B., and de Hoon, M.J. (2009). Biopython: freely available Python tools for computational molecular biology and bioinformatics. Bioinformatics 25, 1422–1423. 10.1093/bioinformatics/btp163.

81. Sokal, R.R., and Michener, C.D. (1958). A statistical method for evaluating systematic relationships.

82. Hu, Y., and Smyth, G.K. (2009). ELDA: extreme limiting dilution analysis for comparing depleted and enriched populations in stem cell and other assays. J Immunol Methods 347, 70–78. 10.1016/j.jim.2009.06.008.

83. Zheng, G.X., Terry, J.M., Belgrader, P., Ryvkin, P., Bent, Z.W., Wilson, R., Ziraldo, S.B., Wheeler, T.D., McDermott, G.P., Zhu, J., et al. (2017). Massively parallel digital transcriptional profiling of single cells. Nat Commun 8, 14049. 10.1038/ncomms14049.

84. Hao, Y., Hao, S., Andersen-Nissen, E., Mauck, W.M, 3rd., Zheng, S., Butler, A., Lee, M.J., Wilk, A.J., Darby, C., Zager, M., et al. (2021). Integrated analysis of multimodal single-cell data. Cell 184, 3573–3587.e3529. 10.1016/j.cell.2021.04.048.

85. Korsunsky, I., Millard, N., Fan, J., Slowikowski, K., Zhang, F., Wei, K., Baglaenko, Y., Brenner, M., Loh, P.R., and Raychaudhuri, S. (2019). Fast, sensitive and accurate integration of single-cell data with Harmony. Nat Methods 16, 1289–1296. 10.1038/s41592-019-0619-0.

86. Hamey, F.K., and Göttgens, B. (2019). Machine learning predicts putative hematopoietic stem cells within large single-cell transcriptomics data sets. Exp Hematol 78, 11–20. 10.1016/j.exphem.2019.08.009.

87. Kuleshov, M.V., Jones, M.R., Rouillard, A.D., Fernandez, N.F., Duan, Q., Wang, Z., Koplev, S., Jenkins, S.L., Jagodnik, K.M., Lachmann, A., et al. (2016). Enrichr: a comprehensive gene set enrichment analysis web server 2016 update. Nucleic Acids Res 44, W90–97. 10.1093/nar/gkw377.

88. Saul, D., Kosinsky, R.L., Atkinson, E.J., Doolittle, M.L., Zhang, X., LeBrasseur, N.K., Pignolo, R.J., Robbins, P.D., Niedernhofer, L.J., Ikeno, Y., et al. (2022). A new gene set identifies senescent cells and predicts senescence-associated pathways across tissues. Nat Commun 13, 4827. 10.1038/s41467-022-32552-1.

89. Zhou, N., Peng, L., Luo, Q., Yin, T., Sun, H., Zhang, Y., Shi, X., Peng, X., Bao, J., Ning, Y., and Yuan, X. (2026). FerrDb V3: expanding the manually curated resource for regulators and disease associations from ferroptosis to regulated cell death. Nucleic Acids Res 54, D572–d582. 10.1093/nar/gkaf1119.

90. Gillespie, M., Jassal, B., Stephan, R., Milacic, M., Rothfels, K., Senff-Ribeiro, A., Griss, J., Sevilla, C., Matthews, L., Gong, C., et al. (2022). The reactome pathway knowledgebase 2022. Nucleic Acids Res 50, D687–d692. 10.1093/nar/gkab1028.

