## SupplementaryFigures for "Hematopoietic stem cells undergo bidirectional fate transitions *in vivo*"

### Supplemetal Figure 1

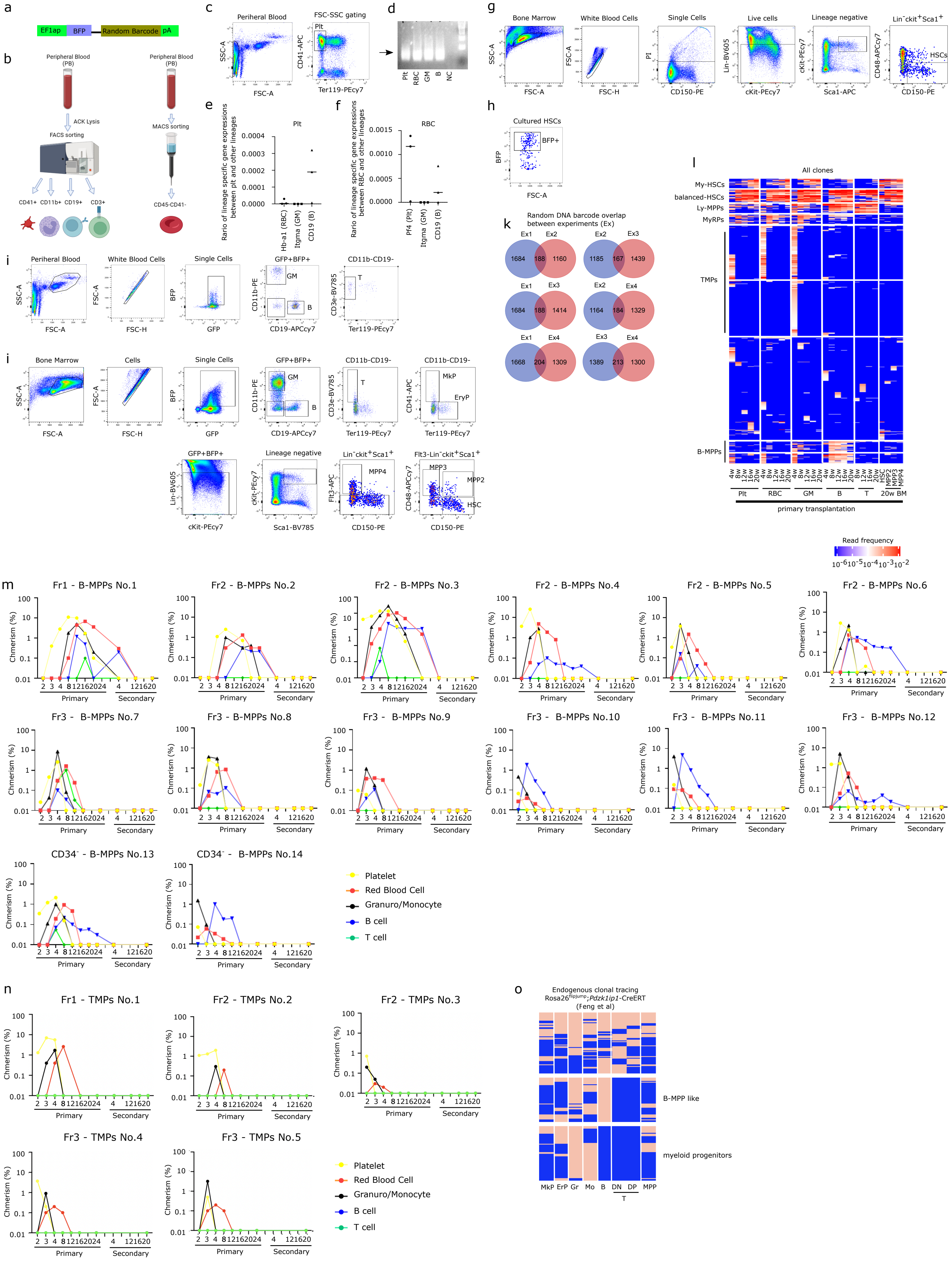

**Supplementary Figure 1 | Random DNA barcoding experiment with time course sampling.**

**a**, Schematic diagram of the barcoding vector. **b**, Schematic diagram of the purification methods for platelets and RBC. **c**, Gating strategy for PB platelets. **d**, Representative agarose gel electrophoresis profile of PCR products from mRNA of platelets and RBCs, and the genome of GM and B cells. **e**, Dot plot showing the expression of Hb-a1, Itgma, and Cd19 in the sorted CD41$^+$ fraction relative to their expression in RBC, GM, and B cells derived from the same amount of PB, respectively. **f**, Dot plot showing the expression of Pf4, Itgma, and Cd19 in the CD45-CD41- fraction relative to their expression in platelets, GM, and B cells derived from the same amount of PB, respectively (**e, f,** n = 3 from 3 independent experiments). **g**, Gating strategy for the BM immunophenotypic HSC fraction (CD150$^+$CD48$^-$Lin$^-$cKit$^+$Sca1$^+$). **h**, Representative FACS plot depicting the induction of random DNA barcode. **i**, Gating strategy for PB barcoded GM, B, and T cells. **j**, Gating strategy for BM barcoded MkP, ErP, GM, B, T, HSC, MPP2, MPP3, and MPP4 (HSC:CD150$^+$CD48$^-$Flt3$^-$Lin$^-$cKit$^+$Sca1$^+$, MPP2:CD150$^+$CD48$^+$Flt3$^-$Lin$^-$cKit$^+$Sca1$^+$, MPP3:CD150$^-$CD48$^+$Flt3$^-$Lin$^-$cKit$^+$Sca1$^+$, MPP4:Flt3$^+$Lin$^-$cKit$^+$Sca1$^+$). **k**, Venn diagram showing random DNA barcode overlap between experiments. **l**, Heatmap showing clones derived from the immunophenotypic HSC fraction, with rows representing unique random DNA barcodes and columns representing time points (4w, 8w, 12w, 16w, 20w) and cell types (plt, RBCs, GM, B, T, HSC, MPP2, MPP3 and MPP4). Read frequency is color-coded; clusters are annotated by output pattern (My-HSCs, balanced-HSCs, MyRPs, Ly-MPPs, B-MPPs, TMPs) (n = 6335 clones, 4 biological replicates from 2 independent experiments). M, Single-cell primary and secondary transplantation results showing the lineage output of B-MPPs. N, Single-cell primary and secondary transplantation results showing the lineage output of TMPs. **m, n**; The contribution of individual transplanted cells to different blood lineages is shown. Lineage colors are as follows: yellow, platelets; red, red blood cells; black, granulocytes/monocytes; blue, B cells; green, T cells. The vertical axis represents donor cell chimerism, and the horizontal axis represents weeks post-transplantation. (Fr1 = CD34$^-$CD150$^+$CD41$^-$cKit$^+$Sca1$^+$Lin$^-$, Fr2 = CD34$^-$CD150$^+$CD41$^+$cKit$^+$Sca1$^+$Lin$^-$, Fr3 = CD34$^-$CD150$^-$cKit$^+$Sca1$^+$Lin$^-$, CD34$^-$ = CD34$^-$cKit$^+$Sca1$^+$Lin$^-$). **o**, Barcode data from Rosa26$^{FlipJump}$; *Pdzk1ip1*-CreERT mice 6-10m after tamoxifen administration (Feng et al., 2024). Each raw represents a clone. Each column represents cell types listed bellow. Multipotent progenitors (MPP; Sca-1$^+$c-Kit$^+$CD48$^+$CD150$^-$), megakaryocyte progenitors (MkP; Sca-1$^-$c-Kit$^+$ CD150$^+$CD41$^+$), and erythroid progenitors (EryP; Sca-1$^-$c-Kit$^+$CD150$^+$CD41$^-$CD71$^+$) from lineage (Ter119, Gr-1, NK1.1, CD11b, B220 and TCRb)-negative BM fraction. Granulocytes (TCRβ$^-$B220$^-$CD11b$^+$Ly6C$^{low}$SSC$^{high\wedge}$), monocytes (TCRβ$^-$B220$^-$CD11b$^+$Ly6C$^{high}$), and B cells (CD11c$^-$B220$^+$SiglecH$^-$CD93$^+$) from total BM. Double-positive (DP; CD4$^+$CD8$^+$TCRβ$^{-/low}$) and double-negative (DN; CD4$^-$CD8$^-$CD25$^+$TCRβ$^-$) from thymus. The presence of a given lineage is indicated in red, whereas its absence is shown in blue. (n = 231 clones)

### Supplemetal Figure 2

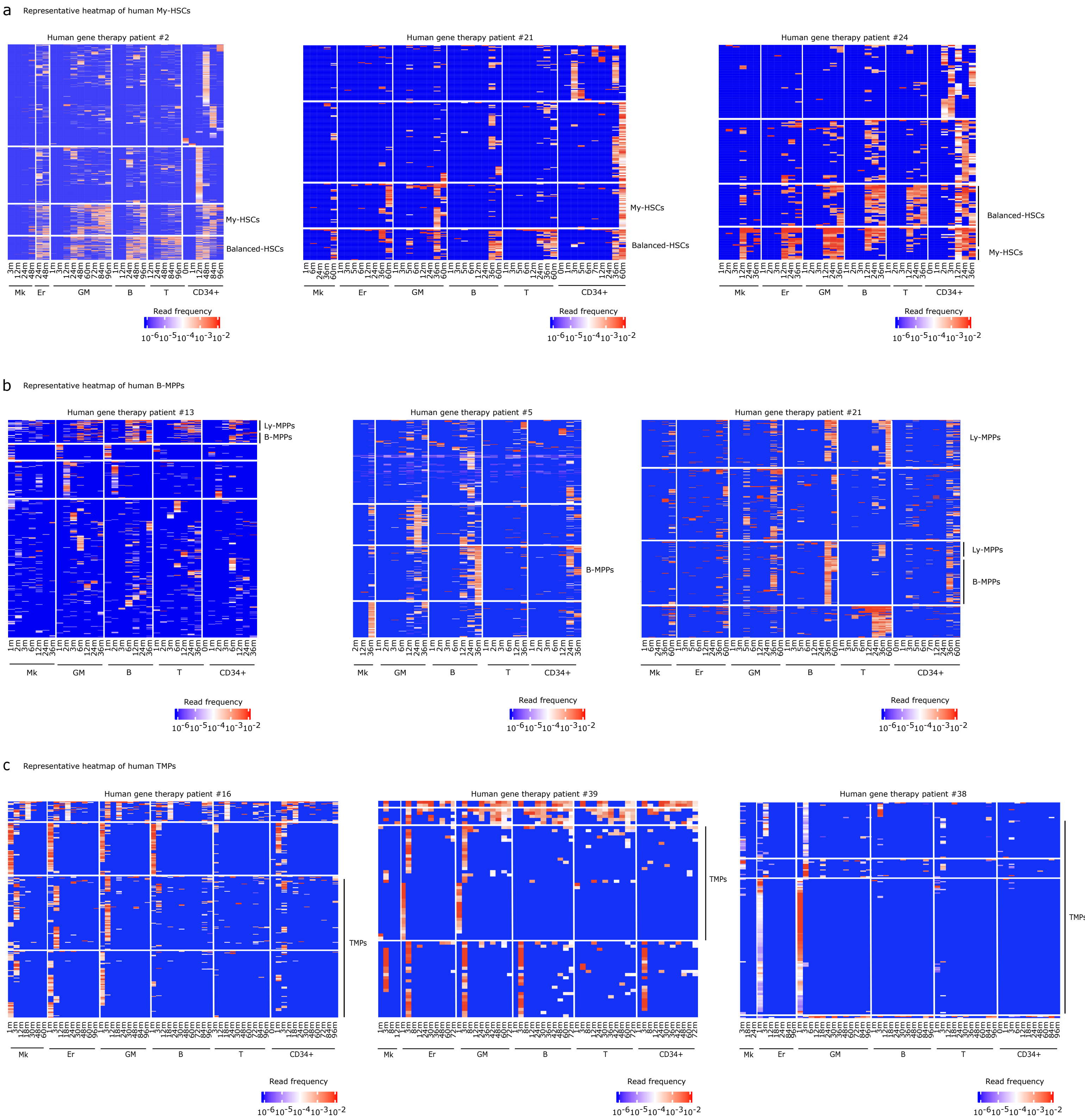

**Supplementary Figure 2 | Clonal tracing data from human gene therapy patients.**

**a,** Representative heatmap of clones reconstituting the BM CD34-positive fraction in gene therapy patients showing the My-HSCs (Carabria et al., 2024). **b**, Representative heatmap of clones in gene therapy patients showing the B-MPPs (Carabria et al., 2024). **c**, Representative heatmap of clones in gene therapy patients showing the TMPs (Carabria et al., 2024). **a-c**; Each rows representing unique lentiviral insertion sites and each columns representing time points (months after transplantation) and cell types (Mk:CD61$^{+}$, Erythrocyte (Er):Gr1$^{+}$, GM:CD13$^{+}$ or CD15$^{+}$, B:CD19$^{+}$, T:CD3$^{+}$ and CD34$^{+}$ cells). Colors indicate read frequency.

### Supplemetal Figure 3

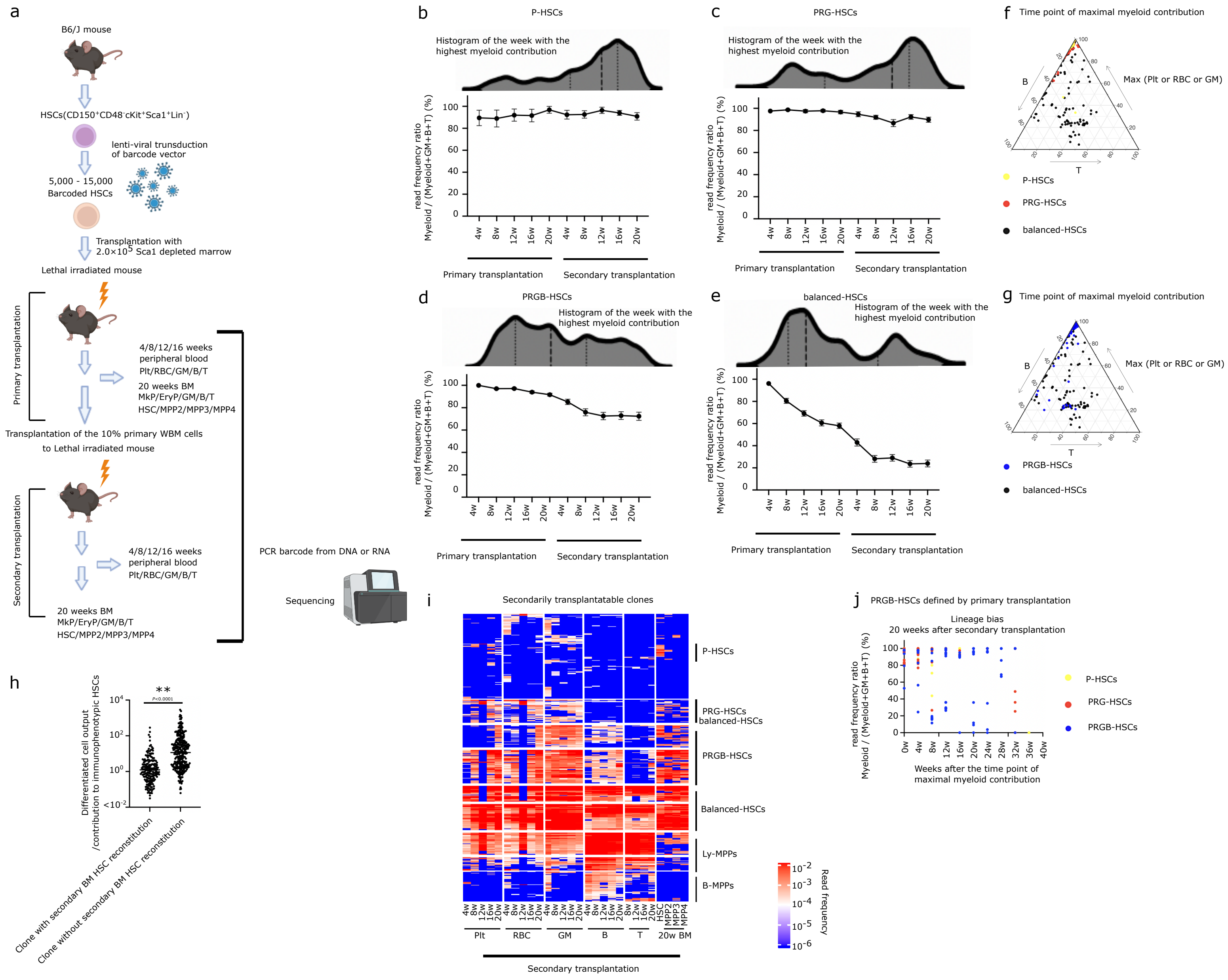

**Supplementary Figure 3 | Lineage tracking during secondary transplantation in a random DNA barcoding experiment**

**a**, Schematic diagram of the experimental setup for secondary transplantation. **b, c, d, e**, The line graph shows the Read frequency ratio of myeloid output Myeloid/(Myeloid+B+T) for **b**, P-HSCs (n = 63) **c**, PRG-HSCs (n = 95) **d**, PRGB-HSCs (n = 135) and **e**, balanced-HSCs (n = 140) over weeks after primary and secondary transplantation. Overlaid histograms indicate the distribution of the week at which each HSC exhibited the highest myeloid contribution. .**f, g,** Ternary plot of lineage output at the time point of maximal myeloid contribution from individual HSCs. Myeloid contribution was defined as the maximum among Plt, RBC, and GM lineages. **f**, Colors indicate HSC subtypes (P-HSCs, yellow (n = 81); PRG-HSCs, red (n = 135); balanced-HSCs, black (n= 140)). **g,** Colors indicate HSC subtypes (PRGB-HSCs, blue (n= 92); balanced-HSCs, black (n= 140)). **h**, Dot plot showing the ratio of differentiation versus self-renewal (highest read frequency of clones across all differentiated cells and all time points / read frequency of the 20w BM immunophenotypic HSC fraction) of clones with or without the capacity to reconstitute BM immunophenotypic HSCs 20weeks after secondary transplantation (n = 185 and 300 respectively). **i**, Heatmap of clones reconstituting secondary transplantation with rows representing unique DNA barcodes and columns representing time points (4w, 8w, 12w, 16w, 20w) and cell types (Plt, RBC, GM, B, T, HSC, MPP2, MPP3 and MPP4) (HSC:$CD150^{+}CD48^{-}Flt3^{-}Lin^{-}cKit^{+}Sca1^{+}$, MPP2:$CD150^{+}CD48^{+}Flt3^{-}Lin^{-}cKit^{+}Sca1^{+}$, MPP3:$CD150^{-}CD48^{+}Flt3^{-}Lin^{-}cKit^{+}Sca1^{+}$, MPP4:$Flt3^{+}Lin^{-}cKit^{+}Sca1^{+}$). Colors indicate read frequency. Clones are annotated based on output patterns (P-HSCs, PRG-HSCs, PRGB-HSCs, balanced-HSCs, Ly-MPPs and B-MPPs) (n = 455 clones). **j,** Scatter plot of PRGB-HSCs defined by primary transplantation. The y-axis indicates the read frequency ratio of myeloid lineages Myeloid/(Myeloid+B+T) measured 20 weeks after secondary transplantation. The x-axis represents the weeks after the time point of maximal myeloid contribution. (n = 92). (**b-j**; 3 biological replicates from 2 independent experiments).

### Supplemental Figure 4

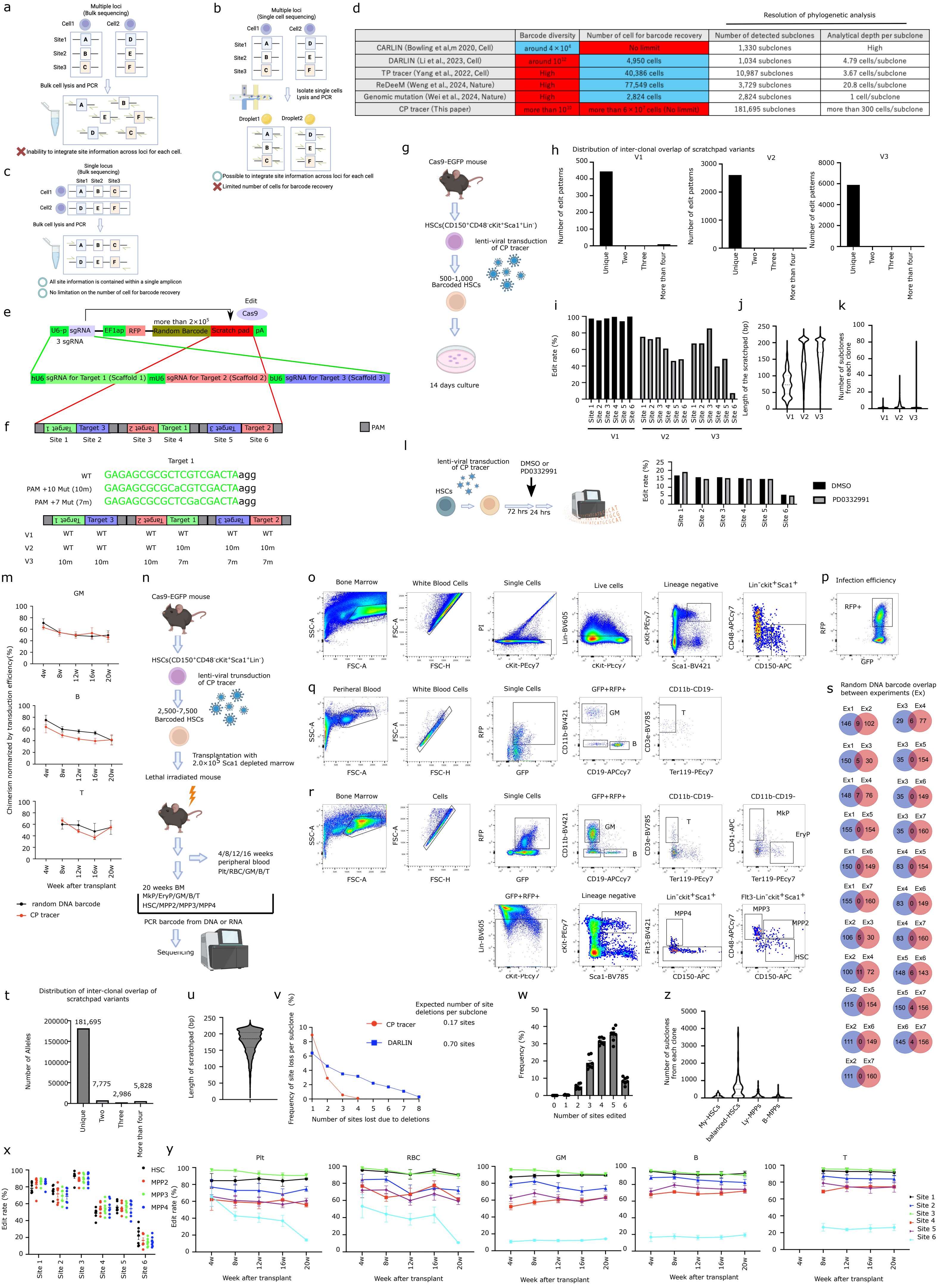


d

| | Barcode diversity | Number of cell for barcode recovery | Number of detected subclones | Analytical depth per subclone |
|---|---|---|---|---|
| | | | Resolution of phylogenetic analysis | |
| CARLIN (Bowling et al,m 2020, Cell) | around $4\times10^4$ | No limmit | 1,330 subclones | High |
| DARLIN (Li et al., 2023, Cell) | around $10^{12}$ | 4,950 cells | 1,034 subclones | 4.79 cells/subclone |
| TP tracer (Yang et al., 2022, Cell) | High | 40,386 cells | 10,987 subclones | 3.67 cells/subclone |
| ReDeeM (Weng et al., 2024, Nature) | High | 77,549 cells | 3,729 subclones | 20.8 cells/subclone |
| Genomic mutation (Wei et al., 2024, Nature) | High | 2,824 cells | 2,824 subclones | 1 cell/subclone |
| CP tracer (This paper) | more than $10^{10}$ | more than $6\times10^7$ cells (No limmit) | 181,695 subclones | more than 300 cells/subclone |

**Supplementary Figure 4 | Development of clonal phylogenic tracer "CP-tracer"**

**a,** The schematic illustrates that bulk cell lysis, PCR, and sequencing of cells carrying barcodes distributed across multiple genomic loci prevents reconstruction of per-cell combinations of site-specific barcode information**. b,** The schematic illustrates that single-cell isolation followed by lysis, PCR, and sequencing enables reconstruction of per-cell combinations of site-specific barcode information across multiple loci. However, current single-cell technologies limit the number of cells available for barcode recovery to up to ~$10^5$ cells, which is insufficient to evaluate lineage distributions for each of more than $10^5$ branches. **c,** The schematic illustrates that, when barcodes are confined to a single genomic locus, bulk cell lysis followed by PCR and sequencing enables analysis of which combinations of multiple site-specific barcode information were present in individual cells, because all site information is contained within a single amplicon. In addition, bulk methods impose no limitation on the number of cells available for barcode recovery and are therefore sufficient to evaluate lineage distributions for each of more than $10^5$ branches. **d**, Table showing the barcode diversity, number of cells for barcode recovery, number of detected subclones, and subclone information content in this and previous studies. The red color highlights the advantages, while the blue color highlights limitations of the method. **e**, Schematic diagram of the structure of the CP tracer. **f**, Schematic diagram of the structure of the scratchpad. **g**, Schematic diagram of the experimental setup for validation of the CP tracer in vitro. **h**, Bar graph showing the number of scratchpad editing patterns shared among clones. **i**, Bar graph showing the edit rate of each site for each version of the CP tracer. **j**, Violin plot showing the length of the scratchpad after editing. **k**, Violin plot showing the number of subclones defined by scratchpad for each clone. (**h-k**; V1: n=339, V2: n=575, V3: n=1327 from 1 experiment). **l**, Bar graph showing the edit rate of each site for each version of the CP tracer. **m,** Chimerism of barcode transduced cells normalized by lentiviral transduction efficiency. HSCs derived from Cas9-EGFP mice received cells transduced with either the CP tracer system (red) or a random DNA barcode vector carrying sgRNA (black) via lentivirus. The y-axis shows chimerism of transduced cells divided by transduction efficiency, and the x-axis indicates weeks after transplantation (CP tracer; n = 7, random DNA barcode; n = 7). **n**, Schematic diagram of the experimental setup for phylogenetic tracing of HSCs. **o**, Gating strategy for the BM immunophenotypic HSC fraction ($CD150^{+}CD48^{-}Lin^{-}cKit^{+}Sca1^{+}$). **p**, Representative FACS plot depicting the induction of CP tracer. **q**, Gating strategy for PB barcoded GM, B, and T cells. **r**, Gating strategy for BM barcoded MkP, ErP, GM, B, T, HSC, MPP2, MPP3, and MPP4 (HSC:$CD150^{+}CD48^{-}Flt3^{-}Lin^{-}cKit^{+}Sca1^{+}$, MPP2:$CD150^{+}CD48^{+}Flt3^{-}Lin^{-}cKit^{+}Sca1^{+}$, MPP3:$CD150^{-}CD48^{+}Flt3^{-}Lin^{-}cKit^{+}Sca1^{+}$, MPP4:$Flt3^{+}Lin^{-}cKit^{+}Sca1^{+}$). **s**, Venn diagram showing random DNA barcode of CP tracer overlap between experiments. **t**, Bar graph showing the number of scratchpad editing patterns overlaped among all clones (n = 871 clones). **u**, Violin plot showing the length of the scratchpad after editing in balanced-HSCs (n = 181,695 subclones). **v**, Distribution of the number of sites lost due to deletions (x-axis) versus the frequency of deletion events per subclone (y-axis) for CP tracer and DARLIN. The expected number of deleted sites per subclone is also indicated. **w**, Bar graph showing the frequency of the number of sites of scratchpad edited in each subclone derived from balanced-HSCs (n = 58760 subclones from biological replicates). **x**, Dot plot showing the edit rate of each site in 20w BM immunophenotypic HSC, MPP2, MPP3 and MPP4 from balanced-HSCs (HSC: n = 1425; MPP2: n = 1785; MPP3: n = 2460; MPP4: n = 2687). **y**, Dot plots showing the edit rate of each site in PB Plt, RBC, GM, B, and T cells. The X-axis represents weeks after transplantation **z**, Violin plot showing the number of subclones from each clone (My-HSCs: n = 178; balanced-HSCs: n = 168; Ly-MPPs: n = 228; B-MPPs: n = 273 clones). (**T-Z**: 7 biological replicates from 4 independent experiments).

### Supplemental Figure 5

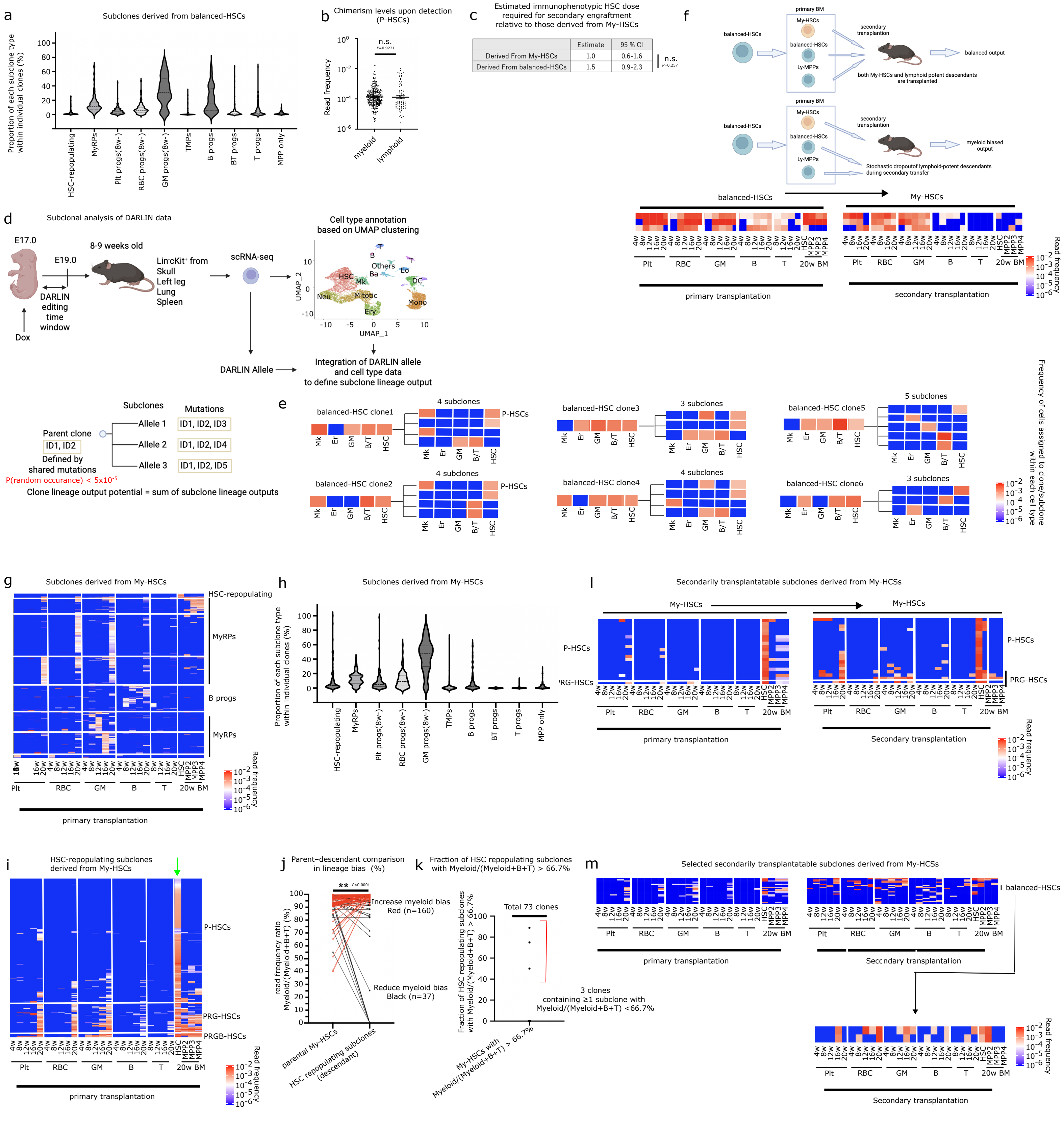

**Supplementary Figure 5 | Subclone analysis of each cell types.**

**a**, Violin plot showing the proportion of each subclone type in balanced-HSCs (n = 123 clones). **b**, Dot plot showing read frequencies of myeloid (maximum of Plt, RBC, and GM) and lymphoid (maximum of B and T) outputs of P-HSCs upon detection (myeloid; n = 321, lymphoid; n = 51).**c**, Table showing the estimated immunophenotypic HSC dose required to achieve secondary bone marrow reconstitution of immunophenotypic HSCs with a 63.2% probability across My-HSCs derived from balanced-HSCs and My-HSCs, relative to those derived from My-HSCs, calculated by ELDA. **d**, Schematic overview of the DARLIN experimental design and UMAP-based cell type annotation (upper). Schematic overview of the method for reconstructing clone–subclone relationships (bottom). **e**, Heatmap showing the lineage output of three Balanced-HSC clones identified by DARLIN that possess descendants; the left panel shows clonal output, and the right panel shows the output of descendant subclones. The frequency of cells harboring barcodes in each lineage is represented by a color scale (bottom). GM is defined as the sum of Neutrophils, Monocytes, Eosinophils, Basophils, and dendritic cells. B/T is defined as the sum of B and T cells. **d,e**, DARLIN data were obtained from Li et al.,2023. **f**, Schematic illustration showing a possible scenario in which My-HSCs are observed after secondary transplantation of balanced-HSC–derived grafts. Among the descendants of balanced-HSCs, balanced-HSCs and lymphoid progenitors may drop out during secondary transplantation by chance, whereas My-HSCs may coincidentally engraft and subsequently be detected (upper). Heatmap of clone which transition from balanced-HSCs to My-HSCs after secondarily transplant (n = 3) (bottom). **g**, Heatmap of subclones derived from My-HSCs, reconstituting more than two time points or cell types (Plt, RBC, GM, B, T, HSC, MPP2, MPP3 and MPP4). Clusters are annotated by output pattern (Self-renewing, MyRPs, B progs) (n = 2720 subclones). **h**, Violin plot showing the proportion of each subclone type in My-HSCs (n = 106 clones). The average and SEM are shown. **i**, Heatmap showing subclones derived from My-HSCs reconstituting 20w BM immunophenotypic HSCs (n = 412 subclones). The lineage output of each subclone was compared to the average outputs of clusters (P-HSCs, PRG-HSCs, PRGB-HSCs, and balanced-HSCs) in the reference heatmap (Fig.2a). Each subclone was then assigned to the cluster it was closest to. The column indicated by the green arrow represents the contribution to BM HSCs at 20 weeks after transplantation. **j**, Paired comparison of lineage bias between parental My-HSCs and their descendant HSC-repopulating subclones. Lineage bias was defined as the percentage contribution of Myeloid (max of Plt, PrB, and GM) relative to total output (Myeloid+B+T). Red (n = 197 subclones), increased myeloid bias; blue (n = 160 subclones), decreased myeloid bias; black (n = 37 subclones). **k**, Fraction of HSC-repopulating subclones with Myeloid/(Myeloid+B+T) > 66.7% among clones derived from My-HSCs with parental Myeloid/(Myeloid+B+T) > 66.7%. Each dot represents an individual clone. **l,** Heatmap of subclones reconstituting 20w BM immunophenotypic HSCs and secondary transplantation derived from My-HSCs (n = 20 subclones). **m,** Heatmap of subclones with lymphoid production during secondary transplantation derived from My-HSCs (n = 24 subclones). (**f, g, i, l,m**) Each row represents a unique random DNA barcode-scratchpad pair. Each column represents time points (4w, 8w, 12w, 16w, 20w after transplantation) and cell types (Plt, RBC, GM, B, T, HSC, MPP2, MPP3 and MPP4). Read frequency is represented by color-code scale. (**a,b, and g-k**, 7 biological replicates from 4 independent experiments, **c**,**l,m,** 2 biological replicates).

### Supplemental Figure 6

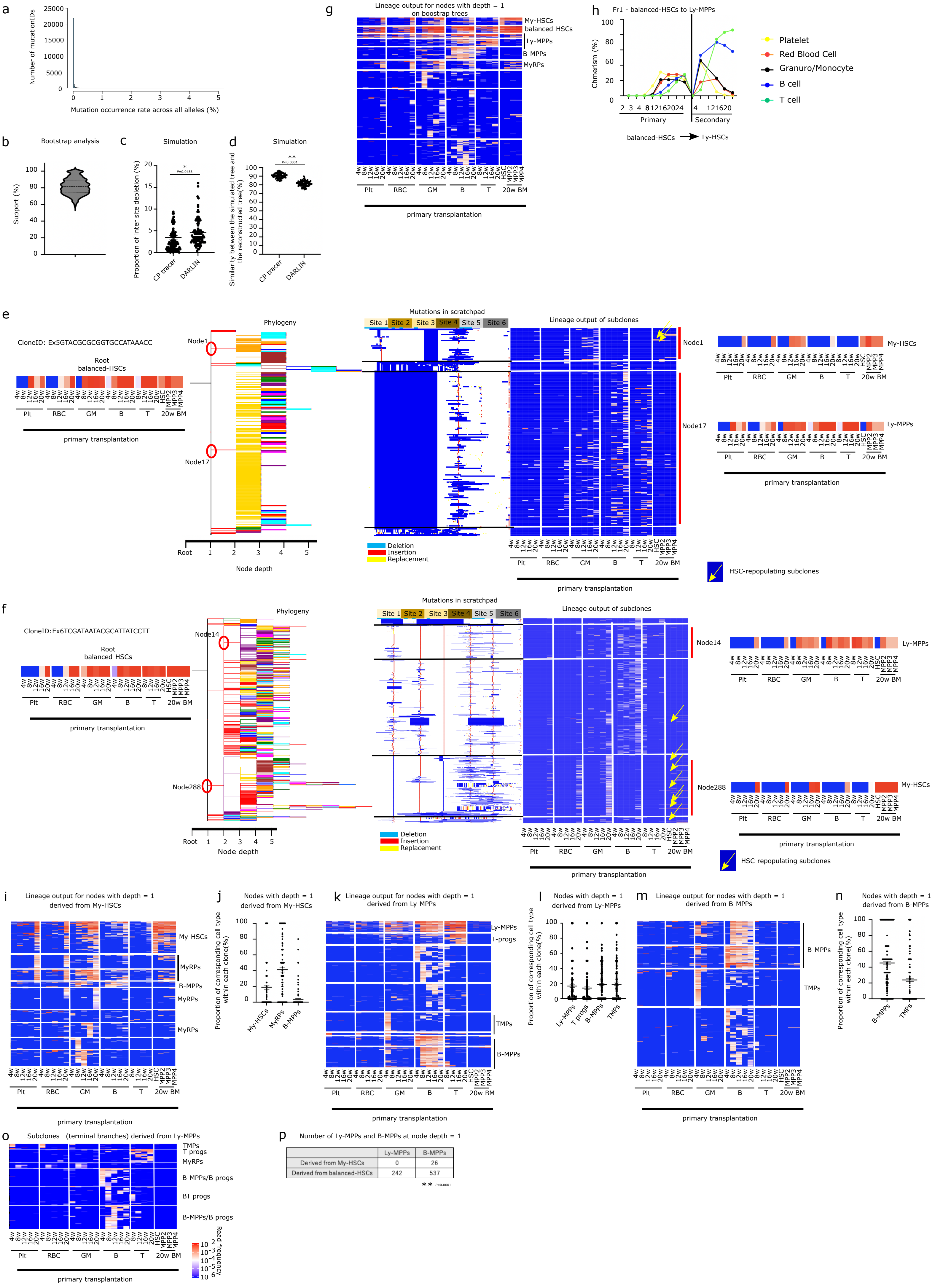


| | Ly-MPPs | B-MPPs |
|---|---|---|
| Derived from My-HSCs | 0 | 26 |
| Derived from balanced-HSCs | 242 | 537 |

** P<0.0001

**Supplementary Figure 6 | Phylogenetic tree analysis.**

**a**, Histogram showing the occurrence frequencies of mutations across alleles. **b**, Violin plot showing the distribution of bootstrap support values for the branches (Number of bootstraps = 100). **c**,Dot plot showing proportion of inter-site depletion in the simulation (n = 100 simulations). **d**, Dot plot showing similarity between the simulated tree and the reconstructed tree (n = 100 simulations). **e-f,** Tree reconstruction of two representative balanced-HSC clones. From left to right: clonal output (1st panel), phylogeny of the clone, with branches originating from the same node shown in the same color (2nd panel), scratchpad editing patterns (3rd panel), heatmap of sub-clonal output at the terminal leaves (4th panel), and heatmap of the nodes highlighted with red circles in the phylogeny (5th panel). In the scratchpad editing panel, rows represent unique alleles (subclones) and columns represent target sites, with edits color-coded as follows: deletion (blue), different base (yellow), and insertion (red). In the sub-clonal output panels, columns indicate time points and cell types (Plt, RBC, GM, B, T, HSC, MPP2, MPP3 and MPP4), and color intensity represents read frequency. **g**, Heatmap of nodes at branch depth 1 in the phylogeny derived from balanced-HSCs, generated from pseudo-datasets created by randomly omitting or duplicating portions of the scratchpad sequence information (Number of bootstraps = 100). **h**, representative single-cell primary and secondary transplantation results showing balanced-HSCs transition toward Ly-MPPs. The contribution of individual transplanted cells to different blood lineages is shown. Lineage colors are as follows: yellow, platelets; red, red blood cells; black, granulocytes/monocytes; blue, B cells; green, T cells. The vertical axis represents donor cell chimerism, and the horizontal axis represents weeks post-transplantation. (Fr1 = $CD34^{-}CD150^{+}CD41^{-}$ $cKit^{+}Sca1^{+}Lin^{-}$). **i,** Heatmap of nodes at branch depth 1 in the phylogeny derived from My-HSCs (n = 516 nodes). **j,** Dot plot showing the proportion of each sub-type of nodes at branch depth 1 in the phylogeny derived from My-HSCs. **k,** Heatmap of nodes at branch depth 1 in the phylogeny derived from Ly-MPPs (n = 1961 nodes). **l,** Dot plot showing the proportion of each sub-type of nodes at branch depth 1 in the phylogeny derived from Ly-MPPs. **m,** Heatmap of nodes at branch depth 1 in the phylogeny derived from B-MPPs (n = 1127 nodes). **n,** Dot plot showing the proportion of each sub-type of nodes at branch depth 1 in the phylogeny derived from B-MPPs.**o**, Heatmap of subclones derived from Ly-MPPs, reconstituting more than three time points or cell types (Plt, RBC, GM, B, T, HSC, MPP2, MPP3 and MPP4). Clusters are annotated by output pattern (MyRPs, T progs, B-MPPs/B progs, BT progs), **p**, Table showing the numbers of B-MPPs and Ly-MPPs identified among first-generation descendant nodes (node depth = 1) derived from balanced-HSC and My-HSC clones. *P* values were calculated by Fisher's exact test. (**a,b,g,i-p**, 7 biological replicates from 4 independent experiments).

### Supplemental Figure 7

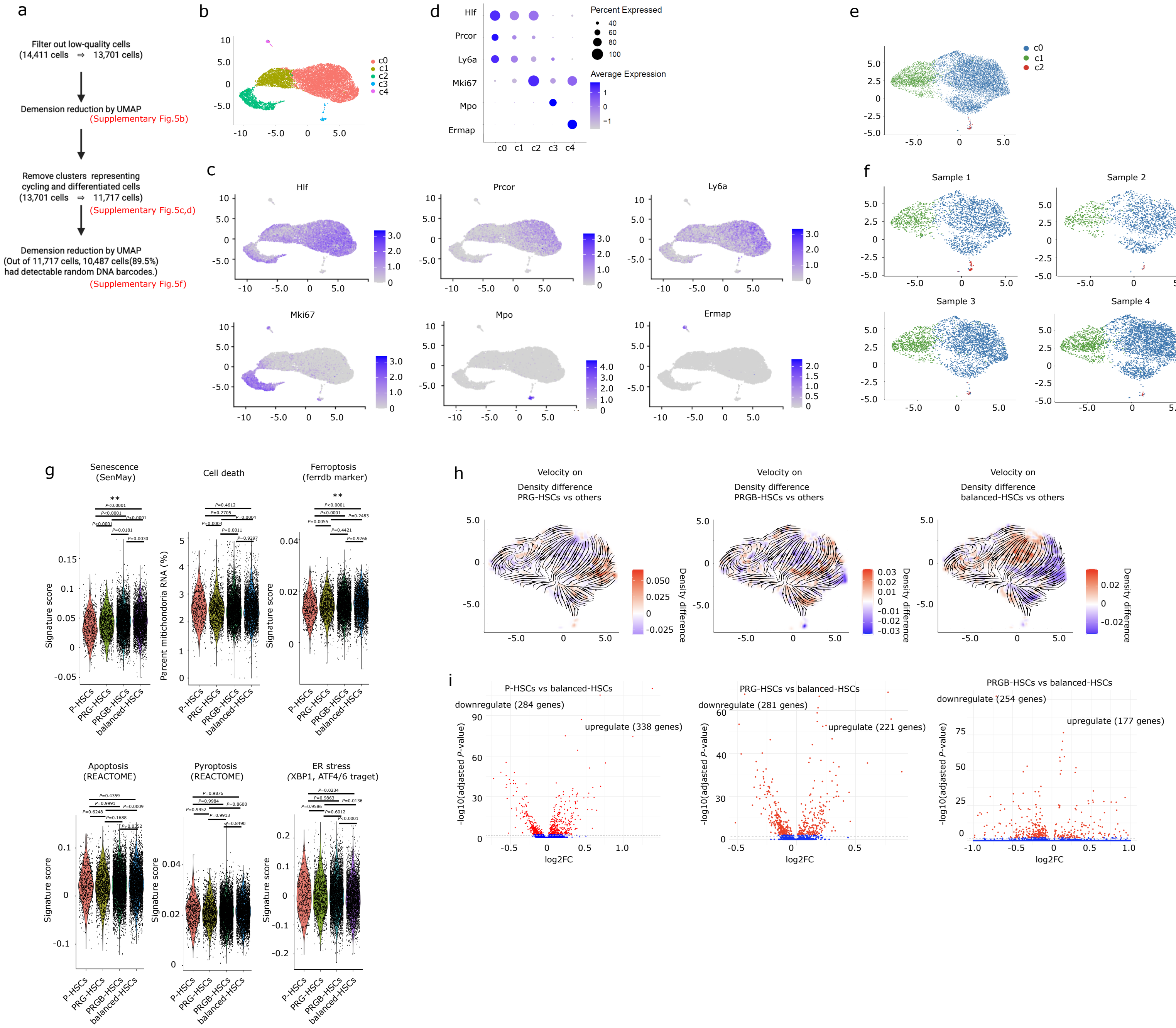

**Supplementary Figure 7 | Molecular analysis of each HSC subsets by scRNA-seq.**

**a,** Schematic illustrating the quality control workflow. **b**, Uniform manifold approximation and projection (UMAP) of cells from primary 20w BM immunophenotypic HSC fraction. **c**, UMAP of cells from primary 20w BM immunophenotypic HSC fraction, with each cell color-coded according to the marker gene expression (**c-e**, n=13701). **d,** Dot plots showing marker gene expression for each cluster. Dot size represents the percentage of cells expressing the marker, and the color represents the average expression level. **e**, UMAP of c0 and c1 cells from Figure S7B. 5c (n = 11,717). **f**, Distribution of cells on the UMAP for each sample. **g**, Violin plots showing expression scores of gene sets of senescence- and cell death–associated marker genes, along with the proportion of mitochondrial RNA (mtRNA) as an indicator of cell death. **h**, Color-coded density difference plots on UMAP for PRG-HSCs (left), PRGB-HSCs (middle) and balanced-HSCs (right), with arrows indicating inferred transcriptional dynamics and potential future states of cells. **i,** Volcano plot showing differentially expressed genes between P-HSCs vs. balanced-HSCs (left), PRG-HSCs vs. balanced-HSCs (middle) and PRG-HSCs vs. balanced-HSCs (right). Red dots represent differentially expressed genes (adjusted *P* value ≤ 0.01; P-HSCs: n = 631, PRG-HSCs: n = 1282, PRG-HSCs: n = 4958, balanced-HSCs: n = 3616).

### Supplemental Figure 8

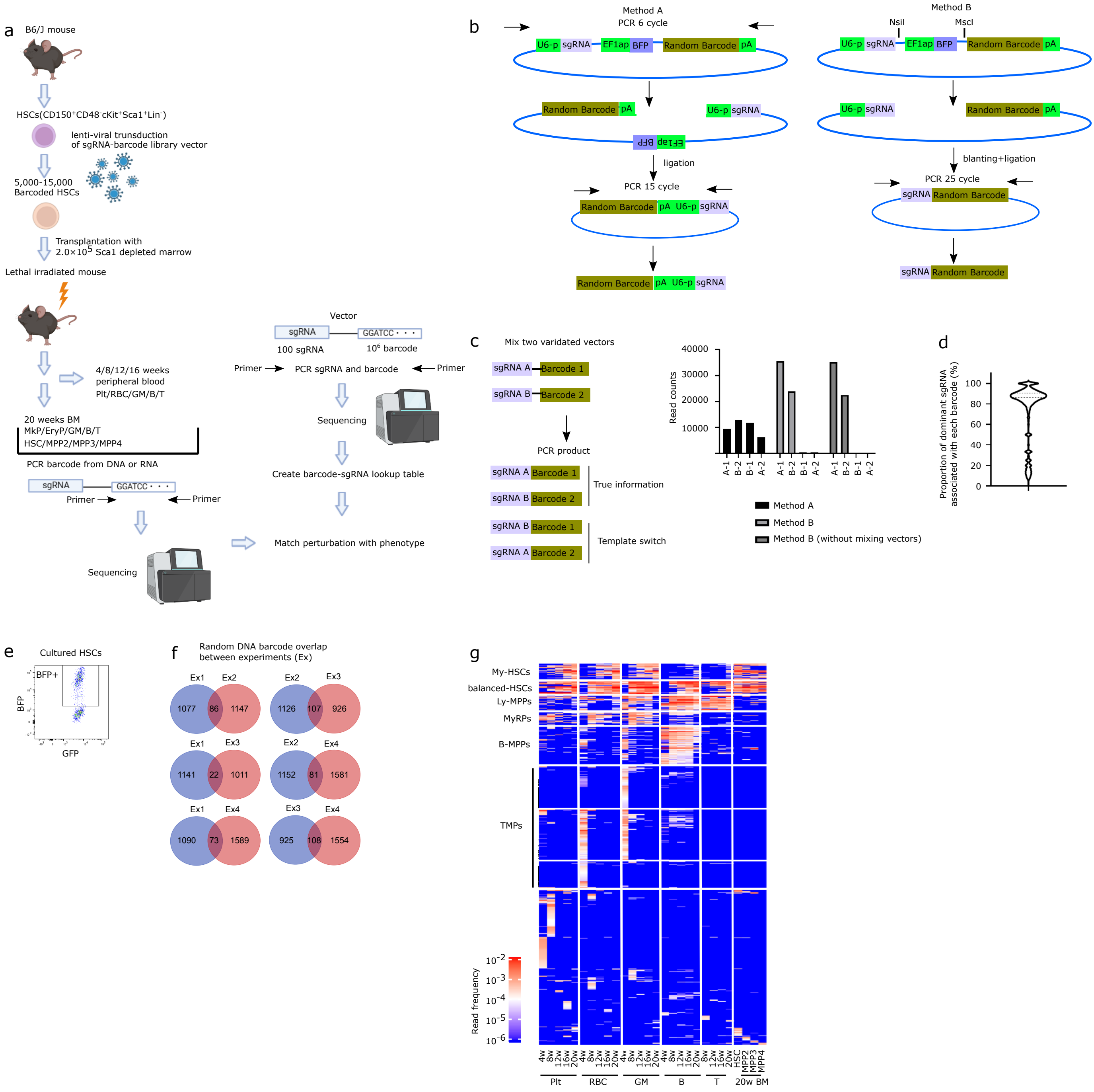

**Supplementary Figure 8 | Design of single cell level perturbation screening.**

**a**, Schematic diagram of the experimental setup and strategy for integrating sgRNA and random DNA barcode information. **b**, Schematic diagram illustrating the method for integrating sgRNA and random DNA barcode information. **c,** Schematic diagram of the strategy for validating the method depicted in Extended Data Fig. 6b (left). Bar graph showing the read count for sgRNA and barcode pairs (right). **d**, Violin plot showing the proportion of read counts of the dominant sgRNA within each barcode. **e**, Representative FACS plot depicting the induction of vector containing both random DNA barcode and sgRNA library. **f,** Venn diagram illustrating the overlap of random DNA barcodes between experiments. **g**, Heatmap showing clones derived from the immunophenotypic HSC fraction. Each row represents a unique random DNA barcode, and each column represents time points (4w, 8w, 12w, 16w, 20w after transplantation) and cell types (Plt, RBCs, GM, B, T, HSC, MPP2, MPP3 and MPP4) (HSC:$CD150^{+}CD48^{-}Flt3^{-}Lin^{-}cKit^{+}Sca1^{+}$, MPP2:$CD150^{+}CD48^{+}Flt3^{-}Lin^{-}cKit^{+}Sca1^{+}$, MPP3:$CD150^{-}CD48^{+}Flt3^{-}Lin^{-}cKit^{+}Sca1^{+}$, MPP4:$Flt3^{+}Lin^{-}cKit^{+}Sca1^{+}$). Read frequency is color-coded. Clusters are annotated by output patterns (My-HSCs, balanced-HSCs, MyRPs, Ly-MPPs, B-MPPs, TMPs) (n = 11,901 clones from 4 biological replicates across 2 independent experiments).

### Supplemental Figure 9

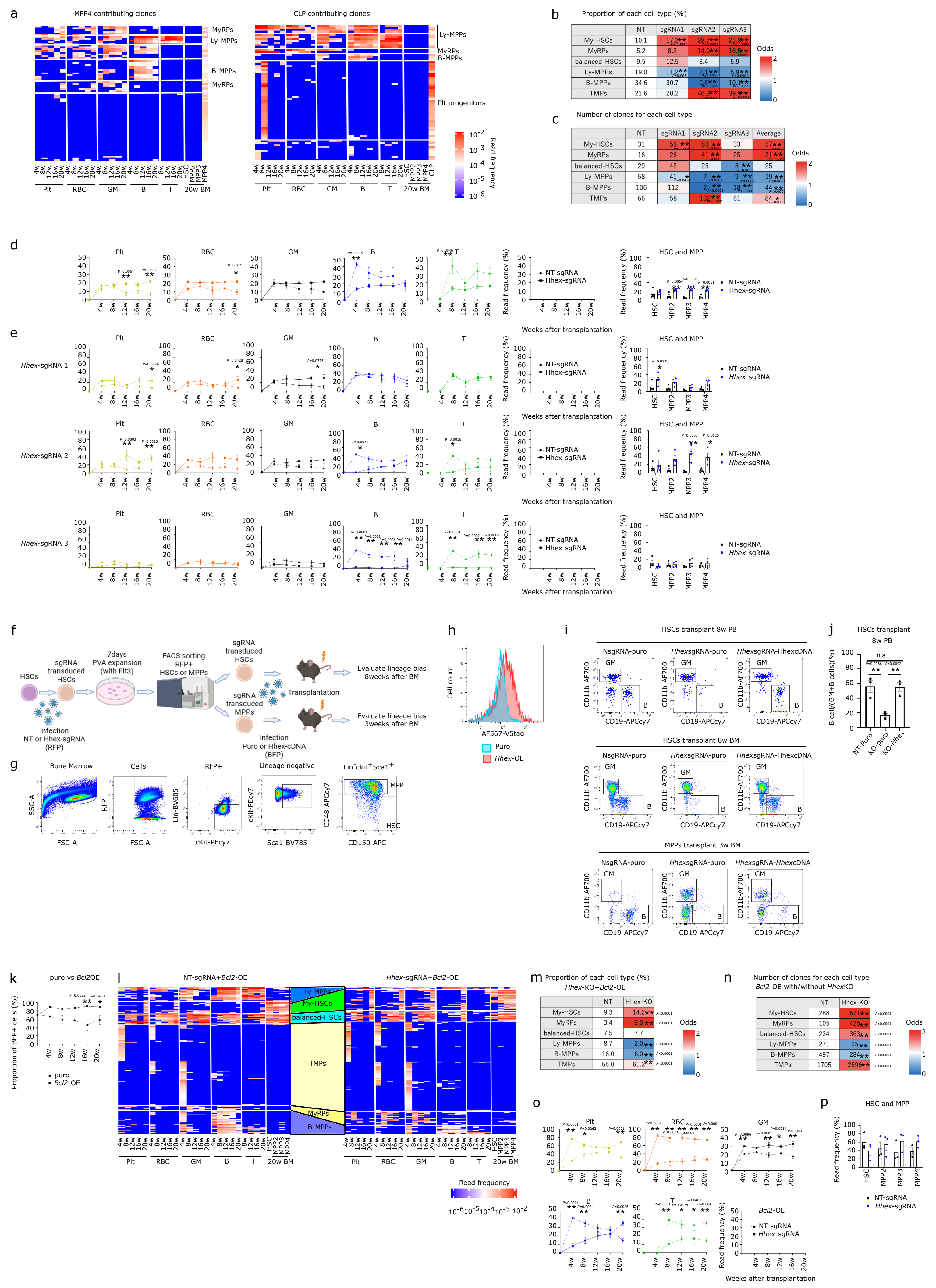


**b** Proportion of each cell type (%)

| | NT | sgRNA1 | sgRNA2 | sgRNA3 |
|---|---|---|---|---|
| My-HSCs | 10.1 | 17.2** | 28.7** | 21.8** |
| MyRPs | 5.2 | 8.2 | 14.0** | 16.5** |
| balanced-HSCs | 9.5 | 12.5 | 8.4 | 5.9 |
| Ly-MPPs | 19.0 | 11.2** | 2.1** | 5.9** |
| B-MPPs | 34.6 | 30.7 | 0.6** | 10.1** |
| TMPs | 21.6 | 20.2 | 46.3** | 39.9** |

**c** Number of clones for each cell type

| | NT | sgRNA1 | sgRNA2 | sgRNA3 | Average |
|---|---|---|---|---|---|
| My-HSCs | 31 | 56** | 83** | 33 | 57** |
| MyRPs | 16 | 26 | 41** | 25 | 31** |
| balanced-HSCs | 29 | 42 | 25 | 8** | 25 |
| Ly-MPPs | 58 | 41* | 7** | 9** | 19** |
| B-MPPs | 106 | 112 | 2** | 18** | 44** |
| TMPs | 66 | 58 | 132** | 61 | 84* |

**m** Proportion of each cell type (%), Hhex-KO+Bcl2-OE

| | NT | Hhex-KO |
|---|---|---|
| My-HSCs | 9.3 | 14.2** |
| MyRPs | 3.4 | 9.0** |
| balanced-HSCs | 7.5 | 7.7 |
| Ly-MPPs | 8.7 | 2.0** |
| B-MPPs | 16.0 | 6.0** |
| TMPs | 55.0 | 61.2** |

**n** Number of clones for each cell type, Bcl2-OE with/without HhexKO

| | NT | Hhex-KO |
|---|---|---|
| My-HSCs | 288 | 671** |
| MyRPs | 105 | 425** |
| balanced-HSCs | 234 | 363** |
| Ly-MPPs | 271 | 95** |
| B-MPPs | 497 | 284** |
| TMPs | 1705 | 2896** |

**Supplementary Figure 9 | Phenotype of *Hhex* KO.**

**a**, Heatmap showing clones contributing to immunophenotypic MPP4 (left) and CLP (right) (MPP4: n=78, 4 biological replicates from 2 independent experiments; CLP: n=66, 3 biological replicates from 3 independent experiments). **b**, Table showing the proportion of each cell type in NT-sgRNA and *Hhex*-sgRNA1, 2, and 3. **c**, Table showing the normalized number of clones of each cell type in NT-sgRNA and *Hhex*-sgRNA1, 2, and 3. **d,e**, Pseudo-bulk analysis of the read frequencies of clones within NT-sgRNA and *Hhex*-sgRNA groups. The x-axis represents weeks after transplantation. (**b-e**, NT-sgRNA; n=306, *Hhex*-sgRNA1; n=401, *Hhex*-sgRNA2; n=335, *Hhex*-sgRNA3; n=188 from 4 biological replicates from 2 independent experiments). **f**, Schematic diagram of the experimental setting for adding back *Hhex* to *Hhex*-KO HSCs and MPPs derived from *Hhex*-KO HSCs. **g**, Representative FACS plot of immunophenotypic HSCs or MPPs after PVA culture. **h**, Representative FACS plot of V5-*Hhex* expression. **i**, Representative FACS plot of PB B cells and GM, 8 weeks after immunophenotypic HSC transplantation (upper), BM B cells and GM, 8 weeks after immunophenotypic HSC transplantation (middle) and BM B cells and GM, 3 weeks after immunophenotypic MPP transplantation (bottom). **j**, The ratio of peripheral blood B cells/(GM + B cells) 8 weeks after the transplantation of immunophenotypic HSCs. **k,** Proportion of $BFP^{+}$ cells in PB mononuclear cells in puro or *Bcl2* OE groups after transplantation showing higher competitive advantage in *Bcl2* OE groups (puro: n=4; *Bcl2* OE: n=3). **l**, Heatmap showing clones from NT-sgRNA-*Bcl2*-OE (left) and *Hhex*-sgRNA-*Bcl2*-OE (right) groups. Each row represents a unique random DNA barcode, and each column represents time points (4w, 8w, 12w, 16w, and 20w after transplantation) and cell types (Plt, RBC, GM, B, T, ). Read frequency is represented by a color-coded scale. Each cluster was annotated based on the output pattern (My-HSCs, balanced-HSCs, MyRPs, Ly-MPPs, B-MPPs, TMPs). **m**, Table showing the proportion of each cell type in NT-sgRNA-*Bcl2*-OE and *Hhex*-sgRNA-*Bcl2*-OE groups. **n**, Table showing the normalized number of each cell type in NT-sgRNA-*Bcl2*-OE and *Hhex*-sgRNA-*Bcl2*-OE groups. **o,** Pseudo-bulk analysis of the read frequencies of clones within NT-sgRNA-*Bcl2*-OE and *Hhex*-sgRNA-*Bcl2*-OE groups in differentiated cells. **p**, Pseudo-bulk analysis of the read frequencies of clones within NT-sgRNA-*Bcl2*-OE and *Hhex*-sgRNA-*Bcl2*-OE groups in 20w BM (**l-p**, NT-sgRNA-*Bcl2*-OE; n=3100, NT-sgRNA-*Bcl2*-OE; n=4743 from 3 biological replicates). **P*<0.05; ***P*<0.01. **b**, **c**, **m, n,** *P* values were calculated by Fisher's exact test. **d,e, j, o,p,** *P* values were calculated by one-way or two-way ANOVA followed by Tukey's multiple comparison test.

### Supplemental Figure 10

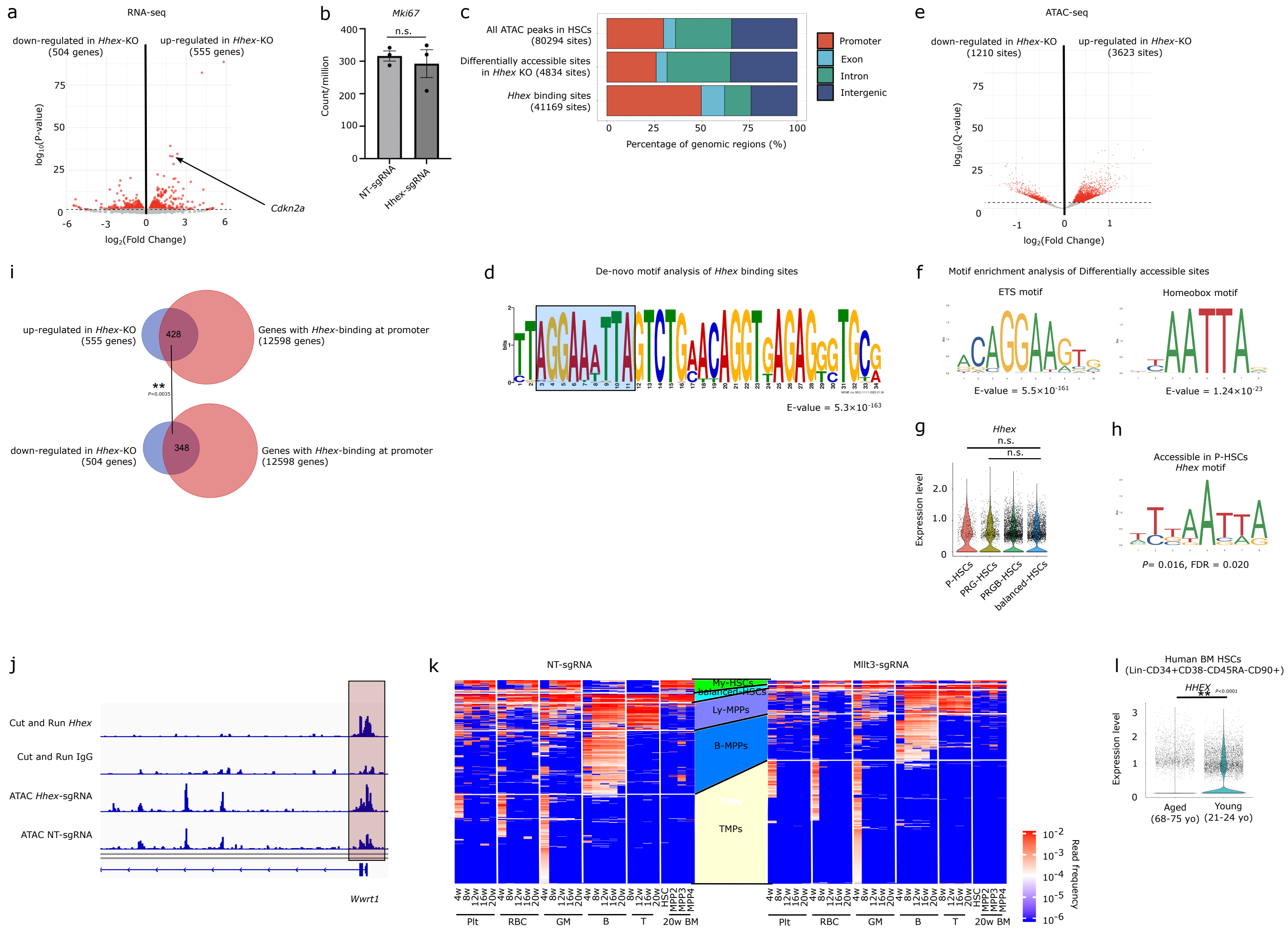

**Supplementary Figure 10 | Molecular analysis of *Hhex* KO.**

**a**, Volcano plot of differential gene expression between *Hhex*-sgRNA and NT-sgRNA groups. **b**, Dot plot showing the expression of *Mki67*. (**a,b,** 3 biological replicates for each group). **c**, Genomic distribution (promoters, exons, introns, and intergenic regions) of genome-wide ATAC-seq peaks detected in HSCs from NT-sgRNA control group (upper), ATAC-seq peaks showing significantly altered chromatin accessibility between *Hhex*-sgRNA and NT-sgRNA groups (middle) and *Hhex* Cut&Run peaks (bottom). **d**, De novo motif analysis of Hhex Cut&Run peaks. **e**, Volcano plot of differential chromatin accessibility between *Hhex*-sgRNA and NT-sgRNA groups. **f**, Motif enrichment analysis of differentially accessible ATAC-seq peaks. (**c-f,** ATAC-seq: 3 biological replicates per group; Cut&Run: 2 biological replicates). **g**, Violin plot of *Hhex* expression across HSC subpopulations. **h**, Motif analysis of ATAC-seq (Yu et al., 2023) showing increased accessibility of the *Hhex* binding motif in P-HSCs. **i**, Venn diagram shows genes upregulated upon *Hhex*-sgRNA perturbation that overlap with promoters bound by *Hhex* (upper) and genes downregulated upon *Hhex*-sgRNA perturbation that overlap with promoters bound by *Hhex* (bottom). **j**, Representative ATAC-seq peaks showing differential chromatin accessibility between *Hhex*-sgRNA and NT-sgRNA HSCs that also overlap *Hhex* Cut&Run peaks. **k**, Heat map showing clones derived from NT-sgRNA (left) and *Mllt3*-sgRNA (right). Each row represents a unique random DNA barcode. Each column represents time points (4w, 8w, 12w, 16w, and 20w after transplantation) and cell types (Plt, RBC, GM, B, T, ). Read frequency is represented by color-code scale. Each cluster was annotated by the output pattern (My-HSCs, balanced-HSCs, MyRPs, Ly-MPPs, B-MPPs, TMPs). **l**, Violin plots showing the expression levels of *HHEX* in human BM HSCs ($Lin^-CD34^+CD38^-CD45RA^-CD90^+$) in aged (68–75 years old; n = 3) and young (21–24 years old; n = 3) groups with adjusted *P* value displayed, based on data from Aksöz et al., 2024. *P<0.05; **P<0.01. **b,l,** *P* values calculated by Student t-test. **g,** *P* values were calculated by one-way or two-way ANOVA followed by Tukey's multiple comparison test. **i**, *P* values were calculated by Wilcoxon rank-sum test.

### Supplemental Figure 11

a

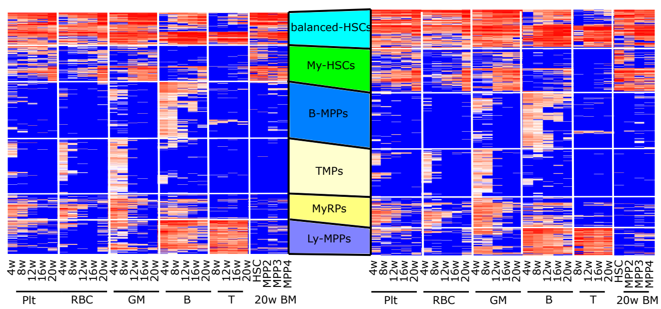


b

Proportion of each cell type (%)

*Bex4*-KO

| | NT | KO |
|---|---|---|
| My-HSCs | 15.1 | 17.9 |
| MyRPs | 8.8 | 10.3 |
| balanced-HSCs | 13.7 | 15.2 |
| Ly-MPPs | 14.1 | 11.3 |
| B-MPPs | 23.4 | 23.5 |
| TMPs | 24.9 | 21.8 |

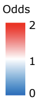


c

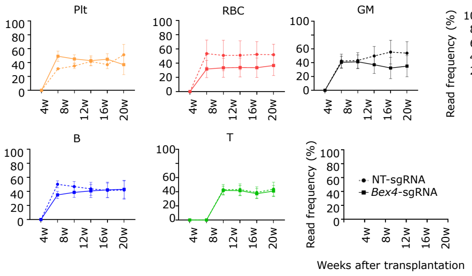


d

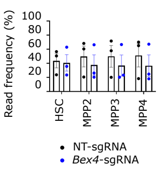

**Supplementary Data Figure 11 | Phenotype of Bex4 KO and OE.**

**a**, Heat map showing clones derived from NT-sgRNA (left) and *Bex4*-sgRNA (right). Each row represents a unique random DNA barcode. Each column represents time points (4w, 8w, 12w, 16w, and 20w after transplantation) and cell types (Plt, RBC, GM, B, T, HSC, MPP2, MPP3 and MPP4) (HSC:$CD150^{+}CD48^{-}Flt3^{-}Lin^{-}cKit^{+}Sca1^{+}$, MPP2:$CD150^{+}CD48^{+}Flt3^{-}Lin^{-}cKit^{+}Sca1^{+}$, MPP3:$CD150^{-}CD48^{+}Flt3^{-}Lin^{-}cKit^{+}Sca1^{+}$, MPP4:$Flt3^{+}Lin^{-}cKit^{+}Sca1^{+}$). Read frequency is represented by color-code scale. Each cluster was annotated by the output pattern (My-HSCs, balanced-HSCs, MyRPs, Ly-MPPs, B-MPPs, TMPs). **b**, Table showing the proportion of each cell type in NT-sgRNA and Bex4-sgRNA. **c**, Pseudo-bulk analysis of the read frequency of NT-sgRNA and Bex4-sgRNA in differentiated cells. **d**, Pseudo-bulk analysis of the read frequency of NT-sgRNA and *Bex4*-sgRNA in 20w BM (**C-F**, NT-sgRNA; n=1028, Bex4-sgRNA; n=1192 from 3 biological replicates from 2 independent experiments). *$P$<0.05; **$P$<0.01. **b**, $P$ values were calculated by Fisher's exact test. **c**, **d**, $P$ values were calculated by one-way or two-way ANOVA.
